# A systematic analysis of *Trypanosoma brucei* chromatin factors identifies novel protein interaction networks associated with sites of transcription initiation and termination

**DOI:** 10.1101/2021.02.09.430399

**Authors:** Desislava P. Staneva, Roberta Carloni, Tatsiana Auchynnikava, Pin Tong, Juri Rappsilber, A. Arockia Jeyaprakash, Keith R. Matthews, Robin C. Allshire

**Author notes:** MRC Human Genetics Unit, Institute of Genetics & Molecular Medicine, University of Edinburgh, Edinburgh EH4 2XU, Scotland, UK. These authors contributed equally to this work (Desislava P. Staneva, Roberta Carloni). Co-Corresponding authors: Robin Allshire, Keith Matthews.

## Abstract

Nucleosomes composed of histones are the fundamental units around which DNA is wrapped to form chromatin. Transcriptionally active euchromatin or repressive heterochromatin is regulated in part by the addition or removal of histone post-translational modifications (PTMs) by ‘writer’ and ‘eraser’ enzymes, respectively. Nucleosomal PTMs are recognised by a variety of ‘reader’ proteins which alter gene expression accordingly. The histone tails of the evolutionarily divergent eukaryotic parasite *Trypanosoma brucei* have atypical sequences and PTMs distinct from those often considered universally conserved. Here we identify 68 predicted readers, writers and erasers of histone acetylation and methylation encoded in the *T. brucei* genome and, by epitope tagging, systemically localize 63 of them in the parasite’s bloodstream form. ChIP-seq demonstrated that fifteen candidate proteins associate with regions of RNAPII transcription initiation. Eight other proteins exhibit a distinct distribution with specific peaks at a subset of RNAPII transcription termination regions marked by RNAPIII-transcribed tRNA and snRNA genes. Proteomic analyses identified distinct protein interaction networks comprising known chromatin regulators and novel trypanosome-specific components. Notably, several SET-domain and Bromo-domain protein networks suggest parallels to RNAPII promoter-associated complexes in conventional eukaryotes. Further, we identify likely components of TbSWR1 and TbNuA4 complexes whose enrichment coincides with the SWR1-C exchange substrate H2A.Z at RNAPII transcriptional start regions. The systematic approach employed provides detail of the composition and organization of the chromatin regulatory machinery in *Trypanosoma brucei* and establishes a route to explore divergence from eukaryotic norms in an evolutionarily ancient but experimentally accessible eukaryote.

## Introduction

Nucleosomes are composed of eight highly conserved core histone subunits (two each of H2A, H2B, H3 and H4) around which approximately 147 bp of DNA is wrapped. Nucleosomes are organised into chromatin fibres which provide the dynamic organisational platform underpinning eukaryotic gene expression regulation. Formation of transcriptionally active and silent chromatin states depends on the presence of DNA methylation (Suzuki and Bird, 2008), histone variants (Henikoff and Smith, 2015) and histone post-translational modifications (PTMs) (Bannister and Kouzarides, 2011). Repressive heterochromatin generally concentrates at the nuclear periphery, while active euchromatin localizes to the nuclear interior and can also associate with nuclear pores (Lemaître and Bickmore, 2015; Taddei et al., 2010). Molecular understanding of the composition and function of distinct chromatin types in nuclear architecture and gene expression regulation is most advanced in well-studied eukaryotic models (plants, yeasts, animals) (Allshire and Madhani, 2018). However, these represent only two eukaryotic supergroups while distinct early-branching lineages have highly divergent histones and chromatin-associated regulators. One particularly tractable model for early branching eukaryotes is *Trypanosoma brucei*, the causative agent of human sleeping sickness and livestock nagana in Africa, which has evolved separately from the main eukaryotic lineage for at least 500 million years. Reflecting their evolutionary divergence, detailed analyses of these parasites have revealed numerous examples of biomolecular novelty, including RNA editing of mitochondrial transcripts (Shapiro and Englund, 1995), polycistronic transcription of nuclear genes (Borst, 1986; Tschudi and Ullu, 1988) and segregation of chromosomes via an unconventional kinetochore apparatus comprising components distinct from other eukaryotic groups (Akiyoshi and Gull, 2014).

During its life cycle, *T. brucei* alternates between a mammalian host and the tsetse fly vector, a transition accompanied by extensive changes in gene expression leading to surface proteome alterations as well as metabolic reprogramming of the parasite (Matthews, 2005; Smith et al., 2017). In the mammalian host, bloodstream form (BF) parasites are covered by a dense surface coat made of variant surface glycoprotein (VSG). Only a single VSG gene is expressed from an archive consisting of ~2000 VSG genes and gene fragments (Horn, 2014). Periodically, *T. brucei* switches to express a new VSG protein to which no host antibodies have been produced, contributing to cyclical parasitaemia. Parasites taken up by the tsetse during blood meals differentiate in the fly midgut to the procyclic form (PF) which replaces all VSGs with procyclin surface proteins (Roditi and Liniger, 2002).

The *T. brucei* genome encodes four core histones (H2A, H2B H3, H4) and a variant for each core histone type (H2A.Z, H2B.V H3.V, H4.V), but it lacks a centromere-specific CENP-A/cenH3 variant (Akiyoshi and Gull, 2014). All trypanosome histones differ significantly in their amino acid sequence from their counterparts in conventional eukaryotes (Lowell and Cross, 2004; Lowell et al., 2005; Mandava et al., 2007; Thatcher and Gorovsky, 1994). For example, lysine 9 of histone H3, methylation of which specifies heterochromatin formation in many eukaryotes, is not conserved. Nonetheless, many histone PTMs have been detected in *T. brucei* and its relative *T. cruzi* (Janzen et al., 2006a; Kraus et al., 2020; Mandava et al., 2007), although only a handful of these have been characterized in some detail.

Unusually for a eukaryote, most trypanosome genes are transcribed in polycistronic units which are resolved by trans-splicing of a spliced leader (SL) RNA sequence to the 5′ end of the mRNA and polyadenylation at the 3′ end (Gunzl, 2010). RNAPII transcription usually initiates from broad (~10 kb) GT-rich divergent Transcriptional Start Regions (TSRs; comparable to promoters of other eukaryotes) enriched in nucleosomes containing the H2A.Z and H2B.V histone variants as well as the H3K4me and H4K10ac PTMs (Siegel et al., 2009; Wedel et al., 2017; Wright et al., 2010). Conversely, RNAPII transcription typically terminates at regions of convergent transcription known as Transcription Termination Regions (TTRs) that are marked by the presence of the DNA modification base J and the H3.V and H4.V variants (Schulz et al., 2016; Siegel et al., 2009). Less frequently, *T. brucei* transcription units are arranged head-to-tail requiring termination ahead of downstream TSRs. Termination between such transcription units is often coincident with genes transcribed by RNAPI or RNAPIII (Marchetti et al., 1998; Siegel et al., 2009).

The consensus view is that trypanosome gene expression is regulated predominantly post-transcriptionally via control of RNA stability and translation (Clayton, 2019). Nonetheless, numerous putative chromatin regulators can be identified as coding sequences in the *T. brucei* genome (Berriman et al., 2005), but their functional contexts are largely unexplored. Here we undertake comprehensive interrogation of bioinformatically identified putative readers, writers and erasers of histone acetyl and methyl marks encoded in the trypanosome genome. Our analysis included 41 predicted writers (HAT, DOT and SET domain proteins), 11 predicted erasers (HDAC and JmjC domain proteins) and 16 predicted readers of various domains. The cellular localization of each was determined in order to prioritise the analysis of those enriched in the nucleus. Subsequent chromatin immunoprecipitation and sequencing (ChIP-seq) showed that fifteen of the candidate proteins exhibit enrichment at RNAPII TSRs. In contrast, eight proteins were found to be specifically enriched at TTRs. Finally, comprehensive proteomics analysis of the chromatin-associated factors showed that they form distinct protein interaction networks. In particular, we find that two SET domain proteins associate with putative readers with which they occupy RNAPII TSRs. Moreover, six of the seven Bromo domain proteins are involved in four interaction networks enriched at TSRs while the BDF7 network alone marks TTRs. The association of SET and Bromo domain proteins with conserved RNAPII subunits, HATs, chromatin chaperones and remodelling factors suggests that they play pivotal roles in defining sites of transcription initiation and termination. In combination, these analyses provide a comprehensive atlas of the chromatin regulatory machinery associated with equivalents of promoter and transcription termination regions in these evolutionarily divergent eukaryotes.

## Results and Discussion

### Identification of putative chromatin regulators

We identified 68 putative regulators or interpreters of histone lysine acetylation and methylation by interrogating the trypanosome genome database (TriTrypDB) and using additional homology-based searches with known chromatin writer, reader and eraser domains (Table 1; Supplemental Fig. S1A; Supplemental Table S1). These approaches allowed us to detect 16 potential readers with the following domains: Bromo (Haynes et al., 1992), PHD (Aasland et al., 1995), Tudor (Ponting, 1997), Chromo (Paro, 1990; Singh et al., 1991), PWWP (Stec et al., 2000) and Znf-CW (Perry and Zhao, 2003). With respect to writers of histone modifications, we found 9 potential histone acetyltransferases (HATs) belonging to the MYST, LPLAT and GNAT families plus the EAF6 component of the NuA4 HAT complex (Hishikawa et al., 2008; Roth et al., 2001), 29 SET domain (Dillon et al., 2005) and 3 DOT domain (Feng et al., 2002) putative histone methyltransferases (HMTs). Our searches for potential erasers identified 7 histone deacetylases (class I, class II and Sir2-related HDACs) (Grozinger and Schreiber, 2002) and 4 JmjC domain demethylases (Klose et al., 2006). The function of some of these putative chromatin regulators has been explored previously (Figueiredo et al., 2009; Maree and Patterton, 2014; Supplemental Table S2).

**Table 1.**
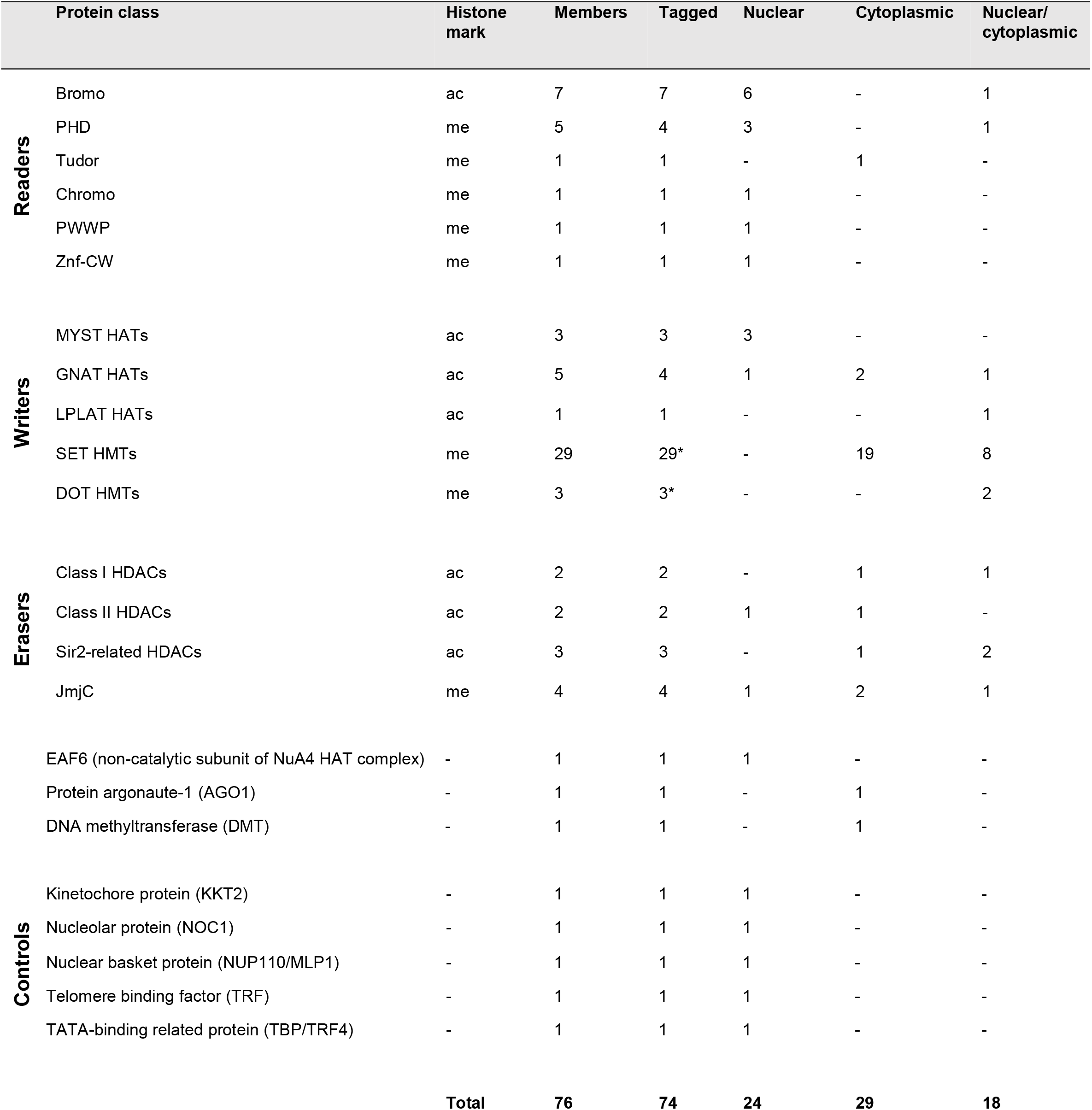
Summary of proteins N-terminally tagged with YFP. * Correct integration of the SET12, SET30 and DOT1 tagging constructs was confirmed by PCR but the tagged proteins were not detected by western analysis. Therefore, we do not report localization for these proteins.

To examine these putative writers, readers and erasers of histone PTMs, each was YFP-tagged and localized in bloodstream form parasites. AGO1 (Shi et al., 2009) and the putative DNA methyltransferase DMT (Militello et al., 2008) were also included in our analysis since they are associated with chromatin-based silencing in other eukaryotes (Allshire and Madhani, 2018; Kloc and Martienssen, 2008; Suzuki and Bird, 2008).

Additionally, we examined several control proteins with known distinctive nuclear distributions; these were the trypanosome kinetochore protein KKT2 (Akiyoshi and Gull, 2014), the nucleolar protein NOC1 (Alsford and Horn, 2012), the nuclear pore basket protein NUP110/MLP1 (DeGrasse et al., 2009) as well as the telomere repeat-binding factor TRF (Li et al., 2005) and the TATA-binding related protein TBP/TRF4 (Ruan et al., 2004).

### Cellular localization of the putative chromatin regulators

Candidate proteins were endogenously tagged with eYFP in bloodstream form parasites of the *T. brucei* Lister 427 strain. Proteins were tagged N-terminally to avoid interference with 3′ UTR sequences involved in mRNA stability control (Clayton, 2019). Of the 76 selected proteins (including controls), 74 were successfully tagged while cells expressing correctly tagged PHD3 and HAT8 were not obtained. The tagging constructs for SET12, SET30 and DOT1 were correctly integrated but YFP-tagged proteins were not detectable by western analysis suggesting tag failure, low protein abundance or no expression in bloodstream form parasites.

Immunolocalization with anti-GFP antibodies was used to identify proteins residing in the nucleus which might associate with chromatin. The control proteins exhibited localization patterns expected for telomeres (TRF – nuclear foci), TATA-binding protein (TBP – nuclear foci), kinetochores (KKT2 – nuclear foci), nucleolus (NOC1 – single nuclear compartment) and nuclear pores (NUP110 – nuclear rim) (Supplemental Fig. S2). Some cytoplasmic signal was also detected for these proteins but this likely corresponds to background as evidenced by the signal observed in untagged control cells (Fig. 1; Supplemental Fig. S2)..

**Figure 1.**
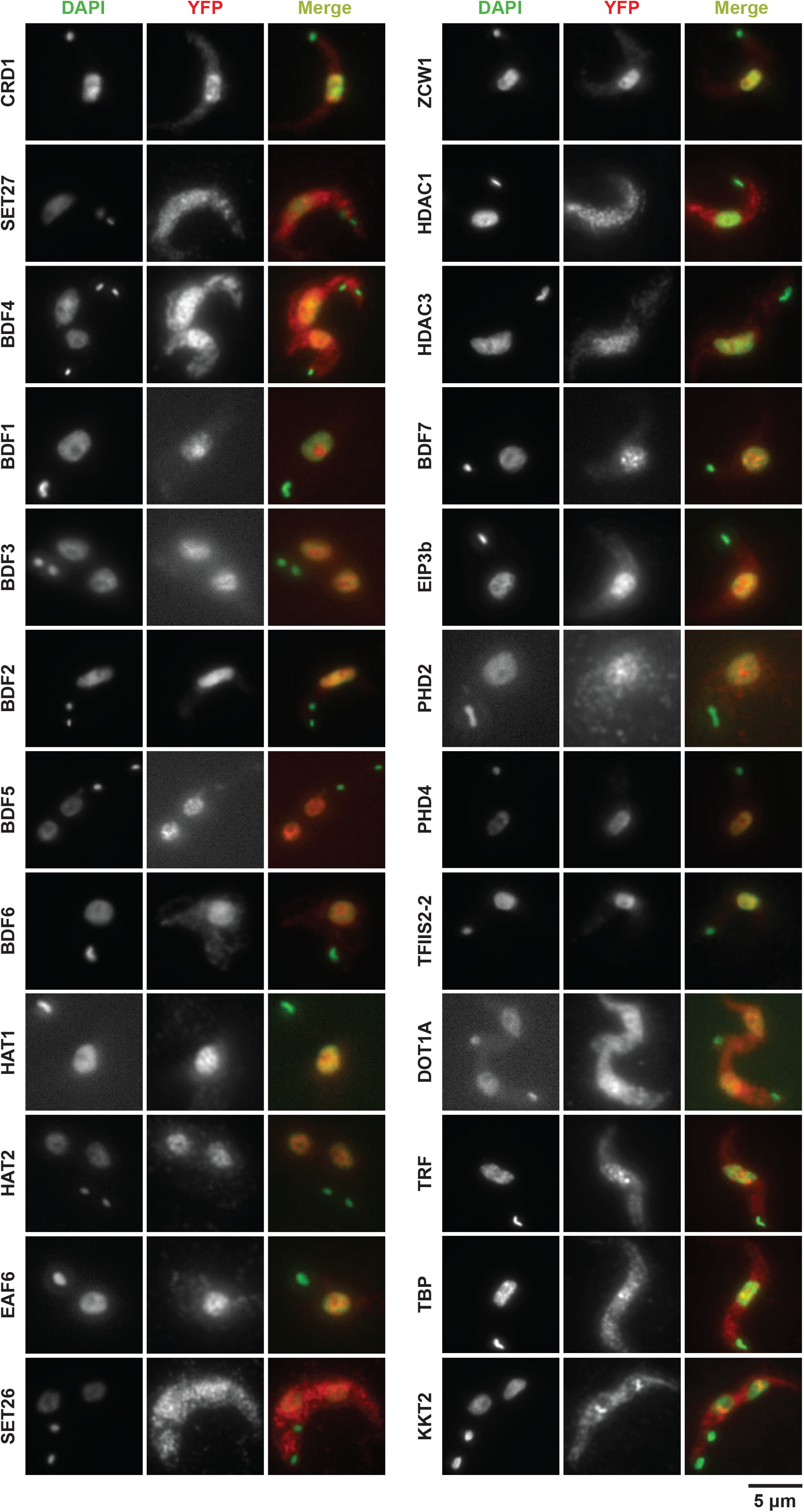
Cellular localization of chromatin-associated *T. brucei* candidate and control proteins. The indicated YFP-tagged proteins expressed in bloodstream Lister 427 cells from their endogenous genomic loci were detected with an anti-GFP primary antibody and an Alexa Fluor 568 labelled secondary antibody. Nuclear and kinetoplast (mitochondrial) DNA were stained with DAPI. Representative images are shown for those proteins that gave a specific ChIP-seq signal and are ordered according to ChIP-seq patterns shown in Figure 1A and Figure 5A. The kinetochore protein KKT2 is included as a positive control. Images for all other tagged proteins and the untagged control are shown in Figure S2. Bar = 5 µm.

Of the YFP-tagged candidate proteins, 19 were exclusively nuclear, 29 exhibited only a cytoplasmic localization and 18 were found in both compartments (Table 1; Supplemental Table S1). Twenty-four candidate and control proteins with exclusive or some nuclear localization were subsequently found to associate with RNAPII transcription initiation and termination regions (see below; Fig. 1), while the remaining proteins that were not detected on chromatin displayed all three localization patterns (Supplemental Figure S2). Consistent with previous observations, HAT1-to-3 decorated nuclear substructures (Kawahara et al., 2008), as did the predicted EAF6 subunit of the NuA4 HAT complex. In contrast, HAT5 gave both nuclear and cytoplasmic signals whereas both HAT6 and HAT7 localized to the cytoplasm. Surprisingly, of the 29 identified putative SET-domain methyltransferases, only eight exhibited some nuclear localization, while most were concentrated in the cytoplasm. Examination of cells expressing YFP-tagged predicted reader proteins revealed that BDF1- to-3, BDF5-to-7, PHD1, PHD2, PHD4, CRD1, TFIIS2-2 and ZCW1 were exclusively nuclear, whereas BDF4 and PHD5 displayed both nuclear and cytoplasmic localization and the sole Tudor domain protein TDR1 was cytoplasmic. In agreement with previous analyses (Wang et al., 2010), HDAC3 was exclusively nuclear whereas HDAC1 was nuclear/cytoplasmic, and both HDAC2 and HDAC4 resided in the cytoplasm. The Sir2-related proteins Sir2rp1 and Sir2rp3 localized to the nucleus and cytoplasm whereas Sir2rp2 was detected only in the cytoplasm. Of the four identified demethylases, JMJ2 was nuclear, JMJ1 and CLD1 were cytoplasmic and LCM1 was found in both compartments. Both AGO1 and DMT were exclusively cytoplasmic suggesting that they are unlikely to be involved in directing chromatin or DNA modifications (Supplemental Fig. S2).

Of the 66 candidate proteins that were successfully YFP-tagged, expressed and localized in bloodstream form cells, 46 have also been also tagged in the TrypTag project in procyclic form cells (Dean et al., 2017; Table S1). For most proteins, our results are in agreement with the TrypTag data and any discrepancies may be indicative of differences in protein localization between the different developmental forms of *T. brucei*.

### Many putative chromatin regulators accumulate at RNAPII TSRs

We performed ChIP-seq for all expressed candidate and control proteins except TDR1 and NOC1 (69 in total) to determine which proteins associate with chromatin and assess their distribution across the *T. brucei* genome. Our rationale was that ChIP-seq might register chromatin association even if cellular localization analysis reported a protein to be predominantly cytoplasmic. As expected, the kinetochore control protein KKT2 was specifically enriched over centromeric regions (Fig. 2A). Consistent with their cytoplasmic localizations, AGO1 and DMT registered no ChIP-seq signal (not shown). Moreover, under our standard fixation and ChIP-seq conditions, no enrichment over any genomic region was detected for 45 of the YFP-tagged proteins, including several that exhibited exclusive nuclear localization (Supplemental Table S3).

**Figure 2.**
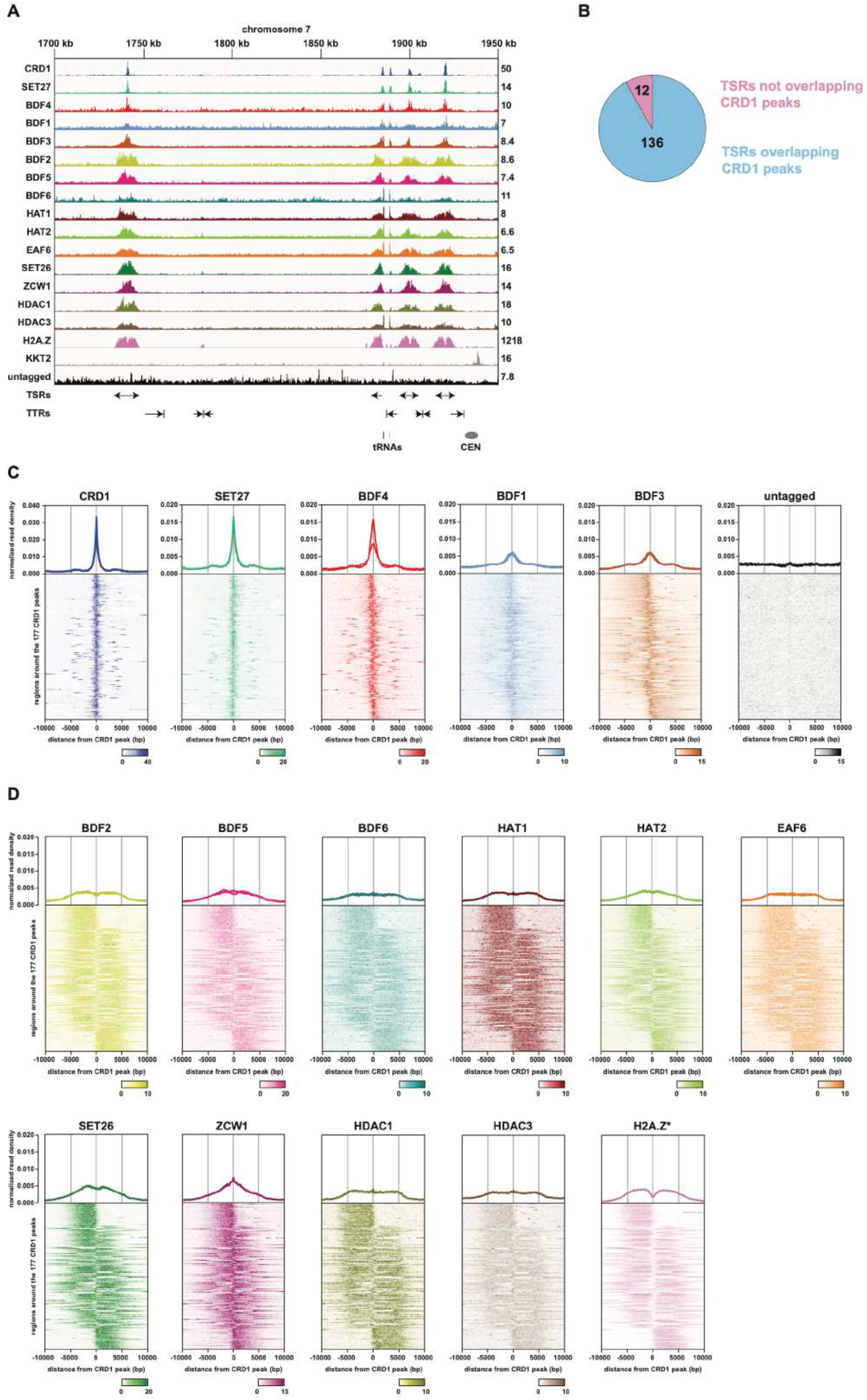
ChIP-seq reveals two classes of proteins at *T. brucei* transcription start regions. **A**. A region of chromosome 7 (coordinates as indicated, kb) is shown with ChIP-seq reads mapped for the indicated proteins. Tracks are scaled separately as reads per million (values shown to right of tracks). The kinetochore protein KKT2 is included as positive control and is enriched at centromeric regions. ChIP-seq performed in Lister 427 cells expressing no tagged protein (untagged) provides a negative control. Protein ChIP-seq profiles are ordered according to their patterns. Previous H2A.Z ChIP-seq data (Wedel et al. 2017) allowed Transcription Start Regions (TSRs) to be identified. No input data was available for normalization of H2A.Z reads resulting in a higher read scale. The position of bidirectional/divergent and unidirectional/single TSRs is indicated with arrows showing the direction of transcription. The position of convergent and single Trancription Termination Regions (TTRs) is shown with arrows indicating the direction from which transcription is halted. The position of tRNA genes (blue bars) and the chromosome 7 centromere (CEN, grey oval) are marked. Positions of the various genomic elements (TSRs, TTRs, tRNAs, CEN) were obtained from the Lister 427 genome annotation (Muller et al, 2018). **B**. Most TSRs annotated in the Lister 427 genome overlap with YFP-CRD1 ChIP-seq peaks. **C**. Enrichment profiles of Class I TSR factors. The metagene plots (top) show normalized read density around all CRD1 peak summits, with individual replicates for each protein shown separately. Note the different scale for CRD1. The heatmaps (bottom) are an average of all replicates for each protein and show protein density around individual CRD1 peaks. Scale bars represent reads that were normalized to input and library size. **D**. As in (C) for Class II TSR factors.

In total, 24 of the YFP-tagged proteins assayed (including the KKT2 control) gave specific enrichment patterns across the *T. brucei* genome. Fifteen of our candidate proteins displayed enrichment that was adjacent to, or overlapping with, the previously reported H2A.Z peaks indicating that these proteins are enriched at known RNAPII TSRs (Fig. 2A; Supplemental Fig. S3A). These TSR-associated proteins were BDF1-to-6, CRD1, EAF6, HAT1, HAT2, HDAC1, HDAC3, SET26, SET27, and ZCW1. The fact that six Bromo domain proteins, three histone acetyltransferases and a NuA4 component are included in this set is consistent with them acting to promote RNAPII-mediated transcription, a hallmark of which is histone acetylation (Roth et al., 2001). Indeed, BDF1, BDF3 and BDF4 have previously been shown to be enriched at sites of *T. brucei* RNAPII transcription initiation where nucleosomes exhibit HAT1-mediated acetylation of H2A.Z and H2B.V and HAT2-mediated acetylation of histone H4, which are important for normal RNAPII transcription from these regions (Kraus et al., 2020; Schulz et al., 2015; Siegel et al., 2009). The remaining 10 proteins we identified as being associated with TSRs have not been previously shown to act at kinetoplastid RNAPII TSRs.

### Putative chromatin regulators exhibit two distinct TSR association patterns

In yeast, where many RNAPII genes are transcriptionally regulated, a group of chromatin regulators are enriched specifically at promoters where they assist in transcription initiation while others travel with RNAPII into gene bodies aiding transcriptional elongation, splicing and termination (Carrozza et al., 2005; Cheung et al., 2008; Jonkers and Lis, 2015; Joshi and Struhl, 2005; Kaplan et al., 2003; Keogh et al., 2005; Li et al., 2007; Mason and Struhl, 2003; Venkatesh and Workman, 2013). We therefore compared the enrichment profiles of the TSR-associated proteins relative to that of CRD1 (<u>C</u>h<u>r</u>omo<u>d</u>omain protein 1) which displayed the sharpest and highest peak signal. We identified 177 CRD1 peaks across the *T. brucei* Lister 427 genome which overlap with 136 of the 148 annotated RNAPII TSRs (Figure 2B). For each TSR-associated factor, normalized reads were assigned to 10 kb windows upstream and downstream of all CRD1 peak summits. The general peak profile for each protein was then displayed as a metagene plot and the read distribution around individual CRD1 peaks represented as a heatmap (Fig. 2C,D; Supplemental Fig. S4). This analysis indicated that CRD1, SET27, BDF4 and, to a lesser extent, BDF1 and BDF3 exhibit sharp peaks at all RNAPII TSRs. We refer to these as Class I TSR-associated factors (Fig. 2C). The remaining 10 proteins were more broadly enriched over the same regions with a slight trough evident in the signal for at least five (BDF2, BDF5, HAT1, HAT2, SET26), perhaps indicative of two overlapping peaks as observed for H2A.Z (Fig. 2D). We refer to these as Class II TSR-associated factors. Class II factor profiles are similar to the previously reported H2A.Z enrichment pattern (Siegel et al., 2009), with the signal gradually declining over 5-10 kb regions on either side of the CRD1 peak summit. We observed similar peak widths of Class II factors across all TSRs regardless of polycistron length, thus there is no apparent relationship between TSR size and the length of the downstream polycistronic transcription unit.

In *T. brucei*, most RNAPII promoters are bidirectional, initiating production of stable transcripts in both directions, but unidirectional promoters are also present. To investigate the relationship between transcription directionality and enrichment of our candidate proteins, we sorted the heatmaps of Class II factors by their distribution around CRD1 peaks. We then compared the sorted heatmaps with previously published RNA-seq data from which the direction of RNAPII transcription was derived (Naguleswaran et al., 2018). This analysis demonstrated that Class II TSR-associated factors exhibit specific enrichment in the same direction as RNAPII transcription initiated from uni- or bi-directional promoters (Fig. 3).

**Figure 3.**
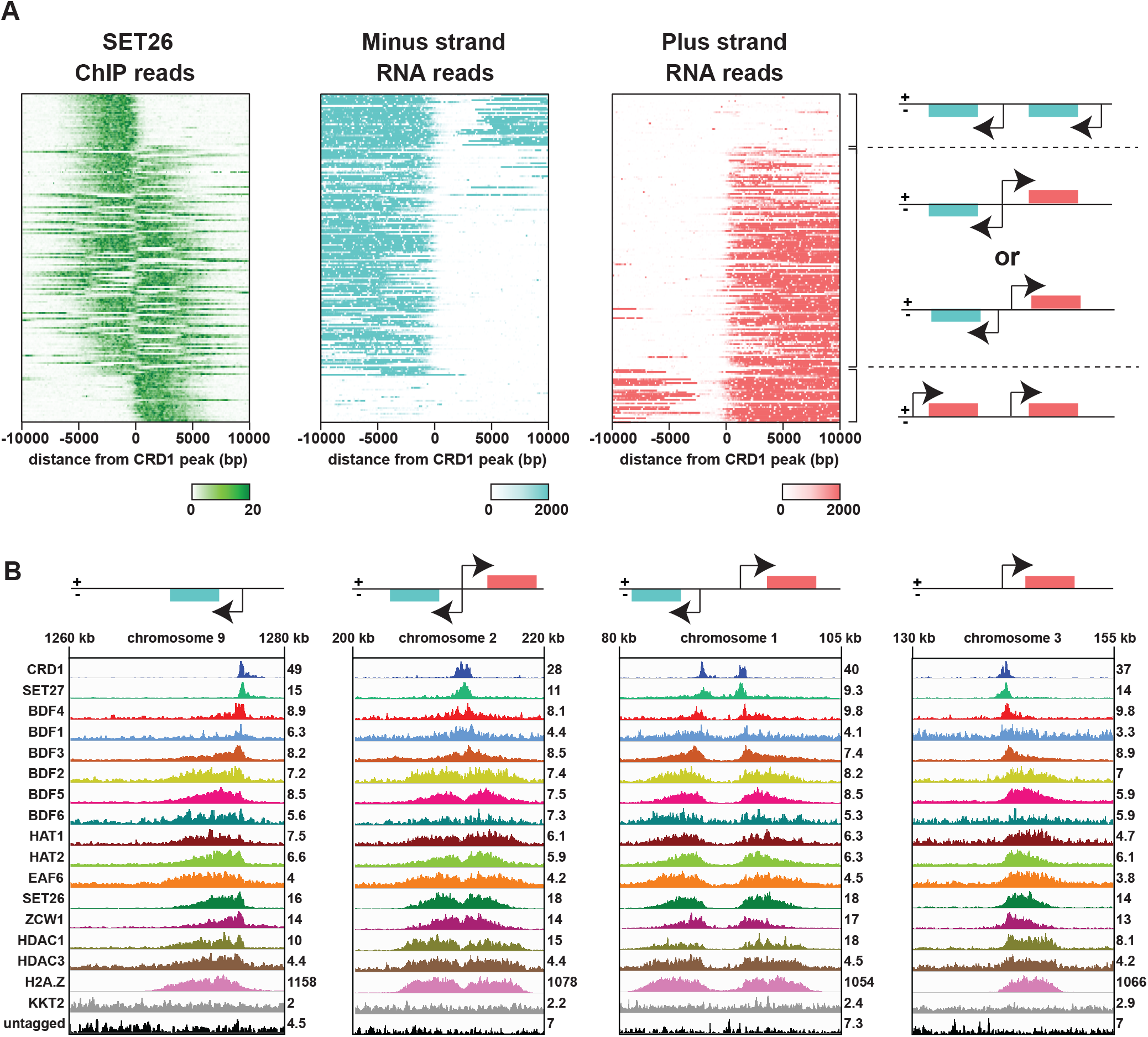
Enrichment of Class II proteins at RNAPII TSRs follows the direction of RNAPII transcription. A. SET26 is used as a representative protein of Class II TSR-associated factors. Comparison of SET26 ChIP-seq data with strand specific RNA-seq data (Naguleswaran et al. 2018) shows that SET26 reads are enriched in the same direction as RNAPII transcript reads. Heatmaps show from top to bottom: minus strand reads from unidirectional TSRs (top), plus and minus strand reads from bidirectional TSRs or from two close unidirectional TSRs (middle), and plus strand reads from unidirectional TSRs (bottom). B. Examples from the different heatmap regions described in (A). Tracks are scaled separately as reads per million (values shown to right of tracks).

### Proteins associated with TSRs participate in discrete interaction networks

The analyses presented above suggest that, as in yeast, proteins which exhibit either narrow (Class I) or broad (Class II) association patterns across RNAPII promoter regions might play different roles such as defining sites of RNAPII transcriptional initiation or facilitating RNAPII processivity through their association with chromatin and interactions with RNAPII auxiliary factors. Determining how these various activities are integrated through association networks should provide insight into how the distinct sets of proteins might influence RNAPII transcription. Therefore, we affinity selected each tagged protein that registered specific ChIP-seq signals at TSRs or TTRs and identified their protein interaction networks by mass spectrometry.

Below we first detail the interaction networks for TSR-associated proteins (Fig. 4; Supplemental Table S5), the homologies for key interacting proteins identified through HHpred searches (Supplemental Table S6) and discuss their possible functional implications. Interacting partners identified by proteomics are also included for nine proteins (PHD1, HAT3, AGO1, NUP110, SET13, SET15, SET20, SET23 and SET25) for which no specific ChIP-seq signal was obtained (Supplemental Fig. S5).

**Figure 4.**
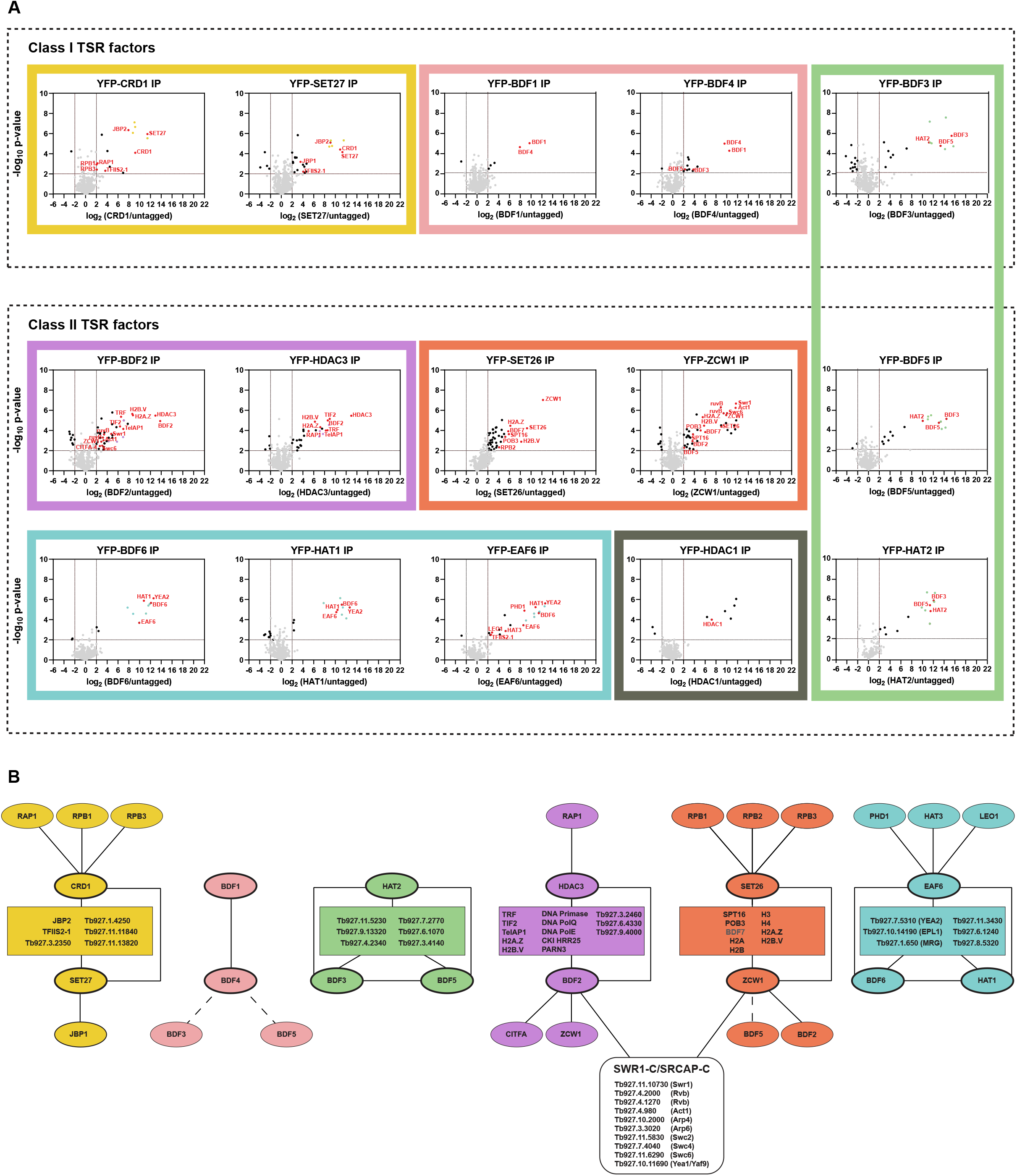
Class I and Class II TSR-associated factors define distinct interaction networks. **A**. YFP-tagged proteins found to be enriched at TSRs were analyses by LC-MS/MS to identify their protein interactions. The data for each plot is based on three biological replicates. Cut-offs used for significance: log_2_ (tagged/untagged) > 2 or < −2 and p < 0.01 (Student’s t-test). Enrichment scores for proteins identified in each affinity selection are presented in Table S5. Significantly enriched proteins are indicated by black or coloured dots. Proteins of interest are indicated by red font. Plots in the same box show reciprocal interactions. Uncharacterized proteins common to several affinity selections are depicted in yellow, green, purple and cyan. **B**. Key proteins identified as being associated with the indicated YFP-tagged bait proteins (thick oval outlines). Rectangles contain proteins common to several affinity purifications. Lines denote associations between proteins. The interactions of BDF3 and BDF5 with BDF4 (dashed lines); BDF5 with ZCW1 (dashed lines); and BDF7 (grey) with ZCW1 and SET26 were not confirmed by reciprocal affinity selections.

#### Class I: CRD1, SET27, BDF4, BDF1 and BDF3

##### CRD1 and SET27

Affinity selection of YFP-CRD1 and YFP-SET27 (Fig. 4; Supplemental Table S5) revealed that they both associate strongly with each other, with four uncharacterized proteins (Tb927.1.4250, Tb927.3.2350 Tb927.11.11840, Tb927.11.13820) and with JBP2. JBP2 is a TET-related hydroxylase that catalyses thymidine oxidation on route to the synthesis of the DNA modification base J which is found at transcription termination regions and telomeres in trypanosomes (Cliffe et al., 2010; Reynolds et al., 2016; Schulz et al., 2016). In addition, SET27 associates with JBP1 -another TET-related thymidine hydroxylase involved in base J synthesis (Borst and Sabatini, 2008). The chromodomain of CRD1 exhibits marginal similarity when aligned with other chromodomain proteins (Supplemental Fig. S1B), however its reciprocal association with SET27 suggests that they might function together as a reader-writer pair at TSRs. In yeast and human cells, the Set1/SETD1 methyltransferase installs H3K4 methylation at promoters (Shilatifard, 2012), thus *T. brucei* SET27 might play a similar role at TSRs. Recently, methylation of histones on H3K4, H4K10, H4K2 and H2AZ K45, K49, K51, K164 and K129 was found to be prevalent at TSRs (Kraus et al., 2020). SET27 likely catalyses the methylation of at least one of these lysine residues which may then be bound by CRD1, ensuring SET27 recruitment and persistence of the methylation event(s) that it installs on histones within resident TSR nucleosomes. The association of the RPB1 and RPB3 RNAPII subunits with CRD1 underscores its likely involvement in linking such chromatin modifications with transcription.

##### BDF1 and BDF4

Consistent with their colocalization in Class I ChIP-seq peaks, YFP-BDF1 and YFP-BDF4 showed strong reciprocal association with each other (Figure 4; Table S5). BDF4 also exhibited weak association with BDF3 and the Class II factor BDF5, suggesting that BDF3, which exhibits a broader peak pattern (Fig. 2), may straddle the interface between both classes of factors at RNAPII TSRs. Bromo domains are known to bind acetylated histones (Zaware and Zhou, 2019), presumably they are attracted to TSRs due to the presence of highly acetylated histones, particularly H2A.Z, H2B.V and H4, in resident nucleosomes (Kraus et al., 2020). The coincidence of the H2A.Z histone variant along with such active histone modifications in evolutionarily distinct eukaryotes suggest that they act together to recruit various chromatin remodelling and modification activities to ensure efficient transcription.

##### BDF3 (Class I), BDF5 (Class II) and HAT2 (Class II)

BDF3, BDF5 and HAT2 reciprocally associate with each other and a set of six uncharacterized proteins (Tb927.3.4140, Tb927.4.2340, Tb927.6.1070, Tb927.7.2770, Tb927.9.13320, Tb927.11.5230) suggesting that these nine proteins may act together in a complex (Figure 4; Table S5). HAT2 mediates acetylation of histone H4 and promotes normal transcriptional initiation by RNAPII (Kraus et al., 2020). The bromodomains of BDF3 and BDF5 may guide HAT2 to pre-existing acetylation at TSRs to maintain the required acetylated state for efficient transcription. We also note that Tb927.3.4140 displays similarity to Poly ADP Ribose Polymerase (PARP; Table S6), ribosylation might contribute to TSR definition by promoting chromatin decompaction as seen upon *Drosophila* heat shock puff induction (Sawatsubashi et al., 2004; Tulin et al., 2003; Tulin and Spradling, 2003) and at some mammalian enhancers-promoter regions (Benabdallah et al., 2019). Tb927.11.5230 contains an EMSY ENT domain whose structure has been determined (Mi et al., 2018; Table S6); such domains are present in several chromatin regulators. Interestingly, Tb927.4.2340 exhibits similarity to the C-terminal region of the vertebrate TFIID TAF1 subunit (Supplemental Table S6). Metazoan TAF1 bears two Bromo domains in its C-terminal region whereas in yeast the double Bromo domain component of TFIID is contributed by the separate Bdf1 (or Bdf2) proteins (Matangkasombut et al., 2000; Timmers, 2020). *T. brucei*, BDF5 contains two Bromo domains and may thus be equivalent to the yeast TFIID BDF1 subunit.

#### Class II: BDF2, BDF5, BDF6, EAF6, HAT1, HAT2, HDAC1, HDAC3, SET26, ZCW1

##### BDF2 and HDAC3

We found that both TSR-enriched histone variants H2A.Z and H2B.V strongly associate with BDF2 and HDAC3, which interact with each other as well as with four uncharacterized proteins (Tb927.3.2460, Tb927.6.4330, Tb927.9.4000, Tb927.9.8520; Figure 4; Table S5). We note that Tb927.9.8520 exhibits similarity to DNA Polymerase Epsilon (DNAPolE; Table S6) and both DNA Polymerase Theta (DNAPolQ) and DNA Primase were also enriched along with the PARN3 poly(A) specific ribonuclease and Casein Kinase I (CKI). Moreover, Tb927.3.2460 exhibits similarity to a nuclear pore protein while Tb927.6.4330, Tb927.9.4000 and DNAPolQ were previously shown to associate with the telomere binding protein TRF and DNA Primase is known to interact with TelAP1 (Reis et al., 2018). Here we find that the telomere-associated proteins TRF, TIF2, TelAP1 and RAP1 are also enriched in both BDF2 and HDAC3 affinity selections. The significance of this association is unknown, however HDAC3 has previously been shown to be required for telomeric VSG expression site silencing (Wang et al., 2010) and RNAi knock-down of Tb927.6.4330 causes defects in telomere-exclusive VSG gene expression (Glover et al., 2016). Given that actively transcribed regions tend to be hyperacetylated (Kouzarides, 2007) it was surprising to observe HDAC3 enrichment at RNAPII TSRs. HDAC3 may be required to reverse acetylation associated with newly deposited histones during S phase (Stewart-Morgan et al., 2020) or to remove acetylation added during expected H2A-H2B:H2A.Z-H2B.V dimer-dimer exchange events at TSR regions (Bonisch and Hake, 2012; Millar et al., 2006).

##### SET26 and ZCW1

SET26 and ZCW1 associated with each other as well as with most histones and histone variants (Fig. 4; Supplemental Table S5). In addition, both SET26 and ZCW1 showed strong interaction with the SPT16 and POB3 subunits of the FACT (Facilitates Chromatin Transcription) complex which aids transcriptional elongation in other eukaryotes (Belotserkovskaya et al., 2003). SET26 could perform an analogous role to yeast Set2 H3K36 histone methyltransferase which travels with RNAPII and, together with FACT, ensures that chromatin integrity is restored behind advancing RNAPII. These activities are known to prevent promiscuous transcriptional initiation events from cryptic promoters within open reading frames (Carrozza et al., 2005; Cheung et al., 2008; Joshi and Struhl, 2005; Kaplan et al., 2003; Keogh et al., 2005; Li et al., 2007; Mason and Struhl, 2003; Venkatesh and Workman, 2013). Interestingly, ZCW1 also exhibits strong association with apparent orthologs of several SWR1/SRCAP/EP400 remodelling complex subunits (Scacchetti and Becker, 2020; Willhoft and Wigley, 2020), including Swr1/SRCAP (Tb927.11.10730), Swc6/ZNHIT1 (Tb927.11.6290), Swc2/YL1 (Tb927.11.5830), the RuvB-related helicases (Tb927.4.2000; Tb927.4.1270), actin and actin-related proteins (Tb927.4.980, Tb927.10.2000, Tb927.3.3020), and the possible equivalents of the Swc4/DMAP1 (Tb927.7.4040) and Yaf9/GAS41 YEATS domain protein (Tb927.10.11690, designated YEA1) subunits (Table S6; Table S7). Most of these putative TbSWR1-C subunits were also detected as being associated with BDF2 (Fig. 4; Supplemental Table S5, S6). Thus, since the yeast Bdf1 and human BRD8 Bromodomain proteins also associate with SWR1-C/SRCAP-C/EP400, it is likely that TbBDF2 forms a similar function in engaging acetylated histones. The SWR/SRCAP remodelling complexes are well known for being required to direct the replacement of H2A with H2A.Z in nucleosomes residing close to transcriptional start sites (Mizuguchi et al., 2004; Ruhl et al., 2006). The prevalence of both H2A.Z and H2B.V with affinity selected ZCW1 suggests that it may also play a role in directing the *T. brucei* SWR/SRCAP complex to TSRs to ensure incorporation of H2A.Z-H2B.V in place of H2A-H2B in resident nucleosomes.

##### BDF6, EAF6 and HAT1

BDF6, EAF6 and HAT1 exhibit robust association with each other, with a second YEATS domain protein - the possible ortholog of acetylated-H2A.Z binding Yaf9/Gas41 (Tb927.7.5310, designated YEA2) and a set of five other uncharacterized proteins (Tb927.1.650, Tb927.6.1240, Tb927.8.5320, Tb927.10.14190, Tb927.11.3430). Tb927.1.650 is an MRG domain protein with similarity to yeast Eaf3 while Tb927.10.14190 appears to be orthologous to yeast Epl1, both of which along with Yaf9 are components of the yeast NuA4 complex (Supplemental Table S6, S8). The yeast Yaf9 YEATS protein contributes to both the NuA4 HAT and SWR1 complexes and can bind acetylated or crotonylated histone tails (Arrowsmith and Schapira, 2019; Timmers, 2020). In *T. brucei*, it appears that distinct YEATS domain proteins contribute to putative SWR1 (YEA1) and NuA4 (YEA2) complexes. HAT1 associates with both BDF6 and EAF6 whereas HAT3, EAF6 and PHD1 reciprocally associate with each other and share Tb927.11.7880, a YNG2-related ING domain protein, as a common interactor (Supplemental Figure S5; Table S6, S8). Thus, HAT3-EAF6-PHD1-YNG2 perhaps represents a *T. brucei* subcomplex analogous to yeast piccolo-NuA4 while HAT1-EAF6-BDF6-EAF3-EPL1-YEA2 may form the larger NuA4 complex (Doyon and Cote, 2004; Wang et al., 2018). In *T. brucei*, Esa1/TIP60 catalytic MYST acetyltransferase function may be shared between HAT1 and HAT3. Surprisingly, however, although we have clearly identified NuA4-like complexes, no association was detected with a *T. brucei* Eaf1/EP400-related Helicase-SANT domain protein that provides the platform for the assembly of distinct modules of the yeast and metazoan NuA4 complexes (Levi et al., 1987; Scacchetti and Becker, 2020).

Together, BDF6, EAF6, and HAT1 appear part of a putative NuA4-related complex that functions at *T. brucei* TSRs. HAT1 has been shown to be required for H2A.Z and H2B.V acetylation and efficient RNAPII engagement and transcription (Kraus et al., 2020). BDF6, YEA2 or both may bind acetylated histones at TSRs to promote stable association of interacting chromatin modification and remodelling activities that enable the required histone dynamics to take place in these highly specialized regions and thereby facilitate efficient transcription.

##### HDAC1

We also readily detect HDAC1 enriched across a broad region at RNAPII TSRs, but notably it does not associate with any of the other factors that are enriched over these regions. However, five uncharacterized proteins (Tb927.3.890, Tb927.4.3730, Tb927.6.3170, Tb927.7.1650, Tb927.9.2070) reproducibly associated with HDAC1. HHpred searches detected some similarity of Tb927.3.890, Tb927.4.3730 and Tb927.6.3170 to chromatin-associated proteins (Supplemental Table S6) and Tb927.9.2070 was also enriched with CRD1. HDAC1 was previously shown to be an essential nuclear protein whose knock down increases silencing of telomere-adjacent reporters in bloodstream form parasites (Wang et al., 2010). However, a general role for HDAC1 at RNAPII TSRs was not anticipated. HDAC1 has been reported to be mainly cytoplasmic in procyclic cells (Wang et al., 2010) and it is therefore expected to be absent from RNAPII TSRs in insect form parasites.

### Eight proteins are specifically enriched over RNAPII TTRs coincident with RNAPIII genes

Our analysis of ChIP-seq association patterns also revealed a distinct set of eight proteins which displayed specific enrichment over a subset of RNAPII TTRs. Six of the selected candidate reader and writer proteins (BDF7, ELP3b, PHD2, PHD4, TFIIS2-2 and DOT1A) displayed sharp peaks at locations distinct from known TSRs (Fig. 5A; Supplemental Figure S3B). Unexpectedly, two of our selected control proteins (TRF and TBP) were also found at the same locations. All eight proteins were enriched at the U2 (Tb927.2.5680) and U6 (Tb927.4.1213) <u>s</u>mall <u>n</u>uclear snRNA genes and at 53 of the 69 annotated transfer tRNA genes in the current assembly of the *T. brucei* 427 genome (Fig. 5A,B; Supplemental Table S4). We observed enrichment of only some of these proteins at 11 additional tRNA genes and no association with the remaining 5 tRNAs. *T. brucei* snRNAs and tRNAs are known to be transcribed by RNAPIII (Nakaar et al., 1997; Tschudi and Ullu, 2002). Additionally, TBP was enriched over the RNAPIII-transcribed 5S rRNA cluster (Fig. 5C), and all eight proteins gave prominent peaks over the spliced leader locus (Fig. 5D). The fifteen RNAPII promoter-associated factors discussed above also exhibit sharp peaks that appear to coincide with those of the TTR-associated factors (Supplemental Fig. S3C); the significance of this colocation is unknown.

**Figure 5.**
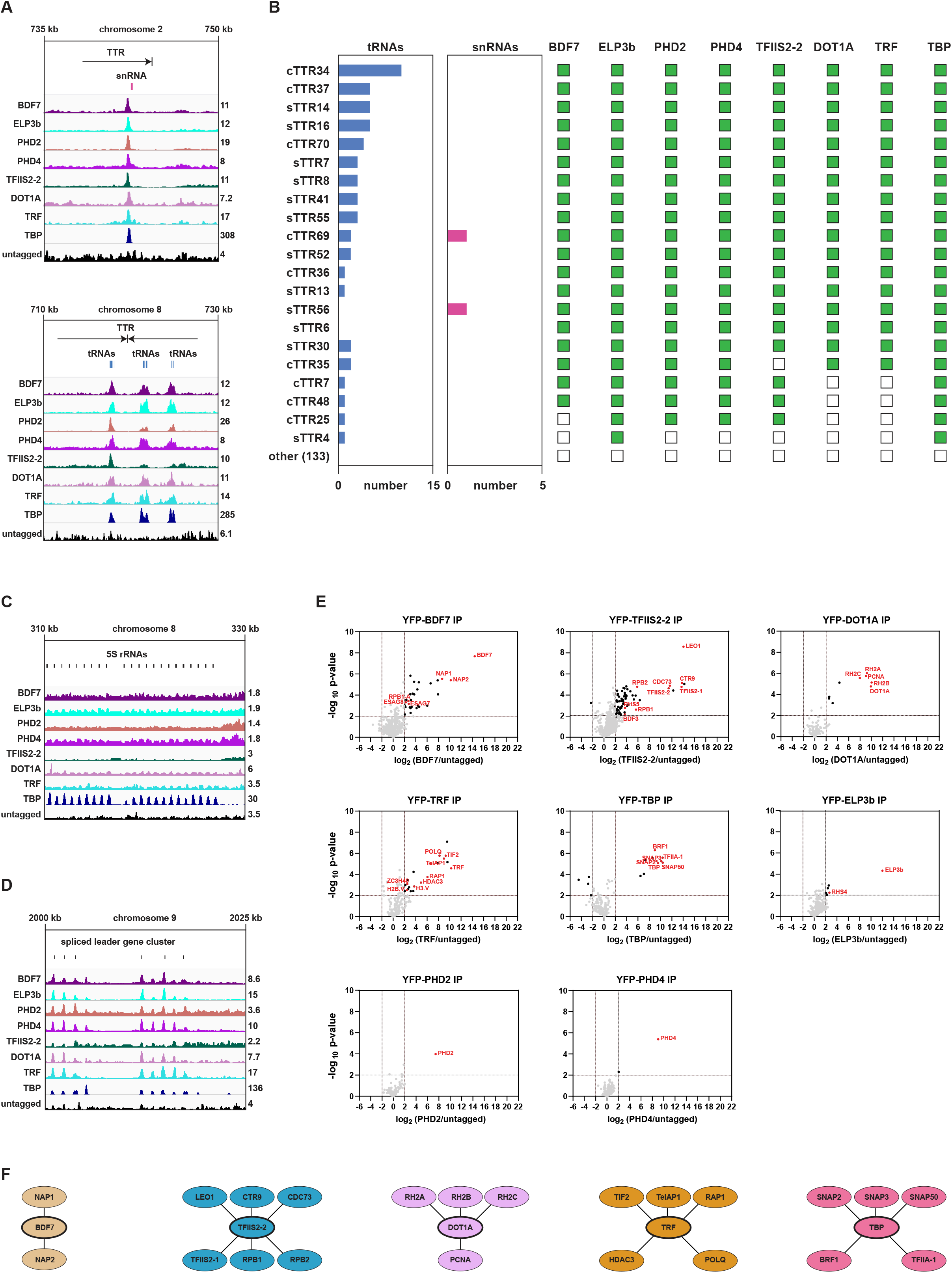
Proteins enriched over RNAPII termination regions coinciding with RNAPIII-transcribed genes define distinct interaction networks. **A**. Examples of protein enrichment over an snRNA (top panel) and tRNAs (bottom panel). Tracks are scaled separately as reads per million (values shown at the end of each track). **B**. Overlap between TTRs, tRNAs, snRNAs and TTR-associated factors. Presence and absence of overlap with specific TTR-associated factors is indicated by green and empty squares, respectively. **C**. Enrichment of the TTR-associated factors at the 5S rRNA gene cluster. **D**. Enrichment of the TTR-associated factors at the spliced leader gene cluster. **E**. YFP-tagged proteins found to be enriched at TTRs were analyses by LC-MS/MS to identify their protein interactions. The data for each plot is based on three biological replicates. Cut-offs used for significance: log_2_ (tagged/untagged) > 2 or < −2 and p < 0.01 (Student’s t-test). Enrichment scores for proteins identified in each affinity selection are presented in Table S5. Significantly enriched proteins are indicated by black or coloured dots. Proteins of interest are indicated by red font. **F**. Key proteins identified as being associated with the indicated YFP-tagged bait proteins (thick oval outlines). Lines denote associations between proteins.

Surprisingly, we found that the terminal (TTAGGG)_n_ telomere-repeat binding protein TRF is enriched at internal chromosomal sites that coincide with peaks of the TTR-associated proteins. We hypothesized that this colocation of TRF with a subset of TTRs could result from the presence of underlying sequence motifs with similarity to canonical (TTAGGG)_n_ telomere repeats, however sequence scrutiny revealed no significant matches. *T. brucei* contains approximately 115 linear chromosomes and consequently it has an abundance of telomeres and telomere binding proteins that cluster at the nuclear periphery (Akiyoshi and Gull, 2013; DuBois et al., 2012; Reis et al., 2018; Yang et al., 2009). The tethering of RNAPIII transcribed genes to the nuclear periphery, as observed in yeast (Chen and Gartenberg, 2014; Iwasaki et al., 2010), would bring them in close proximity to telomeres offering a potential explanation for the association of TRF with these nucleosome depleted regions. In yeasts, both the cohesin and condensin complexes, which shape chromosome architecture, are enriched or loaded at highly transcribed regions such as tRNA genes and specific mutations affect their clustering (D’Ambrosio et al., 2008; Gard et al., 2009; Haeusler et al., 2008; Iwasaki et al., 2010). The *T. brucei* Scc1 cohesin subunit is also enriched over tRNA genes (Muller et al., 2018), perhaps the plethora of factors associated with RNAPIII transcribed genes mediates the formation of nucleosome depleted boundary structures that facilitate termination of transcription at the end of polycistronic units by obstructing the passage of RNAPII. Since none of the eight TTR-associated proteins identified decorate RNAPII TSRs these proteins likely contribute to RNAPII transcription termination and/or facilitate RNAPIII transcription of tRNA and snRNA genes.

### Interaction networks of TTR-associated proteins

To gain more insight into the functional context of the eight proteins identified as being enriched at a subset of TTRs, the same approach used above for TSR-associated proteins was applied to identify interacting factors. Below we detail the interaction networks for TTR-associated proteins and discuss their potential functional implications (Fig. 5E,F; Supplemental Table S5, S6).

Affinity selection of the control TRF telomere binding protein resulted in enrichment of previously identified TRF- and telomere-associated proteins (Reis et al., 2018). The interaction of HDAC3 with TRF and BDF2 and its enrichment over TSRs suggests that it functions at both RNAPII promoters and at subtelomeric regions (Fig. 5E,F). Although ELP3b, PHD2 and PHD4 also exhibit enrichment at RNAPIII genes (Fig. 5A-D), our proteomics analyses detected no significant interaction of other factors with these three proteins (Fig.5E; Supplemental Fig. S5). Indeed, none of the eight TTR-associated proteins identified displayed reciprocal interactions with each other. This lack of crosstalk suggests that each protein performs distinct functions at these locations.

### TFIIS2-2 and the PAF1 complex

TFIIS2-2 was clearly enriched in the vicinity of TTRs and upon affinity selection exhibited strong interaction with PAF1 Complex components (LEO1/Tb927.9.12900, CTR9/Tb927.3.3220, CDC73/Tb927.11.10230) and several RNAPII subunits. Yeast Paf1C acts with TFIIS to enable transcriptional elongation through chromatin templates (Schier and Taatjes, 2020; Van Oss et al., 2017). The accumulation of TFIIS2-2 at TTRs regions presumably reflects the role that these proteins are known to play in transcriptional termination and the 3′ end processing of RNAPII transcripts.

### DOT1A and the RNase H2 complex

The DOT1A and DOT1B histone methyltransferases direct trypanosome H3K76 di- and tri-methylation, respectively (Janzen et al., 2006b). DOT1A is involved in cell cycle progression while DOT1B is necessary for maintaining the silent state of inactive VSGs and for rapid transcriptional VSG switching (Figueiredo et al., 2008; Janzen et al., 2006b). Our analysis revealed that DOT1A associates with all three subunits of the RNase H2 complex (RH2A, RH2B and RH2C). The RH1 and RH2 complexes are necessary for resolving R-loops formed during transcription (Cerritelli and Crouch, 2009). While both RH1 and RH2 complexes are involved in antigenic variation, only RH2 has a role in trypanosome RNAPII transcription (Briggs et al., 2019). Interestingly, a recent study suggests that DOT1B is required to clear R-loops by suppressing RNA-DNA hybrid formation and resulting DNA damage (Eisenhuth et al., 2020). DOT1A may act with RH2 to prevent the accumulation of RNA-DNA hybrids at TTRs and RNAPIII transcribed regions.

### BDF7 and NAP proteins

BDF7 is a Bromodomain protein containing an AAA^+^ ATPase domain, equivalent to Yta7, Abo1 and ATAD2 of budding and fission yeast, and vertebrates, respectively. These proteins have been implicated in altering nucleosome density to facilitate transcription, and Abo1 has recently been shown to bind and mediate H3-H4 deposition onto DNA in vitro (Cho et al., 2019; Lombardi et al., 2011; Murawska and Ladurner, 2020). Affinity selected YFP-BDF7 showed strong association with two Nucleosome Assembly Proteins (Tb927.1.2210, designated NAP1; and Tb927.3.4880, designated NAP2), supporting a potential role for BDF7 as a histone chaperone involved in nucleosome formation. H3.V and H4.V tend to be enriched at the end of polycistronic transcription units where RNAPII transcription is terminated (Siegel et al., 2009), and we find that BDF7 is enriched at a subset of these TTRs (Fig. 5B). It is possible that BDF7 acts with NAP1 and NAP2 to mediate H3.V-H4.V deposition at these locations. Distinct nucleosome depleted regions are formed over tRNA genes and RNAPII transcription is known to terminate in regions coincident with tRNA genes (Maree and Patterton, 2014). Indeed, it has been suggested that tRNA genes may act as boundaries that block the passage of advancing RNAPII into convergent or downstream transcription units (Siegel et al., 2009).

### TBP, BRF1 and RNAPIII genes

TBP (TATA-box related protein) has largely been studied with respect to its role in RNAPII transcription from spliced-leader (SL) RNA gene promoters (Das et al., 2005). However, TBP was previously shown to strongly associate with the TFIIIB component BRF1 (Schimanski et al., 2005) and thus, like BRF1, TBP may also promote RNAPIII transcription (Vélez-Ramírez et al., 2015). Indeed, directed ChIP assays indicate that TBP associates with specific RNAPIII transcribed genes (Ruan et al., 2004; Vélez-Ramírez et al., 2015). Consistent with a dual role, we confirmed that YFP-TBP associates with both SNAP complex components involved in SL transcription and with BRF1. Our ChIP-seq analysis shows that TBP is concentrated in sharp peaks that coincide with both RNAPII promoters for SL RNA genes as well as RNAPIII transcribed tRNA and snRNA genes (Fig. 5A,D). Additionally, TBP was significantly enriched at arrays of RNAPIII transcribed 5S rRNA genes (Fig. 5C). Thus, apart from SL RNA gene promoters, TBP appears to mark all known sites of RNAPIII-directed transcription.

### Concluding Remarks

Protein domain homology allowed the identification of a collection of 68 putative chromatin regulators in *Trypanosoma brucei* that were predicted to act as writers, readers or erasers of histone post-translational modifications. Many of these proteins exhibited a discernible nuclear localization and displayed distinct patterns of association across the genome, frequently coinciding with regions responsible for RNAPII transcription initiation and termination or RNAPIII transcription (Fig. 6). Robust proteomic analysis allowed the interaction networks of these proteins to be identified thereby providing further insight into their possible functions at specific genomic locations by revealing putative complexes that are likely involved in the distinct phases of transcription: initiation, elongation and termination. Counterparts of yeast SWR11 H2A-to-H2A.Z exchange complex and NuA4 HAT complex components were identified, some of which were enriched where H2A.Z is prevalent at TSRs.

**Figure 6.**
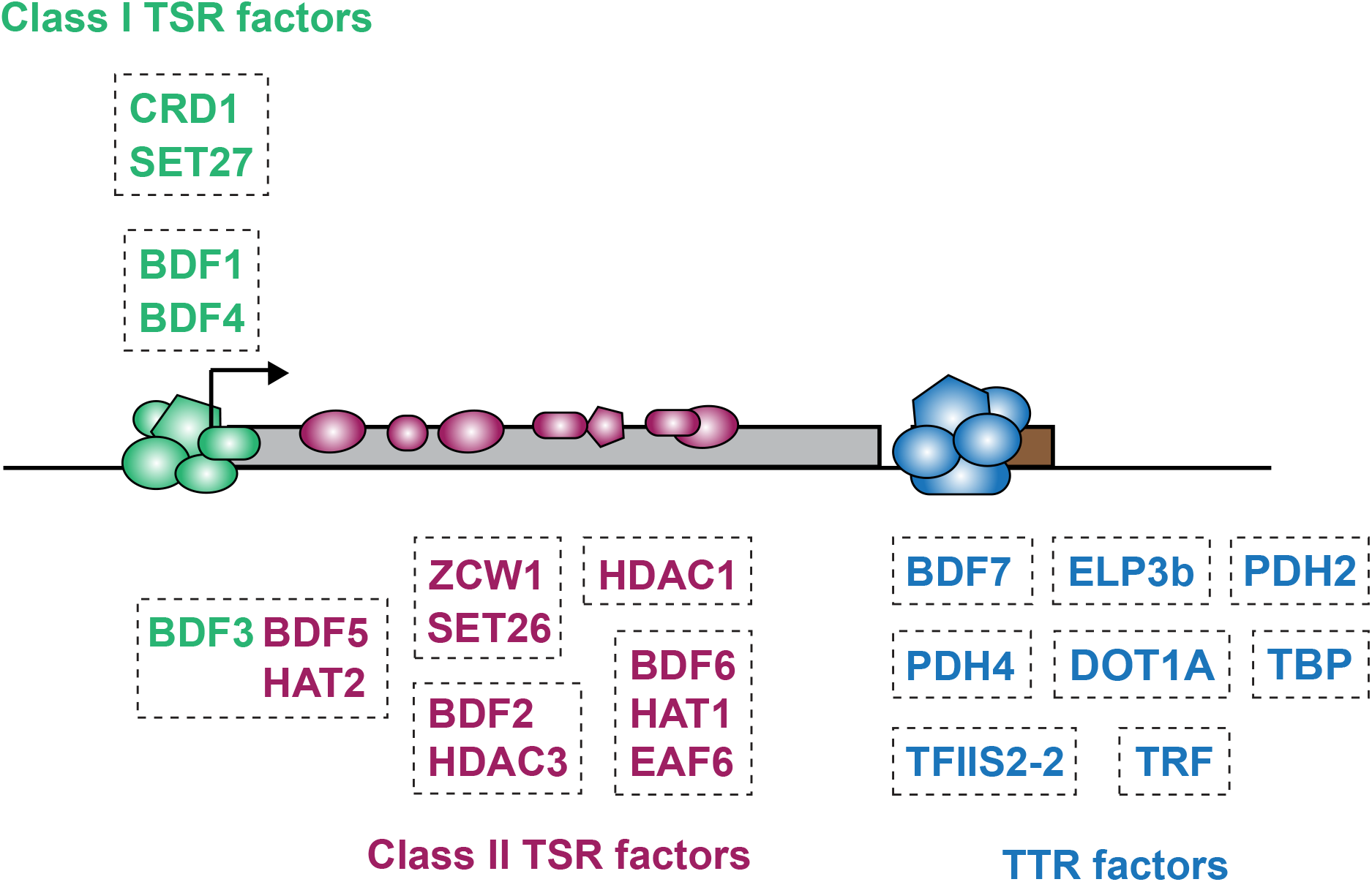
Model depicting distribution of chromatin regulators across a trypanosome polycistronic transcription unit. Diagram shows the five Class I (sharp; green) and ten Class II (broad; purple) TSR-associated factors at a unidirectional RNAPII promoter. The arrows indicate the direction of transcription. The grey rectangle represents a single polycistron. Class II proteins are enriched in the direction of transcription. Eight proteins found at TTRs are shown (blue). tRNA or snRNA RNAPIII transcribed genes (brown). Proteins within boxes were found to interact in affinity purification proteomics analyses.

The data presented provides a comprehensive depiction of the operational context of chromatin writers, readers and erasers at important genomic regulatory elements in this experimentally tractable but divergent eukaryote. Critically, our analyses identify many novel interacting proteins unrelated to, or divergent from, known chromatin regulators of conventional eukaryotes. The identification of these distinct factors highlights the utility of our approach to reveal novelty in trypanosome chromatin regulatory complex composition that differs from the paradigms established using conventional eukaryotic models. Interestingly, although trypanosome gene expression is widely considered to be regulated predominantly at the mRNA and translational level, the enrichment of multiple factors, possibly with antagonistic activities, near transcription initiation and termination regions invokes a more complex regulatory landscape where transcriptional control contributes alongside post-transcriptional mechanisms in trypanosome gene expression.

## Materials and Methods

### Cell culture

Lister 427 bloodstream form *T. brucei* was used throughout this study. Parasites were grown in HMI-9 medium (Hirumi and Hirumi, 1989) at 37°C and 5% CO_2_. Cell lines with YFP-tagged proteins were grown in the presence of 5 µg/ml blasticidin. The density of cell cultures was maintained below 3 × 10^6^ cells/ml.

### Protein tagging

Candidate proteins were tagged endogenously on their N termini with YFP using the pPOTv4 plasmid (Dean et al., 2015). Tagging constructs were produced by fusion PCR of three fragments: a ~500 bp fragment homologous to the end of the 5′ UTR of each gene, a region of the pPOTv4 plasmid containing a blasticidin resistance cassette and a YFP tag, and a ~500 bp fragment homologous to the beginning of the coding sequence of each gene. Fusion constructs were transfected into bloodstream form parasites by electroporation as previously described (Burkard et al., 2007). The cell lines obtained after blasticidin selection were tested for correct integration of the tagging constructs by PCR and for expression of the tagged proteins via western blotting analysis.

### Western analysis

Protein samples (4×10^6^ cell equivalents per lane) were separated on 4%-12% Bis-Tris gels (Thermo Fisher Scientific) and then transferred to nitrocellulose membranes. Membranes were incubated overnight with mouse anti-GFP primary antibody (Sigma-Aldrich) used at 1:1000 dilution. HRP-conjugated anti-mouse secondary antibody (Sigma-Aldrich) was used at 1:2500 dilution. The blots were incubated with ECL Prime Western Blotting Detection Reagent (GE Healthcare) and visualised on films.

### Fluorescent immunolocalization

Cells were fixed with 4% paraformaldehyde for 10 min on ice. Fixation was stopped with 0.1 M glycine. Cells were added to polylysine-coated slides and permeabilised with 0.1% Triton X-100. The slides were blocked with 2% BSA. Rabbit anti-GFP primary antibody (Thermo Fisher Scientific) was used at 1:500 dilution and secondary Alexa fluor 568 anti-rabbit antibody (Thermo Fisher Scientific) was used at 1:1000 dilution. Images were taken with a Zeiss Axio Imager microscope.

### Chromatin immunoprecipitation and sequencing (ChIP-seq)

4 × 10^8^ parasites were fixed with 0.8% formaldehyde for 20 min at room temperature. Cells were lysed and sonicated in the presence of 0.2% SDS for 30 cycles (30 s on, 30 s off) using the high setting on a Bioruptor sonicator (Diagenode). Cell debris were pelleted by centrifugation and SDS in the lysate supernatants was diluted to 0.07%. Input samples were taken before incubating the rest of the cell lysates overnight with 10 µg rabbit anti-GFP antibody (Thermo Fisher Scientific) and Protein G Dynabeads. The beads were washed, and the DNA eluted from them was treated with RNase and Proteinase K. DNA was then purified using a QIAquick PCR Purification Kit (Qiagen) and libraries were prepared using NEXTflex barcoded adapters (Bioo Scientific). The libraries were sequenced on Illumina HiSeq 4000 (Edinburgh Genomics), Illumina NextSeq (Western General Hospital, Edinburgh) or Illumina MiniSeq (Allshire lab). In all cases, 75 bp paired-end sequencing was performed.

### ChIP-seq data analysis

Sequencing reads were de-duplicated and subsequently aligned to the Tb427v9.2 genome (Muller et al., 2018) using Bowtie2 (Langmead and Salzberg, 2012). ChIP samples were normalized to their respective inputs and to library size. CRD1 peak summits were called using MACS2 (Feng et al., 2012) followed by manual filtering of false positives. 10 kb regions upstream and downstream of CRD1 peak summits were divided into 50 bp windows. The metagene plots displayed individual ChIP-seq replicates separately and were generated by summing normalized reads in each 50 bp window and representing them as density centered around CRD1. The average metagene plots were generated analogously, except that the reads around CRD1 peaks were averaged before plotting. Heatmaps represented normalized reads around individual CRD1 peaks and were generated as an average of all replicates for each protein.

### Affinity purification and LC MS/MS proteomic analysis

4 × 10^8^ cells were lysed per IP in the presence of 0.2% NP-40 and 150 mM KCl. Lysates were sonicated briefly (3 cycles, 12 s on, 12 s off) at a high setting in a Bioruptor (Diagenode) sonicator. The soluble and insoluble fractions were separated by centrifugation, and the soluble fraction was incubated for 1 h at 4°C with beads crosslinked to mouse anti-GFP antibody (Roche). Resulting Immunoprecipitates were washed three times with lysis buffer and protein eluted with RapiGest surfactant (Waters) at 55°C for 15 min. Next, filter-aided sample preparation (FASP) (Wiśniewski et al., 2009) was used to digest the protein samples for mass spectrometric analysis. Briefly, proteins were reduced with DTT and then denatured with 8 M Urea in Vivakon spin (filter) column 30K cartridges. Samples were alkylated with 0.05 M IAA and digested with 0.5 μg MS Grade Pierce Trypsin Protease (Thermo Fisher Scientific) overnight, desalted using stage tips (Rappsilber et al., 2007) and resuspended in 0.1%TFA for LC MS/MS. Peptides were separated using RSLC Ultimate3000 system (Thermo Scientific) fitted with an EasySpray column (50 cm; Thermo Scientific) utilising 2-40-95% non-linear gradients with solvent A (0.1% formic acid) and solvent B (80% acetonitrile in 0.1% formic acid). The EasySpray column was directly coupled to an Orbitrap Fusion Lumos (Thermo Scientific) operated in DDA mode. “TopSpeed” mode was used with 3 s cycles with standard settings to maximize identification rates: MS1 scan range - 350-1500mz, RF lens 30%, AGC target 4.0e^5^ with intensity threshold 5.0e^3^, filling time 50 ms and resolution 60000, monoisotopic precursor selection and filter for charge states 2-5. HCD (27%) was selected as fragmentation mode. MS2 scans were performed using Ion Trap mass analyser operated in rapid mode with AGC set to 2.0e^4^ and filling time to 50ms. The resulting shot-gun data were processed using Maxquant 1.3.8. and visualized using Perseus 1.6.0.2 (Tyanova et al., 2016).

### Data Access

All ChIP-seq data generated are available under accesseion number GSE150253 at https://www.ncbi.nlm.nih.gov/geo/query/acc.cgi?acc=GSE150253

## Supporting information

Indexed All Supplemental DATA

## Acknowlegements

We thank members of both the Allshire and Matthews labs for vaulable input and discussion. In particular, we thank Manu Shukla and Sito Torres-Garcia for advice on ChIP-seq; Alison Pidoux and Eleanor Silvester for advice on immunolocalization and microscopy; Marcel Lafos for input on bioinformatics; and Julie Young for supplying HMI-9 media. Additionally, we thank Samuel Dean for the pPOTv4 plasmid used for YFP-tagging. We also thank Edinburgh Genomics (NERC, R8/H10/56; MRC, MR/K001744/1; BBSRC, BB/J004243/1) and Genetics Core, Edinburgh Clinical Research Facility for their valued sequencing services. This research was made possible by core funding to the Wellcome Centre for Cell Biology (203149); a Wellcome 4-year PhD in Cell Biology studentship (102336) to DPS; a Wellcome Senior Research Fellowship (103139) and Wellcome Instrument grant (108504) to J.R; a Wellcome Senior Research Fellowship (202811) to AAJ; a Wellcome Investigator award (103740) to KM; a Wellcome Trust Principal Research Fellowship (095021 and 200885) to RCA; and an MRC Research Grant (MR/T04702X/1) to RCA and KRM.

## Disclosure Declaration

The authors have no conflicts of interest or competing interests to declare.

### Open Access

This research was funded in whole, or in part, by the Wellcome Trust [*Grant numbers as stated above*]. For the purpose of Open Access, the authors have applied a CC BY public copyright license to any Author Accepted Manuscript version arising from this submission.

