## Supplementary material for "A systematic analysis of *Trypanosoma brucei* chromatin factors identifies novel protein interaction networks associated with sites of transcription initiation and termination": Indexed All Supplemental DATA

**Page**
**1** **Supplemental Figure S1. Domain architecture of the candidate proteins.**
**8** **Supplemental Figure S2. Cellular localization of candidate and control *T. brucei* proteins not detected as being enriched over any specific genomic region.**
**11** **Supplemental Figure S3. Examples of ChIP-seq enrichment of TSR- and TTR-associated factors over other genomic regions.**
**13** **Supplemental Figure S4. Average metagene plots.**
**15** **Supplemental Figure S5. Proteomics analyses of affinity selections for several non-chromatin associated candidate proteins.**
**17** **Supplemental Table S1. Protein localization in bloodstream and procyclic form cells.**
**21** **Supplemental Table S2. Functions and phenotypes of previously characterized candidate proteins.**
**22** **Supplemental Table S3. Candidate proteins with no genomic enrichment by ChIP-seq.**
**23** **Supplemental Table S4. Overlap of tRNAs and snRNAs annotated in the assembled Lister 427 genome with TTRs and TTR- associated factors.**
**26** **Supplemental Table S5. Significantly enriched interactors detected by proteomic analysis of affinity selected YFP-tagged proteins**
26 CRD1 IP significantly enriched interactors (over untagged control).
27 SET27 IP significantly enriched interactors (over untagged control).
28 BDF1 IP significantly enriched interactors (over untagged control).
29 BDF4 IP significantly enriched interactors (over untagged control).
30 BDF3 IP significantly enriched interactors (over untagged control).
31 BDF5 IP significantly enriched interactors (over untagged control).
32 HAT2 IP significantly enriched interactors (over untagged control).
33 BDF2 IP significantly enriched interactors (over untagged control).
35 HDAC3 IP significantly enriched interactors (over untagged control).
36 SET26 IP significantly enriched interactors (over untagged control).
38 ZCW1 IP significantly enriched interactors (over untagged control).
40 BDF6 IP significantly enriched interactors (over untagged control).
41 HAT1 IP significantly enriched interactors (over untagged control).
42 EAF6 IP significantly enriched interactors (over untagged control).
43 HDAC1 IP significantly enriched interactors (over untagged control).
44 BDF7 IP significantly enriched interactors (over untagged control).
45 TFIIS2-2 IP significantly enriched interactors (over untagged control).
48 DOT1A IP significantly enriched interactors (over untagged control).
49 TRF IP significantly enriched interactors (over untagged control).
50 TBP IP significantly enriched interactors (over untagged control).
51 ELP3b IP significantly enriched interactors (over untagged control).
52 PHD2 IP significantly enriched interactors (over untagged control).
53 PHD4 IP significantly enriched interactors (over untagged control).
54 PHD1 IP significantly enriched interactors (over untagged control).
55 HAT3 IP significantly enriched interactors (over untagged control).
56 AGO1 IP significantly enriched interactors (over untagged control).
57 NUP110 IP significantly enriched interactors (over untagged control).
60 SET13 IP significantly enriched interactors (over untagged control).
61 SET15 IP significantly enriched interactors (over untagged control).
70 SET20 IP significantly enriched interactors (over untagged control).
71 SET23 IP significantly enriched interactors (over untagged control).
72 SET20 IP significantly enriched interactors (over untagged control).

**Supplemental Table S6. Homology detected for proteins enriched with candidate chromatin regulators.**

CRD1-SET27 Network.
Enriched with YFP-CRD1 affinity selections.
Enriched with YFP-SET27 affinity selections.
BDF1-BDF4 Network.
Enriched with YFP-BDF1 affinity selections.
Enriched with YFP-BDF4 affinity selections.
BDF3-BDF5-HAT2 Network.
Enriched with YFP-BDF3 affinity selections.
Enriched with YFP-BDF5 affinity selections.
Enriched with YFP-HAT2 affinity selections.
BDF2-HDAC3 Network.
Enriched with YFP-BDF2 affinity selections.
Enriched with YFP-HDAC3 affinity selections.
SET26-ZCW1 Network.
Enriched with YFP-SET26 affinity selections.
Enriched with YFP-ZCW1 affinity selections.
BDF6-EAF6-HAT1 Network.
Enriched with YFP-BDF6 affinity selections.
Enriched with YFP-HAT1 affinity selections.
Enriched with YFP-EAF6 affinity selections.

Enriched with YFP-HDAC1 affinity selections.
Enriched with YFP-BDF7 affinity selections.
Enriched with YFP-TFIIS2-2 affinity selections.
Enriched with YFP-DOT1A affinity selections.
Enriched with YFP-TRF affinity selections.
Enriched with YFP-TBP affinity selections.
Enriched with YFP-ELP3b affinity selections.
Enriched with YFP-PHD2 affinity selections.
Enriched with YFP-PHD4 affinity selections.
Enriched with YFP-PHD1 affinity selections.
Enriched with YFP-HAT3 affinity selections.
Enriched with YFP-AGO1 affinity selections.
Enriched with YFP-NUP110 affinity selections.
Frequent non-specific contaminants.

**Supplemental Table S7. *T. brucei* homologs of SWR1-C/SRCAP-C/EP400 complex subunits.**

**Supplemental Table S8. *T. brucei* homologs of NuA4 complex subunits.**

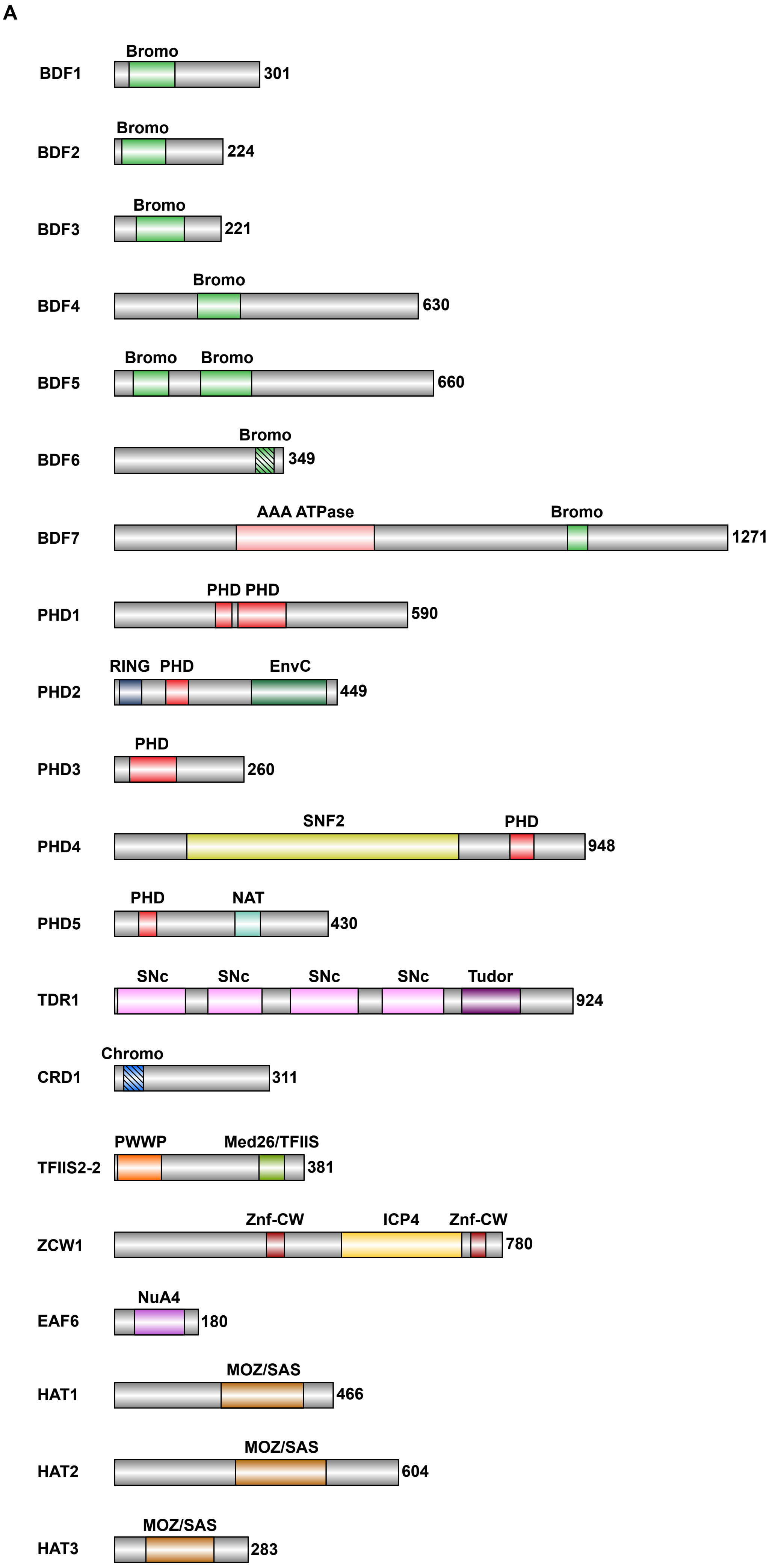


Figure S1 part 1

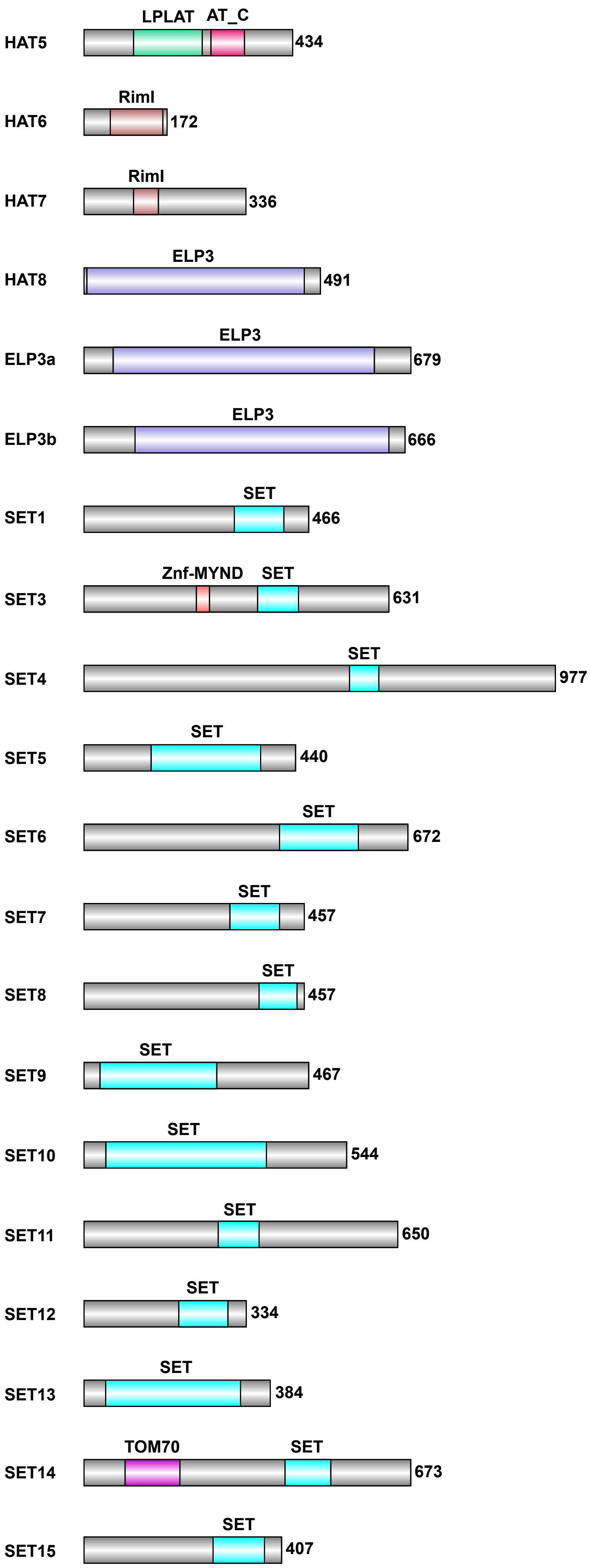


Figure S1 part 2

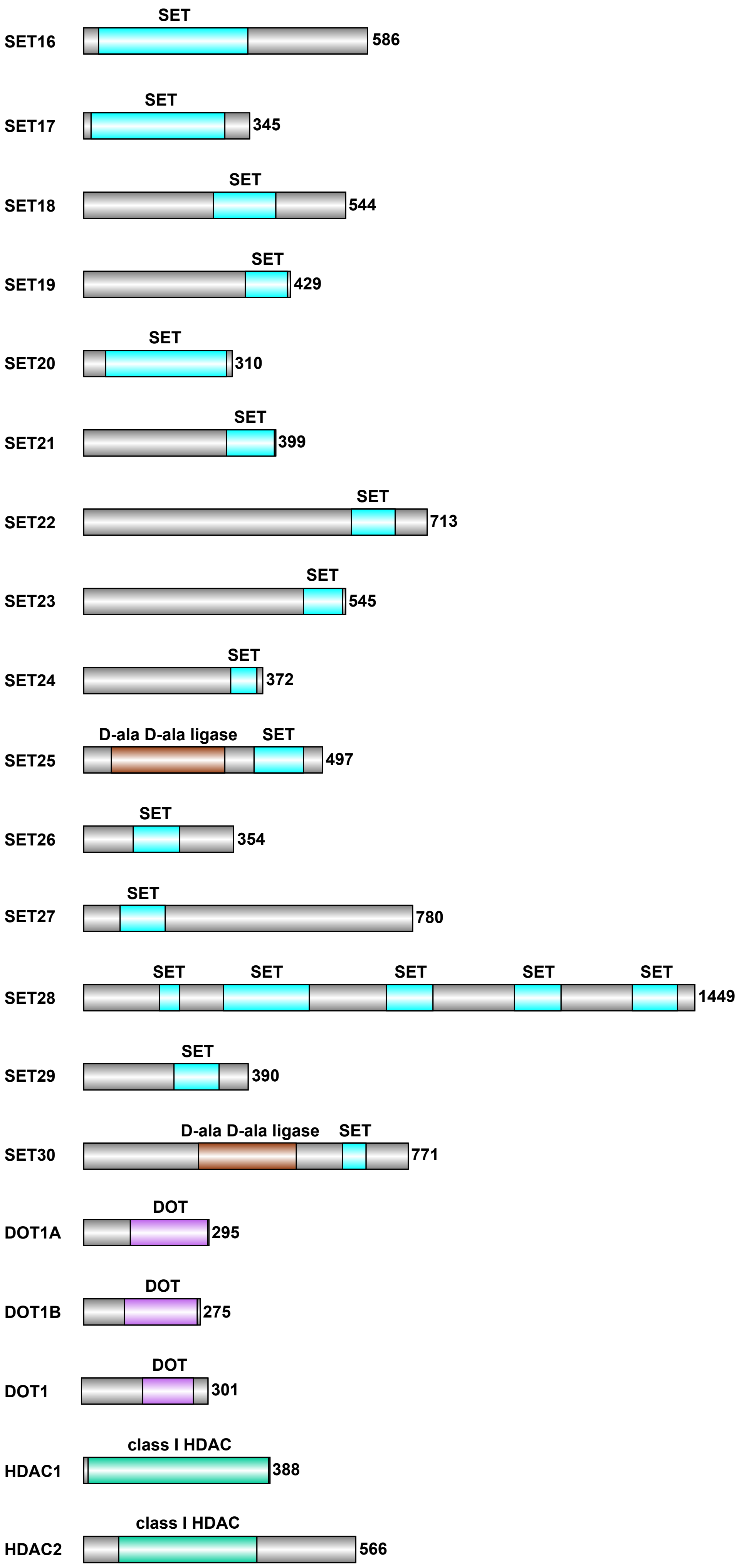


Figure S1 part 3

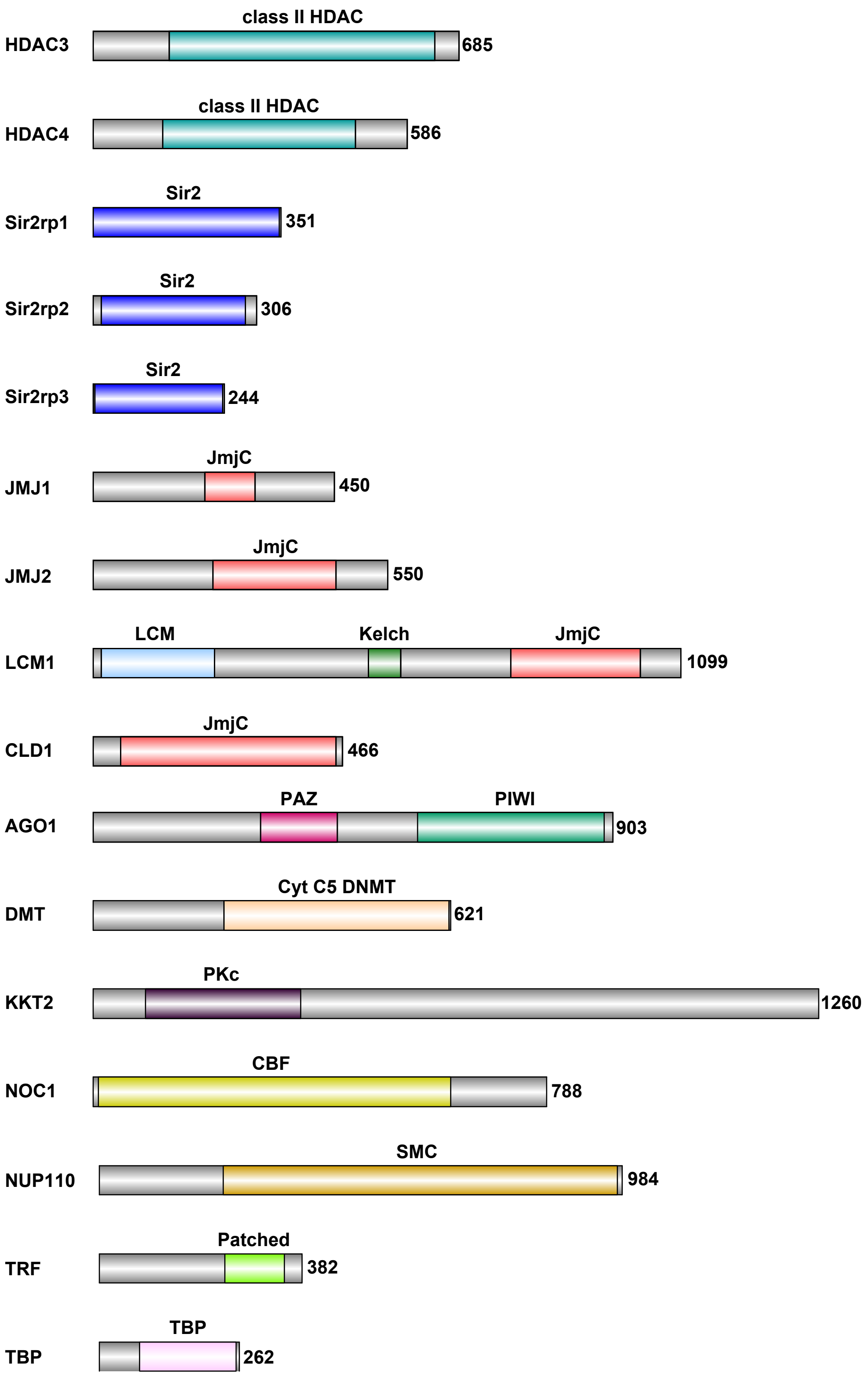


Figure S1 part 4

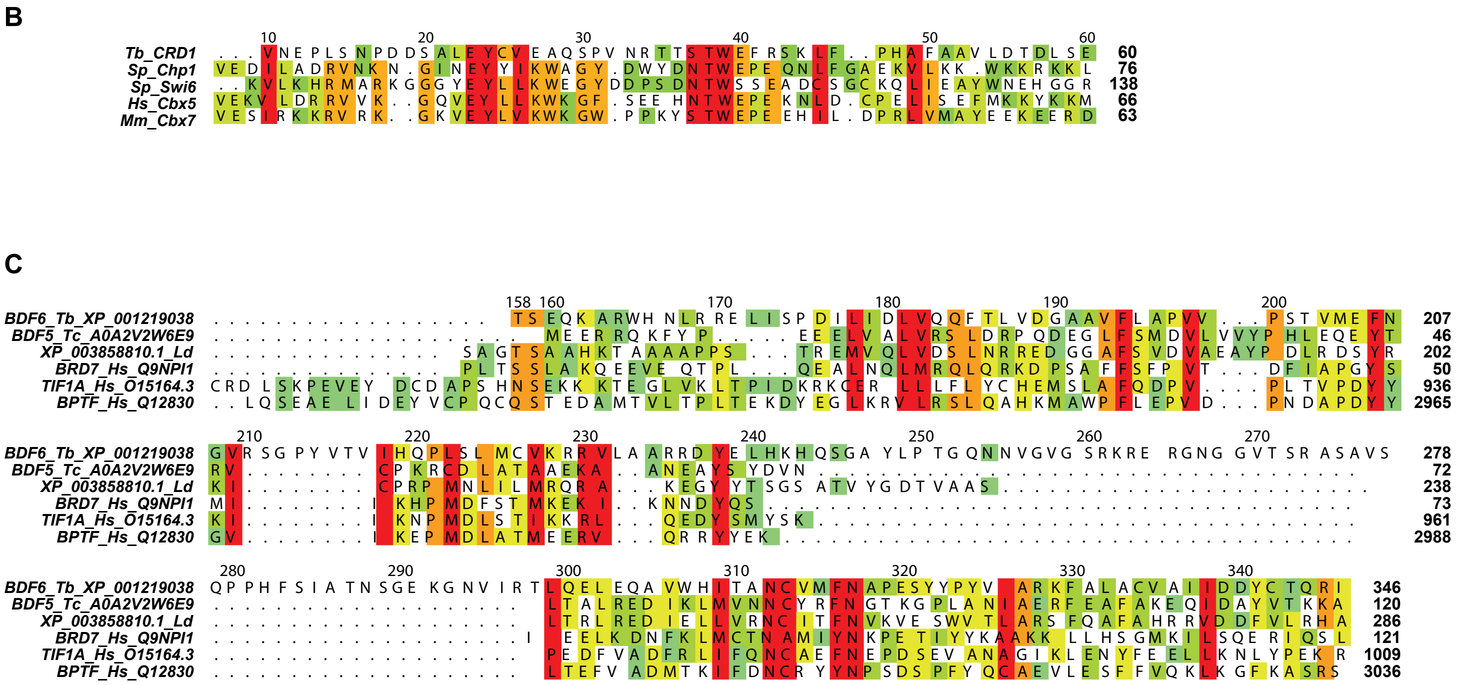


Figure S1 part 5

**Figure S1. Domain architecture of the candidate proteins.**

**A.** A short description of the conserved sequences and domains found in the candidate proteins is provided below. The shaded domains of BDF6 and CRD1 are weakly predicted.

**Bromo** – domain which binds acetylated lysines; found in chromatin-associated proteins and in histone acetyltransferases;

**AAA ATPase** – has chaperone-like functions that aid assembly, function or disassembly of protein complexes;

**PHD (plant homeodomain)** - a $Cys_4$-His-$Cys_3$ zinc finger motif found in nuclear proteins involved in chromatin-mediated transcriptional regulation; some PHD fingers bind methylated histones;

**RING** – a specialised type of zinc finger often found in ubiquitin protein ligases

**EnvC** – bacterial protein which activates cell wall hydrolases and is required for daughter cell separation following cell division;

**SNF2** – domain found in proteins involved in a variety of processes including transcription regulation, DNA repair, DNA recombination and chromatin unwinding;

**NAT** – N-acetyltransferase

**SNc** - Staphylococcal nuclease fold

**Tudor** – domain which can recognise methylated histone lysines and arginines; present in several RNA-binding proteins;

**Chromo (chromatin organisation modifier)** – domain which binds methylated histones; involved in chromatin organisation, specifically heterochromatin formation

**PWWP** – domain which contains a Pro-Trp-Trp-Pro motif; binds methylated histone lysines; found in DNA-binding proteins that function as transcription factors

**Med26/TFIIS** - TFIIS helical bundle-like domain; component of the mediator complex involved in the regulation of RNAPII-transcribed genes;

**Znf-CW** – a zinc finger domain containing conserved Cys and Trp residues; implicated in DNA binding and protein-protein interactions, particularly recognition of methylated histones;

**ICP4** – Herpesvirus protein required for transcription of viral genes; binds DNA in a sequence-specific manner;

**NuA4** – histone acetyltransferase subunit

**MOZ/SAS** - suggested to be homologous to acetyltransferases

**LPLAT** - lysophospholipid acyltransferase;

**AT_C** – domain found at the C-terminus of several acyltransferases

**RimI** – ribosomal protein acetyltransferase found in bacteria; mediates acetylation of N-terminal residues;

**ELP3** - radical SAM enzyme/protein acetyltransferase; this family includes elongator complex protein 3 (ELP3) which is a component of the RNAPII holoenzyme;
**SET** - Su(var)3-9, Enhancer-of-zeste, Trithorax; catalytic domain of lysine methyltransferases;
**Znf-MYND** - MYND-type zinc finger; protein-protein interaction domain
**TOM70** – component of the translocase of outer membrane (TOM) complex involved in mitochondrial import;
**D-ala D-ala ligase** – bacterial enzyme involved in peptidoglycan synthesis and cell wall biogenesis;
**DOT** - Disruptor of telomeric silencing; domain which regulates gene expression via histone methylation
**Class I HDACs** - Zn-dependent histone deacetylases
**Class II HDACs**- Zn-dependent histone deacetylases
**Sir2 HDACs** – sirtuins; NAD-dependent histone deacetylases
**JmjC** – found in metalloenzymes that adopt the cupin fold; function in histone demethylation
**LCM** - leucine carboxyl methyltransferase;
**Kelch** – sequence motif present in proteins with diverse functions including cytoskeletal support and oxidation
**SMC (structural maintenance of chromosomes)** – chromosome segregation protein
**Patched** – transmembrane receptor for Sonic Hedgehog
**TBP (TATA-binding protein)** – part of the DNA-binding transcription factor complex TFIID

**B.** Multiple sequence alignment of the putative TbCRD1 chromodomain with *S. pombe* Chp2 and Swi6; Human Cbx5 and Mouse Cbx7.
**C.** Multiple sequence alignments of the TbBDF6 Bromo domain with indicated protein accession numbers including Human BRD7, TIF1A and BPTF.
Level of sequence similarity is indicated as a color gradient from cyan (low similarity) to red (high similarity).

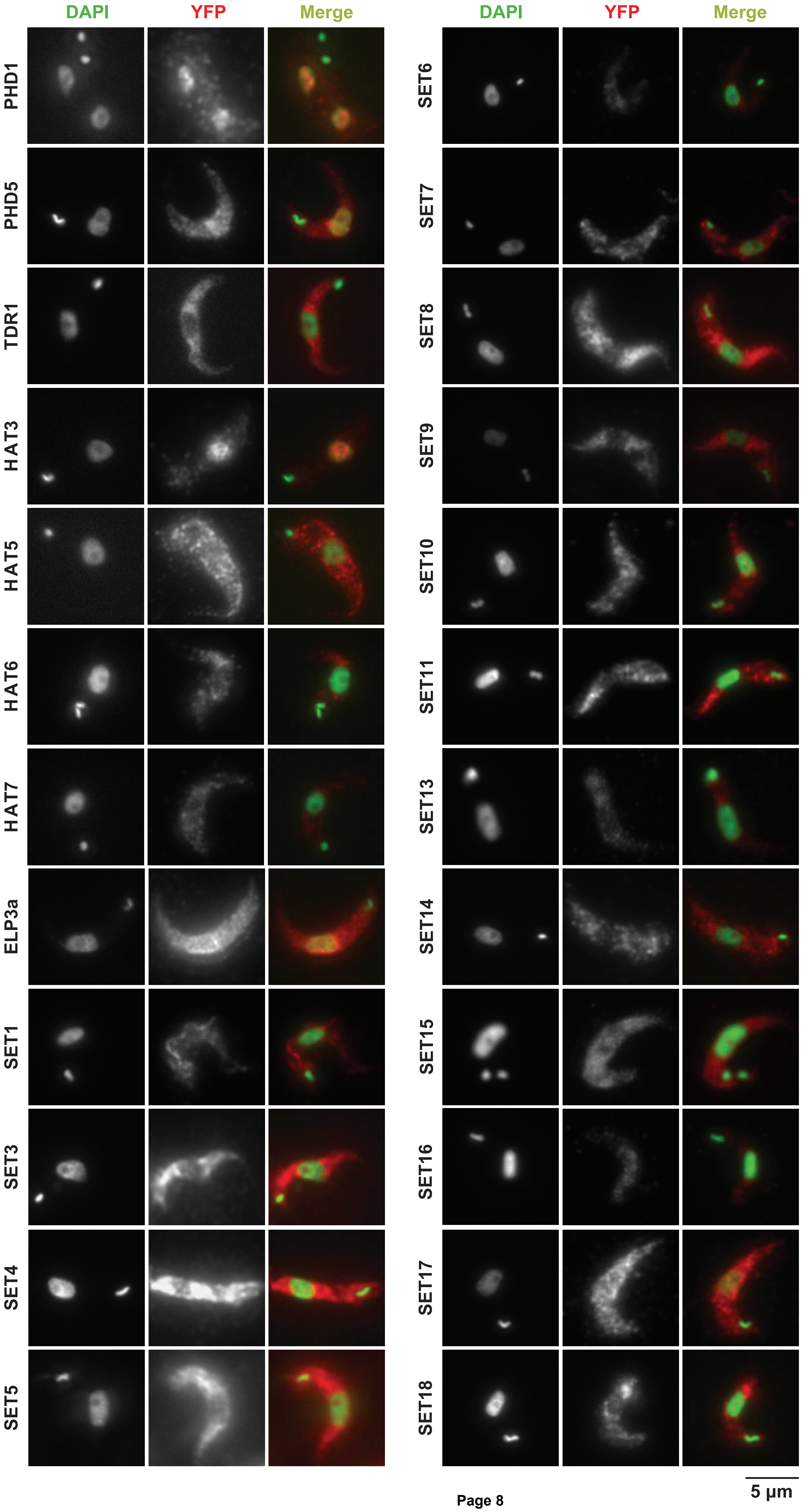


Figure S2 part 1

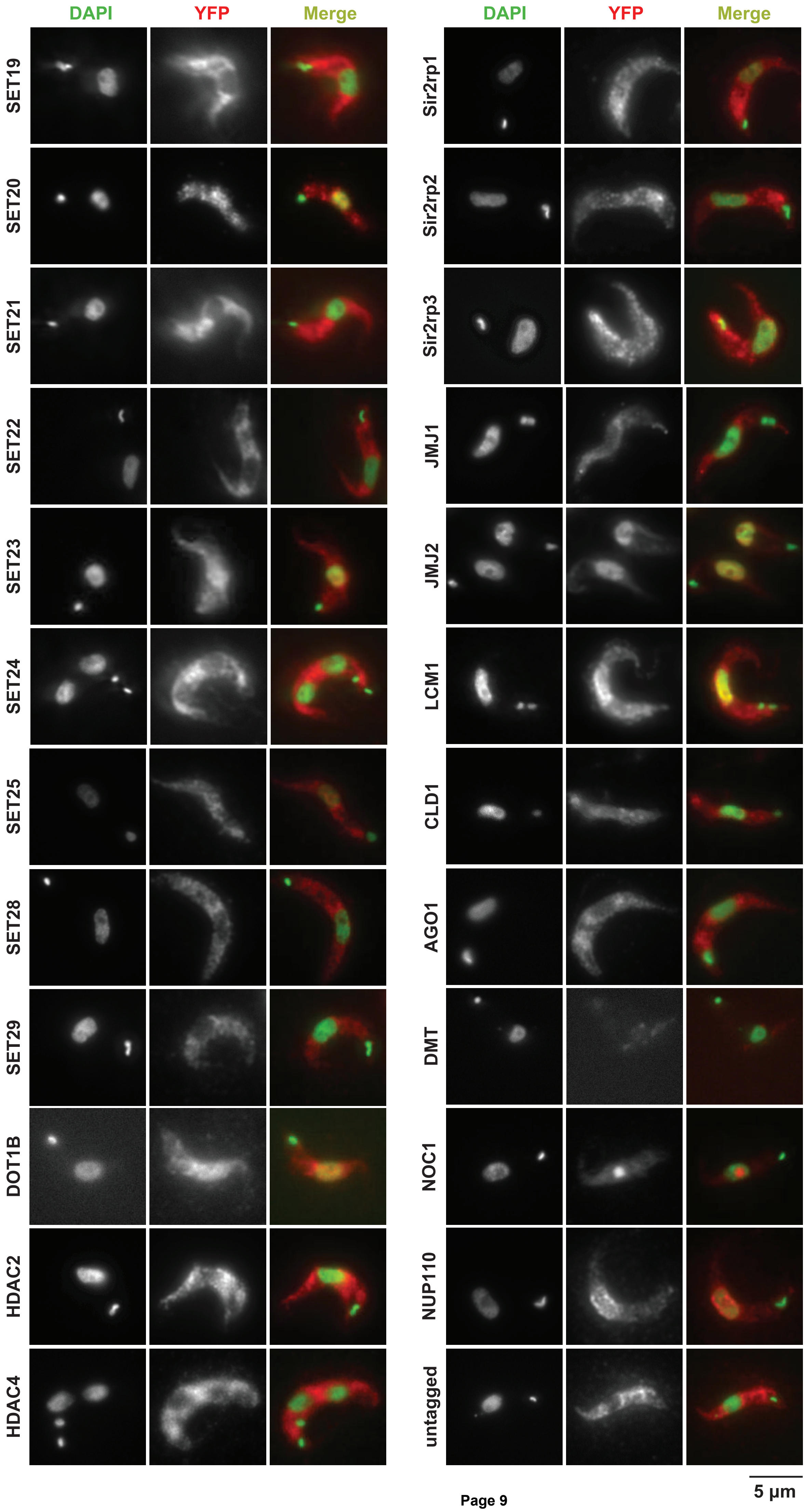


Figure S2 part 2

**Figure S2. Cellular localization of candidate and control *T. brucei* proteins not detected as being enriched over any specific genomic region.**

The indicated YFP-tagged proteins expressed in bloodstream Lister 427 cells from their endogenous genomic loci were detected with an anti-GFP primary antibody and an Alexa Fluor 568 labelled secondary antibody. Nuclear and kinetoplast (mitochondrial) DNA were stained with DAPI. Staining of untagged 427 parasites serves as a negative control. Representative images are shown for each protein for which no specific ChIP-seq pattern was detected. Bar = 5 μm.

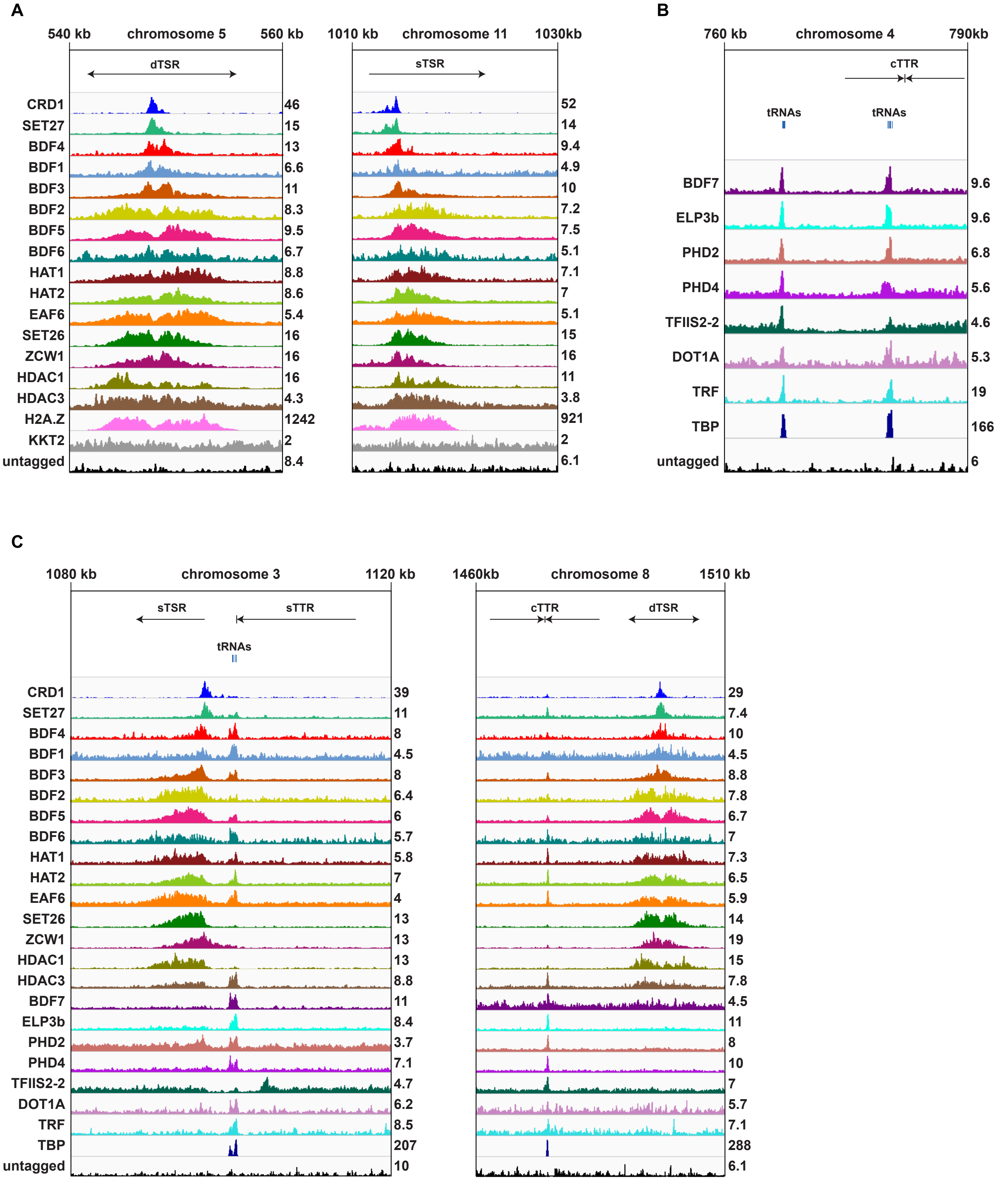
A
540 kb
chromosome 5
560 kb
dTSR
1010 kb
chromosome 11
1030kb
sTSR
CRD1
SET27
BDF4
BDF1
BDF3
BDF2
BDF5
BDF6
HAT1
HAT2
EAF6
SET26
ZCW1
HDAC1
HDAC3
H2A.Z
KKT2
untagged
46 15 13 6.6 11 8.3 9.5 6.7 8.8 8.6 5.4 16 16 16 4.3 1242 2 8.4
52 14 9.4 4.9 10 7.2 7.5 5.1 7.1 7 5.1 15 16 11 3.8 921 2 6.1
B
760 kb
chromosome 4
790kb
cTTR
tRNAs
tRNAs
BDF7
ELP3b
PHD2
PHD4
TFIIS2-2
DOT1A
TRF
TBP
untagged
9.6 9.6 6.8 5.6 4.6 5.3 19 166 6
C
1080 kb
chromosome 3
1120 kb
sTSR
sTTR
tRNAs
1460kb
chromosome 8
1510 kb
cTTR
dTSR
CRD1
SET27
BDF4
BDF1
BDF3
BDF2
BDF5
BDF6
HAT1
HAT2
EAF6
SET26
ZCW1
HDAC1
HDAC3
BDF7
ELP3b
PHD2
PHD4
TFIIS2-2
DOT1A
TRF
TBP
untagged
39 11 8 4.5 8 6.4 6 5.7 5.8 7 4 13 13 13 8.8 11 8.4 3.7 7.1 4.7 6.2 8.5 207 10
29 7.4 10 4.5 8.8 7.8 6.7 7 7.3 6.5 5.9 14 19 15 7.8 4.5 11 8 10 7 5.7 7.1 288 6.1

Figure S3

**Figure S3. Examples of ChIP-seq enrichment of TSR- and TTR-associated factors over other genomic regions**

Tracks are scaled separately as reads per million (values shown at the end of each track).

**A.** Left, enrichment at a bidirectional/divergent TSR.
Right, enrichment at a unidirectional/single TSR.

**B.** Enrichment at two tRNA clusters, one of which overlaps a convergent TTR.

**C.** Coincidence of peaks of the TSR- and TTR-associated factors.

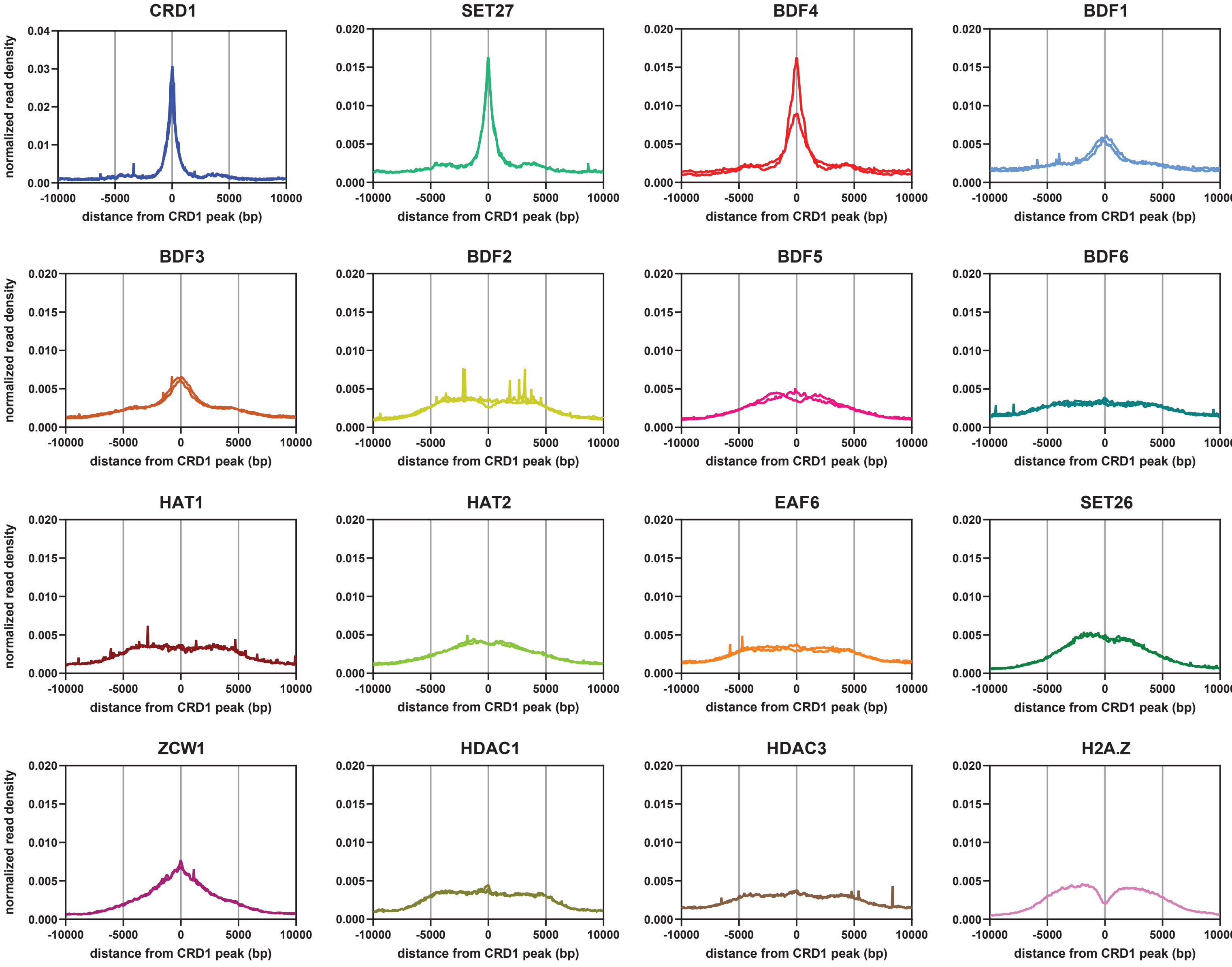
CRD1
SET27
BDF4
BDF1
BDF3
BDF2
BDF5
BDF6
HAT1
HAT2
EAF6
SET26
ZCW1
HDAC1
HDAC3
H2A.Z
normalized read density
distance from CRD1 peak (bp)
-10000
-5000
0
5000
10000
0.00
0.01
0.02
0.03
0.04
0.000
0.005
0.010
0.015
0.020

Figure S4

**Figure S4. Average metagene plots.**

Normalized reads around CRD1 peak summits were averaged and plotted as density.

Plots show separately data from individual ChIP-seq replicates.

Note the different scale for CRD1.

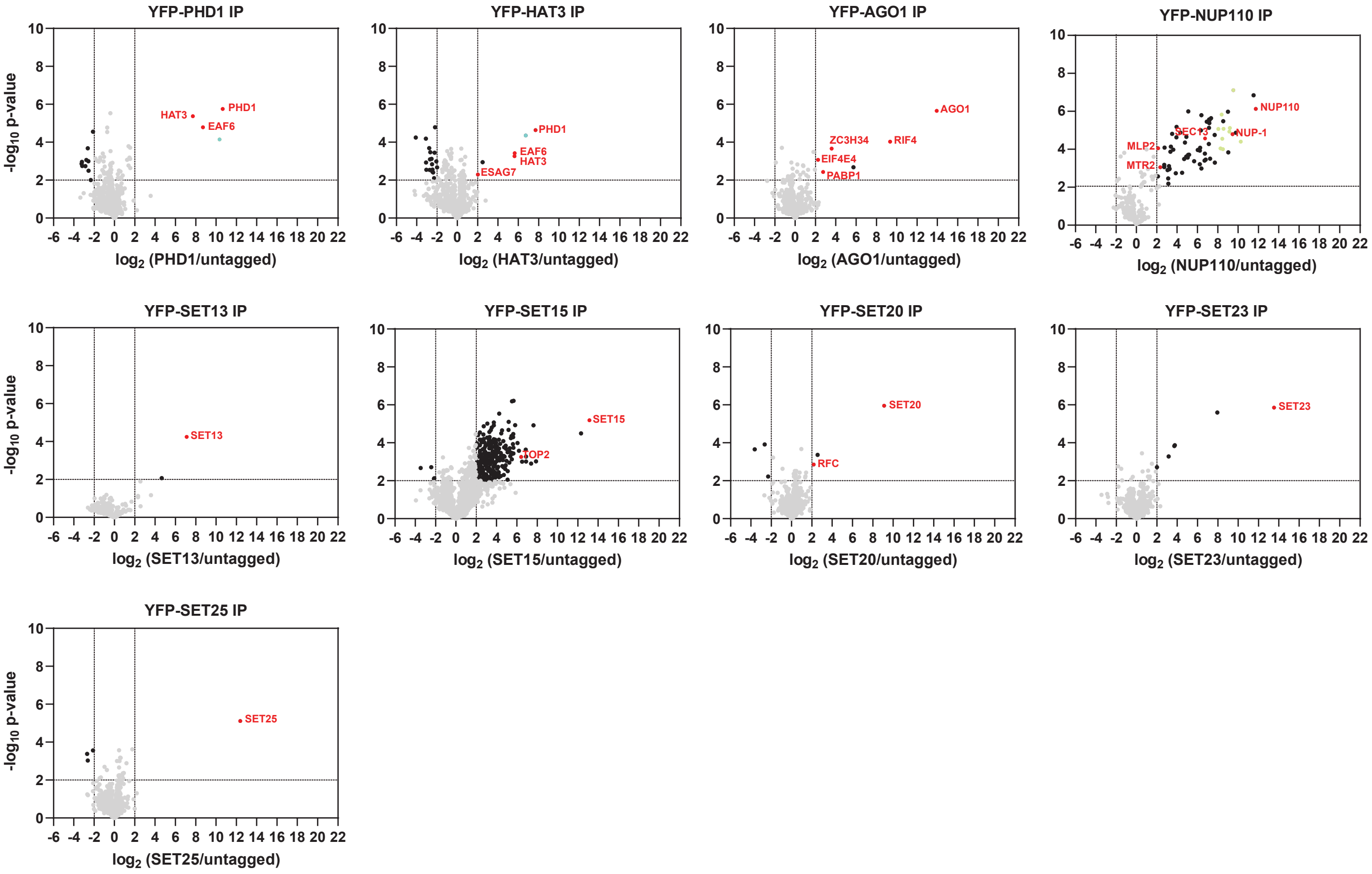
YFP-PHD1 IP
HAT3
PHD1
EAF6
-log10 p-value
log2 (PHD1/untagged)
YFP-HAT3 IP
PHD1
EAF6
HAT3
ESAG7
log2 (HAT3/untagged)
YFP-AGO1 IP
AGO1
ZC3H34
RIF4
EIF4E4
PABP1
log2 (AGO1/untagged)
YFP-NUP110 IP
NUP110
MLP2
SEC13
NUP-1
MTR2
log2 (NUP110/untagged)
YFP-SET13 IP
SET13
log2 (SET13/untagged)
YFP-SET15 IP
SET15
TOP2
log2 (SET15/untagged)
YFP-SET20 IP
SET20
RFC
log2 (SET20/untagged)
YFP-SET23 IP
SET23
log2 (SET23/untagged)
YFP-SET25 IP
SET25
log2 (SET25/untagged)

Figure S5

2

**Figure S5. Proteomics analyses of affinity selections for several non-chromatin associated candidate proteins.**

Data for each plot is based on three biological replicates. Cut-offs used for significance: $log_2$ (tagged/untagged) > 2 or < -2 and $p < 0.01$ (Student's t-test). Enrichment scores for proteins identified in each affinity selection are presented in Table S5. Significantly enriched proteins are indicated by black or coloured dots. Proteins of interest are indicated by red font. Blue dots correspond to some proteins identified also in the BDF6, HAT1 and EAF6 affinity purifications (see Figure 4A). The ten most significant nucleoporins in the YFP-NUP110 affinity selection are marked in pale green.

### Table S1. Protein localization in bloodstream and procyclic form cells

[1] RIT-seq data from Alsford et al (2011). Scoring matrix key: 0-0-0-0 group - significant loss of fitness in all four experiments; 0-0-1-0 group - significant loss of fitness in BF experiments only; 1-1-0-0 group - significant loss of fitness in PF experiments only; 1-1-1-0 group - significant loss of fitness in the differentiation (BF to PF) experiment only; 1-1-1-1 group - no significant loss of fitness in any experiment; none of the above - other groups.

[2] Data from www.tryptag.org was accessed on 04 February 2021.

HAT2 and HAT4 were found to be identical, as were SET2 and SET19. Thus, only HAT2 and SET19 are included in this dataset.

| Protein name | TriTryp ID | UniProt ID | Protein domain/family | [1]RIT-seq scoring matrix | Localization in this study (BF) | [2]TrypTag localization (PF) |
|---|---|---|---|---|---|---|
| BDF1 | Tb927.10.8150 | Q38AE9 | Bromo | 1-1-1-1 | nucleus | N-terminal tag: nuclear lumen; C-terminal tag: nucleus, cytoplasm |
| BDF2 | Tb927.10.7420 | Q38AM1 | Bromo | 1-1-1-1 | nucleus | N-terminal tag: nucleus |
| BDF3 | Tb927.11.10070 | Q383S2 | Bromo | other | nucleus | N-terminal tag and C-terminal tag: nucleoplasm |
| BDF4 | Tb927.7.4380 | Q57UR8 | Bromo | other | nucleus and cytoplasm | - |
| BDF5 | Tb927.11.13400 | Q382J7 | Bromo | 0-0-1-0 | nucleus | N-terminal tag: nucleoplasm |
| BDF6 | Tb927.1.3400 | Q4GYN3 | Bromo | other | nucleus | N-terminal tag: nucleoplasm; C-terminal tag: nucleoplasm, cytoplasm |
| BDF7 | Tb927.11.6350 | Q385D4 | Bromo | 1-1-1-1 | nucleus | N-terminal tag: nucleus; C-terminal tag: nucleoplasm |
| PHD1 | Tb927.10.9930 | Q389Y3 | PHD | 1-1-1-1 | nucleus | N-terminal tag: nuclear lumen; C-terminal tag: nucleoplasm |
| PHD2 | Tb11.v5.0388 | Q387D3 | PHD | other | nucleus | N-terminal tag and C-terminal tag: nuclear lumen |
| PHD3 | Tb11.v5.0794 | - | PHD | - | tagging unsuccessful | - |
| PHD4 | Tb927.3.2140 | Q57Z97 | PHD | 1-1-1-0 | nucleus | C-terminal tag: nucleoplasm, cytoplasm |
| PHD5 | Tb927.11.5870 | Q385I1 | PHD | 1-1-1-1 | nucleus and cytoplasm | N-terminal tag and C-terminal tag: nucleus and cytoplasm |
| TDR1 | Tb927.11.14190 | Q382B8 | Tudor | 1-1-1-1 | cytoplasm | N-terminal tag: cytoplasm, flagellar cytoplasm, nuclear lumen |
| CRD1 | Tb11.v5.0267 | Q57X70 | Chromo | 1-1-1-1 | nucleus | - |
| TFIIS2-2 | Tb927.2.3480 | Q586Y0 | PWWP | other | nucleus | N-terminal tag and C-terminal tag: nucleoplasm |
| ZCW1 | Tb927.10.11720 | Q389G9 | Znf-CW | other | nucleus | N-terminal tag and C-terminal tag: nucleoplasm |
| EAF6 | Tb927.9.2910 | Q38FN2 | NuA4 complex component | 1-1-1-1 | nucleus | N-terminal tag: nucleoplasm; C-terminal tag: nucleus, cytoplasm |
| HAT1 | Tb927.7.4560 | D6XIY7 | MYST | 1-1-1-1 | nucleus | - |
| HAT2 | Tb927.11.11530 | Q383C5 | MYST | 1-1-1-1 | nucleus | N-terminal tag and C-terminal tag: nucleoplasm |
| HAT3 | Tb927.10.8310 | Q38AD3 | MYST | 0-0-0-0 | nucleus | N-terminal tag: nucleus; C-terminal tag: nucleus, cytoplasm, flagellar cytoplasm |

| Protein name | TriTryp ID | UniProt ID | Protein domain/family | [1]RIT-seq scoring matrix | Localization in this study (BF) | [2]TrypTag localization (PF) |
|---|---|---|---|---|---|---|
| HAT5 | Tb927.5.2280 | Q582D7 | LPLAT | 1-1-0-0 | nucleus and cytoplasm | C-terminal tag: cytoplasm, flagellar cytoplasm, nuclear lumen |
| HAT6 | Tb927.1.4490 | Q4GYC9 | GNAT | 1-1-1-0 | cytoplasm | C-terminal tag: cytoplasm |
| HAT7 | Tb927.10.12830 | Q388W3 | GNAT | 1-1-1-1 | cytoplasm | N-terminal tag and C-terminal tag: cytoplasm |
| HAT8 | Tb11.v5.0520 | - | GNAT | - | tagging unsuccessful | - |
| ELP3a | Tb927.8.5770 | Q57X05 | GNAT | 1-1-1-1 | nuclear periphery and cytoplasm | N-terminal tag: nuclear envelope; C-terminal tag: nucleus, cytoplasm |
| ELP3b | Tb927.8.3310 | Q57YY2 | GNAT | other | nucleus | N-terminal tag: nucleus, spindle; C-terminal tag: nucleoplasm |
| SET1 | Tb11.v5.0422 | Q38BE1 | SET | 1-1-1-1 | cytoplasm | - |
| SET3 | Tb927.1.4720 | Q4GYA6 | SET | 1-1-1-1 | cytoplasm | C-terminal tag: cytoplasm |
| SET4 | Tb927.10.11130 | Q389M6 | SET | 1-1-1-1 | cytoplasm | N-terminal tag and C-terminal tag: cytoplasm |
| SET5 | Tb927.10.12880 | Q388V8 | SET | 1-1-1-1 | cytoplasm | N-terminal tag: cytoplasm; C-terminal tag: mitochondrion, kinetoplast |
| SET6 | Tb927.10.3730 | Q38BM6 | SET | other | cytoplasm | N-terminal tag: cytoplasm |
| SET7; similar to SET1 | Tb927.10.4600 | - | SET | - | cytoplasm | N-terminal tag and C-terminal tag: paraflagellar rod, cytoplasm |
| SET8 | Tb927.10.8060 | Q38AF8 | SET | 0-0-1-0 | cytoplasm | N-terminal tag: cytoplasm, flagellar cytoplasm; C-terminal tag: cytoplasm |
| SET9 | Tb927.10.9680 | Q38A07 | SET | 1-1-1-1 | cytoplasm | N-terminal tag: cytoplasm, flagellar cytoplasm, nuclear lumen |
| SET10 | Tb927.11.13560 | Q382I1 | SET | 0-0-0-0 | nucleus and cytoplasm | - |
| SET11 | Tb927.11.5120 | Q385Q6 | SET | 1-1-1-1 | cytoplasm | - |
| SET12 | Tb927.3.750 | Q57WV8 | SET | other | protein not detected by western analysis | N-terminal tag: endocytic, cytoplasm |
| SET13 | Tb927.4.2440 | Q584E3 | SET | other | cytoplasm | N-terminal tag: nucleolus, nucleus |
| SET14 | Tb927.4.3310 | Q584A8 | SET | other | cytoplasm | N-terminal tag: cytoplasm, nucleoplasm; C-terminal tag: cytoplasm, nucleoplasm, flagellum |
| SET15 | Tb927.5.2770 | Q57ZV3 | SET | other | cytoplasm | N-terminal tag and C-terminal tag: cytoplasm, flagellar cytoplasm, nucleus, nucleolus |
| SET16 | Tb927.5.3500 | Q57UB6 | SET | 1-1-0-0 | nucleus and cytoplasm | - |
| SET17 | Tb927.6.3610 | Q585V9 | SET | other | nucleus and cytoplasm | - |
| SET18 | Tb927.6.910 | Q585F5 | SET | 1-1-1-1 | cytoplasm | - |

| Protein name | TriTryp ID | UniProt ID | Protein domain/family | [1]RIT-seq scoring matrix | Localization in this study (BF) | [2]TrypTag localization (PF) |
|---|---|---|---|---|---|---|
| SET19 | Tb927.7.5620 | Q582H7 | SET | 1-1-1-1 | cytoplasm | - |
| SET20 | Tb927.8.2490 | Q57XE0 | SET | 1-1-1-1 | nucleus and cytoplasm | N-terminal tag: cytoplasm, flagellar cytoplasm, nuclear lumen;<br>C-terminal tag: mitochondrion |
| SET21 | Tb927.8.2690 | Q57XC0 | SET | 1-1-1-1 | cytoplasm | N-terminal tag: cytoplasm;<br>C-terminal tag: nucleoplasm, cytoplasm, flagellar cytoplasm |
| SET22 | Tb927.8.2710 | Q57XB8 | SET | 1-1-1-1 | cytoplasm | - |
| SET23 | Tb927.8.6470 | Q57YP8 | SET | 1-1-1-1 | nucleus and cytoplasm | N-terminal tag: cytoplasm, flagellar cytoplasm, nuclear lumen |
| SET24 | Tb927.8.6530 | Q57TW0 | SET | 1-1-1-1 | cytoplasm | N-terminal tag: cytoplasm;<br>C-terminal tag: kinetoplast, mitochondrion |
| SET25 | Tb927.9.1510 | Q38FZ5 | SET | other | nucleus and cytoplasm | - |
| SET26 | Tb927.10.8100 | Q38AF4 | SET | 1-1-1-1 | nucleus and cytoplasm | N-terminal tag and C-terminal tag: nucleoplasm, cytoplasm; |
| SET27 | Tb927.9.13470 | Q38D80 | SET | 1-1-1-1 | nucleus and cytoplasm | - |
| SET28 | Tb927.9.10100 | Q38DZ4 | SET | 1-1-1-1 | cytoplasm | - |
| SET29 | Tb927.3.3370 | Q57V49 | SET | other | cytoplasm | N-terminal tag: cytoplasm;<br>C-terminal tag: cytoplasm, flagellar cytoplasm, nuclear lumen |
| SET30 | Tb927.3.1860 | Q57ZC5 | SET | 1-1-1-0 | protein not detected by western analysis | N-terminal tag: cytoplasm;<br>C-terminal tag: kinetoplast, mitochondrion |
| DOT1A | Tb927.8.1920 | Q581Z0 | DOT | 1-1-1-1 | nucleus and cytoplasm | - |
| DOT1B | Tb927.1.570 | Q4GZF2 | DOT | 1-1-1-1 | nucleus and cytoplasm | N-terminal tag: nuclear lumen;<br>C-terminal tag: nuclear lumen, cytoplasm |
| DOT1 | Tb927.11.3200 | Q386J1 | DOT | 1-1-1-1 | protein not detected by western analysis | - |
| HDAC1 | Tb927.10.1680 | Q38C74 | Class I HDAC | other | nucleus and cytoplasm | C-terminal tag: cytoplasm, flagellar cytoplasm, nuclear lumen |
| HDAC2 | Tb927.11.15600 | Q381M6 | Class I HDAC | 1-1-1-1 | cytoplasm | N-terminal tag: cytoplasm, flagellar cytoplasm, nuclear lumen;<br>C-terminal tag: cytoplasm |
| HDAC3 | Tb927.2.2190 | Q586J9 | Class II HDAC | 0-0-0-0 | nucleus | N-terminal tag: nucleolus, nucleoplasm;<br>C-terminal tag: nucleoplasm |
| HDAC4 | Tb927.5.2900 | D6XGM6 | Class II HDAC | 1-1-1-0 | cytoplasm | - |
| Sir2rp1 | Tb927.7.1690 | Q57V41 | Sir2 HDAC | 1-1-1-1 | nucleus and cytoplasm | - |
| Sir2rp2 | Tb927.8.3140 | Q57YZ9 | Sir2 HDAC | 1-1-1-1 | cytoplasm | - |
| Sir2rp3 | Tb927.4.2520 | Q584D5 | Sir2 HDAC | 0-0-0-0 | nucleus and cytoplasm | N-terminal tag: cytoplasm, flagellar cytoplasm, nuclear lumen;<br>C-terminal tag: mitochondrion |
| JMJ1 | Tb927.11.1760 | Q386X8 | JmjC | other | cytoplasm | N-terminal tag: cytoplasm;<br>C-terminal tag: cytoplasm, flagellar cytoplasm, nucleoplasm |

| Protein name | TriTryp ID | UniProt ID | Protein domain/family | [1]RIT-seq scoring matrix | Localization in this study (BF) | [2]TrypTag localization (PF) |
|---|---|---|---|---|---|---|
| JMJ2 | Tb927.11.2000 | Q386V9 | JmjC | 0-0-0-0 | nucleus | N-terminal tag and C-terminal tag: nuclear lumen |
| LCM1 | Tb927.9.12780 | Q38DD6 | JmjC | 1-1-1-1 | nucleus and cytoplasm | N-terminal tag: cytoplasm |
| CLD1 | Tb927.7.660 | Q57VR2 | JmjC | 1-1-1-1 | cytoplasm | C-terminal tag: cytoplasm |
| AGO1 | Tb927.10.10850 | Q389P5 | PAZ and PIWI | 1-1-1-1 | cytoplasm | C-terminal tag: cytoplasm |
| DMT | Tb927.3.1360 | Q57XW9 | DNMT | 1-1-1-1 | cytoplasm (weak signal) | N-terminal tag: cytoplasm (weak signal); C-terminal tag: cytoplasm (weak signal), flagellar cytoplasm |
| KKT2 | Tb927.11.10520 | Q383M7 | Protein kinase | 0-0-1-0 | dots in nucleus | C-terminal tag: nucleoplasm, cytoplasm, kinetochore |
| NOC1 | Tb927.11.2120 | Q386U7 | CBF/Mak21 | 0-0-1-0 | nucleolus | N-terminal tag and C-terminal tag: nucleolus |
| NUP110/ MLP1 | Tb927.11.330 | Q387L8 | SMC | 0-0-1-0 | nuclear pore | N-terminal tag: nuclear pore; C-terminal tag: cytoplasm |
| TRF | Tb927.10.12850 | Q388W1 | Patched | 0-0-0-0 | dots in nucleus | N-terminal tag and C-terminal tag: nucleoplasm |
| TBP/TRF4 | Tb927.10.15950 | Q387R9 | TATA binding | other | nucleus | N-terminal tag and C-terminal tag: nucleoplasm and cytoplasm |

**Table S2. Functions and phenotypes of previously characterized candidate proteins**

| Protein names | Functions/phenotypes | References |
|---|---|---|
| BDF1-to-4 | enriched around RNAPII TSRs | Schulz et al., 2015; |
| BDF5 | essential in BF and PF cells | Alsford and Horn,2012 |
| TFIIS2-2 | binds histone peptides bearing H4K17me3 or H3K32me3 | Wang et al., 2019 |
| HAT1 | essential in BF cells; modulates telomeric silencing;<br>required for DNA replication and growth, and RNA transcription | Kawahara et al., 2008;<br>Kraus et al., 2020 |
| HAT2 | essential in BF cells; responsible for writing the H4K10ac mark found at RNAPII TSRs;<br>required for H2A.Z deposition | Kawahara et al., 2008;<br>Kraus et al., 2020 |
| HAT3 | nonessential in BF cells; acetylates H4K4 | ElBashir et al., 2015 |
| ELP3b | negatively regulates rDNA expression | Alsford and Horn, 2011 |
| DOT1A and DOT1B | H3K76 methylation; involved in trypanosome cell cycle regulation and VSG switching | Figueiredo et al., 2008;<br>Janzen et al., 2006b |
| HDAC1 | essential in BF cells; counteracts Sir2rp1 dependent telomeric silencing in BF cells | Ingram and Horn, 2002;<br>Wang et al., 2010 |
| HDAC2 | nonessential in BF cells | Ingram and Horn, 2002 |
| HDAC3 | essential in BF cells;<br>required for VSG promoter silencing throughout the developmental cycle of *T. brucei* | Ingram and Horn, 2002;<br>Wang et al., 2010 |
| HDAC4 | nonessential; knock out results in a cell cycle delay | Ingram and Horn, 2002 |
| Sir2rp1 | DNA repair and telomere-adjacent silencing of RNAPI transcription | Alsford et al., 2007 |

**Table S3. Candidate proteins with no genomic enrichment by ChIP-seq**

| Protein name | TriTryp ID | UniProt ID | Protein name | TriTryp ID | UniProt ID |
|---|---|---|---|---|---|
| PHD1 | Tb927.10.9930 | Q389Y3 | SET19 | Tb927.7.5620 | Q582H7 |
| PHD5 | Tb927.11.5870 | Q385I1 | SET20 | Tb927.8.2490 | Q57XE0 |
| HAT3 | Tb927.10.8310 | Q38AD3 | SET21 | Tb927.8.2690 | Q57XC0 |
| HAT5 | Tb927.5.2280 | Q582D7 | SET22 | Tb927.8.2710 | Q57XB8 |
| HAT6 | Tb927.1.4490 | Q4GYC9 | SET23 | Tb927.8.6470 | Q57YP8 |
| HAT7 | Tb927.10.12830 | Q388W3 | SET24 | Tb927.8.6530 | Q57TW0 |
| ELP3a | Tb927.8.5770 | Q57X05 | SET25 | Tb927.9.1510 | Q38FZ5 |
| SET1 | Tb11.v5.0422 | Q38BE1 | SET28 | Tb927.9.10100 | Q38DZ4 |
| SET3 | Tb927.1.4720 | Q4GYA6 | SET29 | Tb927.3.3370 | Q57V49 |
| SET4 | Tb927.10.11130 | Q389M6 | DOT1B | Tb927.1.570 | Q4GZF2 |
| SET5 | Tb927.10.12880 | Q388V8 | HDAC2 | Tb927.11.15600 | Q381M6 |
| SET6 | Tb927.10.3730 | Q38BM6 | HDAC4 | Tb927.5.2900 | D6XGM6 |
| SET7 | Tb927.10.4600 | - | Sir2rp1 | Tb927.7.1690 | Q57V41 |
| SET8 | Tb927.10.8060 | Q38AF8 | Sir2rp2 | Tb927.8.3140 | Q57YZ9 |
| SET9 | Tb927.10.9680 | Q38A07 | Sir2rp3 | Tb927.4.2520 | Q584D5 |
| SET10 | Tb927.11.13560 | Q382I1 | JMJ1 | Tb927.11.1760 | Q386X8 |
| SET11 | Tb927.11.5120 | Q385Q6 | JMJ2 | Tb927.11.2000 | Q386V9 |
| SET13 | Tb927.4.2440 | Q584E3 | LCM1 | Tb927.9.12780 | Q38DD6 |
| SET14 | Tb927.4.3310 | Q584A8 | CLD1 | Tb927.7.660 | Q57VR2 |
| SET15 | Tb927.5.2770 | Q57ZV3 | AGO1 | Tb927.10.10850 | Q389P5 |
| SET16 | Tb927.5.3500 | Q57UB6 | DMT | Tb927.3.1360 | Q57XW9 |
| SET17 | Tb927.6.3610 | Q585V9 | NUP110/MLP1 | Tb927.11.330 | Q387L8 |
| SET18 | Tb927.6.910 | Q585F5 | | | |

**Table S4. Overlap of tRNAs and snRNAs annotated in the assembled Lister 427 genome with TTRs and TTR-associated factors**

| Chromosome | Start position | End position | tRNA/ snRNA | ID | Overlap with TTR regions | Overlap with TTR factors |
|---|---|---|---|---|---|---|
| Chr2_core | 742379 | 742527 | snRNA | Tb427v9_000695200:snRNA (U2) | sTTR_56 | yes |
| Chr3_core | 1100184 | 1100266 | tRNA | Tb427v9_000761300:tRNA;aa=Ser;anticodon=aga | sTTR_55 | yes |
| Chr3_core | 1100470 | 1100552 | tRNA | Tb427v9_000761400:tRNA;aa=Leu;anticodon=caa | sTTR_55 | yes |
| Chr3_core | 1100611 | 1100693 | tRNA | Tb427v9_000761500:tRNA;aa=Ser;anticodon=cga | sTTR_55 | yes |
| Chr4_core | 238353 | 238424 | tRNA | Tb427v9_000825700:tRNA;aa=Pro;anticodon=cgg | no | only TBP |
| Chr4_core | 246708 | 246806 | snRNA | Tb427v9_000826000:snRNA (U6) | cTTR_69 | yes |
| Chr4_core | 246907 | 246978 | tRNA | Tb427v9_000826100:tRNA;aa=Thr;anticodon=cgt | cTTR_69 | yes |
| Chr4_core | 247045 | 247129 | tRNA | Tb427v9_000826200:tRNA;aa=Tyr;anticodon=gta | cTTR_69 | yes |
| Chr4_core | 767118 | 767191 | tRNA | Tb427v9_000846700:tRNA;aa=Val;anticodon=aac | no | yes |
| Chr4_core | 767254 | 767335 | tRNA | Tb427v9_000846800:tRNA;aa=Leu;anticodon=tag | no | yes |
| Chr4_core | 767393 | 767466 | tRNA | Tb427v9_000846900:tRNA;aa=Met;anticodon=cat | no | yes |
| Chr4_core | 780197 | 780268 | tRNA | Tb427v9_000847600:tRNA;aa=Glu;anticodon=ctc | cTTR_70 | yes |
| Chr4_core | 780356 | 780429 | tRNA | Tb427v9_000847700:tRNA;aa=Met;anticodon=cat | cTTR_70 | yes |
| Chr4_core | 780471 | 780553 | tRNA | Tb427v9_000847800:tRNA;aa=Ser;anticodon=tga | cTTR_70 | yes |
| Chr4_core | 780669 | 780742 | tRNA | Tb427v9_000847900:tRNA;aa=Val;anticodon=aac | cTTR_70 | yes |
| Chr4_core | 1212837 | 1212909 | tRNA | Tb427v9_000862100:tRNA;aa=Ser;anticodon=cga | sTTR_52 | yes |
| Chr4_core | 1212978 | 1213053 | tRNA | Tb427v9_000862200:tRNA;aa=Val;anticodon=cac | sTTR_52 | yes |
| Chr5_core | 147572 | 147644 | tRNA | Tb427v9_000901600:tRNA;aa=Arg;anticodon=acg | no | only PHD2 and TBP |
| Chr7_core | 284886 | 284957 | tRNA | Tb427v9_001104600:tRNA;aa=Gly;anticodon=gcc | no | only PHD2 and TBP |
| Chr7_core | 1885443 | 1885516 | tRNA | Tb427v9_001164700:tRNA;aa=Ala;anticodon=agc | no | yes |
| Chr7_core | 1885585 | 1885667 | tRNA | Tb427v9_001164800:tRNA;aa=Ser;anticodon=gct | no | yes |
| Chr7_core | 1885743 | 1885815 | tRNA | Tb427v9_001164900:tRNA;aa=Asp;anticodon=gtc | no | yes |
| Chr7_core | 1888962 | 1889034 | tRNA | Tb427v9_001165000:tRNA;aa=Asp;anticodon=gtc | sTTR_41 | yes |
| Chr7_core | 1889110 | 1889192 | tRNA | Tb427v9_001165100:tRNA;aa=Ser;anticodon=gct | sTTR_41 | yes |
| Chr7_core | 1889261 | 1889334 | tRNA | Tb427v9_001165200:tRNA;aa=Ala;anticodon=agc | sTTR_41 | yes |
| Chr7_core | 1905849 | 1905923 | tRNA | Tb427v9_001165500:tRNA;aa=Lys;anticodon=ctt | cTTR_48 | all except DOT1A and TRF |
| Chr7_core | 2057285 | 2057357 | tRNA | Tb427v9_001169700:tRNA;aa=Ala;anticodon=cgc | no | no |
| Chr8_core | 657296 | 657369 | tRNA | Tb427v9_001263400:tRNA;aa=Asn;anticodon=gtt | sTTR_30 | yes |
| Chr8_core | 657453 | 657526 | tRNA | Tb427v9_001263500:tRNA;aa=Trp;anticodon=cca | sTTR_30 | yes |
| Chr8_core | 717533 | 717608 | tRNA | Tb427v9_001265400:tRNA;aa=Ile;anticodon=tat | cTTR_34 | yes |

| Chromosome | Start position | End position | tRNA/ snRNA | ID | Overlap with TTR regions | Overlap with TTR factors |
|---|---|---|---|---|---|---|
| Chr8_core | 717702 | 717773 | tRNA | Tb427v9_001265500:tRNA;aa=Gln;anticodon=ttg | cTTR_34 | yes |
| Chr8_core | 717830 | 717905 | tRNA | Tb427v9_001265600:tRNA;aa=Val;anticodon=tac | cTTR_34 | yes |
| Chr8_core | 717974 | 718047 | tRNA | Tb427v9_001265700:tRNA;aa=Lys;anticodon=ctt | cTTR_34 | yes |
| Chr8_core | 721366 | 721438 | tRNA | Tb427v9_001265800:tRNA;aa=Gly;anticodon=tcc | cTTR_34 | yes |
| Chr8_core | 721524 | 721607 | tRNA | Tb427v9_001265900:tRNA;aa=Leu;anticodon=cag | cTTR_34 | yes |
| Chr8_core | 721690 | 721761 | tRNA | Tb427v9_001266000:tRNA;aa=Thr;anticodon=tgt | cTTR_34 | yes |
| Chr8_core | 721859 | 721931 | tRNA | Tb427v9_001266100:tRNA;aa=Arg;anticodon=tcg | cTTR_34 | yes |
| Chr8_core | 724624 | 724697 | tRNA | Tb427v9_001266200:tRNA;aa=Lys;anticodon=ctt | cTTR_34 | yes |
| Chr8_core | 724788 | 724861 | tRNA | Tb427v9_001266300:tRNA;aa=Arg;anticodon=acg | cTTR_34 | yes |
| Chr8_core | 916520 | 916591 | tRNA | Tb427v9_001273200:tRNA;aa=Cys;anticodon=gca | cTTR_35 | all except TFIIS2-2 |
| Chr8_core | 916732 | 916803 | tRNA | Tb427v9_001273300:tRNA;aa=Cys;anticodon=gca | cTTR_35 | all except TFIIS2-2 |
| Chr8_core | 1474462 | 1474534 | tRNA | Tb427v9_001293300:tRNA;aa=Arg;anticodon=acg | cTTR_36 | yes |
| Chr8_core | 1777934 | 1778008 | tRNA | Tb427v9_001305900:tRNA;aa=Ile;anticodon=aat | cTTR_37 | yes |
| Chr8_core | 1778055 | 1778139 | tRNA | Tb427v9_001306000:tRNA;aa=Leu;anticodon=taa | cTTR_37 | yes |
| Chr8_core | 1778232 | 1778304 | tRNA | Tb427v9_001306100:tRNA;aa=Gln;anticodon=ctg | cTTR_37 | yes |
| Chr8_core | 1778390 | 1778463 | tRNA | Tb427v9_001306200:tRNA;aa=Gln;anticodon=ctg | cTTR_37 | yes |
| Chr8_core | 1778542 | 1778616 | tRNA | Tb427v9_001306300:tRNA;aa=Ile;anticodon=aat | cTTR_37 | yes |
| Chr9_core | 236288 | 236373 | tRNA | Tb427v9_001409300:tRNA;aa=seC;anticodon=tca | no | no |
| Chr9_core | 240125 | 240210 | tRNA | Tb427v9_001409500:tRNA;aa=seC;anticodon=tca | no | no |
| Chr9_core | 1795249 | 1795319 | tRNA | Tb427v9_001488600:tRNA;aa=Gly;anticodon=gcc | cTTR_25 | all except BDF7, DOT1A and TRF |
| Chr10_core | 1888402 | 1888472 | tRNA | Tb427v9_000188600:tRNA;aa=Gly;anticodon=gcc | sTTR_13 | yes |
| Chr10_core | 1924887 | 1924958 | tRNA | Tb427v9_000190100:tRNA;aa=Pro;anticodon=agg | sTTR_14 | all except TFIIS2-2 |
| Chr10_core | 1925019 | 1925091 | tRNA | Tb427v9_000190200:tRNA;aa=Thr;anticodon=agt | sTTR_14 | all except TFIIS2-2 |
| Chr10_core | 1928536 | 1928620 | tRNA | Tb427v9_000190400:tRNA;aa=Leu;anticodon=aag | sTTR_14 | yes |
| Chr10_core | 1928681 | 1928752 | tRNA | Tb427v9_000190500:tRNA;aa=Met;anticodon=cat | sTTR_14 | yes |
| Chr10_core | 1928825 | 1928896 | tRNA | Tb427v9_000190600:tRNA;aa=Pro;anticodon=tgg | sTTR_14 | yes |
| Chr10_core | 2645688 | 2645762 | tRNA | Tb427v9_000223600:tRNA;aa=Asn;anticodon=gtt | sTTR_16 | yes |
| Chr10_core | 2645841 | 2645913 | tRNA | Tb427v9_000223700:tRNA;aa=Ala;anticodon=tgc | sTTR_16 | yes |
| Chr10_core | 2646016 | 2646089 | tRNA | Tb427v9_000223800:tRNA;aa=Arg;anticodon=tct | sTTR_16 | yes |
| Chr10_core | 2646149 | 2646221 | tRNA | Tb427v9_000223900:tRNA;aa=Lys;anticodon=ttt | sTTR_16 | yes |
| Chr10_core | 2646305 | 2646379 | tRNA | Tb427v9_000224000:tRNA;aa=Arg;anticodon=cct | sTTR_16 | yes |

| Chromosome | Start position | End position | tRNA/ snRNA | ID | Overlap with TTR regions | Overlap with TTR factors |
|---|---|---|---|---|---|---|
| Chr11_3B | 397659 | 397755 | tRNA | Tb427v9_000324300:tRNA;aa=Ile;anticodon=aat | no | no |
| Chr11_3B | 559600 | 559683 | tRNA | Tb427v9_000329900:tRNA;aa=Gly;anticodon=ccc | no | no |
| Chr11_core | 1863938 | 1864008 | tRNA | Tb427v9_000410900:tRNA;aa=Gly;anticodon=ccc | sTTR_4 | only ELP3b and TBP |
| Chr11_core | 2213880 | 2213954 | tRNA | Tb427v9_000425500:tRNA;aa=Lys;anticodon=ctt | cTTR_7 | all except DOT1A and TRF |
| Chr11_core | 3580246 | 3580317 | tRNA | Tb427v9_000484400:tRNA;aa=His;anticodon=gtg | sTTR_7 | yes |
| Chr11_core | 3580378 | 3580449 | tRNA | Tb427v9_000484500:tRNA;aa=Glu;anticodon=ttc | sTTR_7 | yes |
| Chr11_core | 3580509 | 3580581 | tRNA | Tb427v9_000484600:tRNA;aa=Phe;anticodon=gaa | sTTR_7 | yes |
| Chr11_core | 3603481 | 3603552 | tRNA | Tb427v9_000485500:tRNA;aa=Ala;anticodon=cgc | sTTR_8 | yes |
| Chr11_core | 3603618 | 3603691 | tRNA | Tb427v9_000485600:tRNA;aa=Arg;anticodon=ccg | sTTR_8 | yes |
| Chr11_core | 3603801 | 3603873 | tRNA | Tb427v9_000485700:tRNA;aa=Phe;anticodon=gaa | sTTR_8 | yes |

**Table S5. Significantly enriched interactors detected by proteomic analysis of affinity selected YFP-tagged proteins**

| CRD1 IP significantly enriched interactors (over untagged control) | | | | |
|---|---|---|---|---|
| **UniProt ID** | **TriTryp ID** | **Description** | **-$\log_{10}$ Student's t-test p-value** | **$\log_2$ (tagged/untagged)** |
| Q4GYF3 | Tb927.1.4250 | hypothetical protein | 5.53089 | 11.4431 |
| Q38D80 | Tb927.9.13470 | SET27 | 5.94395 | 11.4397 |
| Q57X70 | Tb11.v5.0267 | CRD1 | 4.08625 | 9.18821 |
| Q382F5 | Tb927.11.13820 | hypothetical protein | 6.65416 | 9.17537 |
| Q582U8 | Tb927.3.2350 | hypothetical protein | 7.10211 | 9.04339 |
| Q382Z7 | Tb927.11.11840 | hypothetical protein | 6.06473 | 8.75264 |
| D6XIZ6 | Tb927.7.4650 | J Biosynthesis Protein 2 (JBP2) | 6.33932 | 7.9307 |
| Q38FV1 | Tb927.9.2070 | hypothetical protein | 2.09326 | 6.95152 |
| Q38BC1 | Tb927.10.4800 | hypothetical protein | 2.70589 | 4.47099 |
| Q38BS4 | Tb927.10.3220 | nucleolus protein | 4.25958 | 4.04605 |
| Q586X9 | Tb927.2.3580 | transcription elongation factor s-II (TFIIS2-1) | 2.30696 | 3.65927 |
| Q57W25 | Tb927.5.800 | casein kinase I, isoform 2 | 5.87744 | 2.9952 |
| Q387L4 | Tb927.11.370 | repressor activator protein 1 (RAP1) | 3.04907 | 2.19994 |
| Q580S5 | Tb927.3.5500 | DNA-directed RNA polymerase II subunit 3 (RPB3) | 2.41931 | 2.06975 |
| Q57YU2 | Tb927.8.7400 | RNA polymerase IIA largest subunit (RPB1) | 2.92416 | 2.01862 |

### SET27 IP significantly enriched interactors (over untagged control)

| UniProt ID | TriTryp ID | Description | -$log_{10}$ Student's t-test p-value | $log_2$ (tagged/untagged) |
|---|---|---|---|---|
| Q4GYF3 | Tb927.1.4250 | hypothetical protein | 5.32451 | 11.5549 |
| Q38D80 | Tb927.9.13470 | SET27 | 4.15023 | 11.3024 |
| Q57X70 | Tb11.v5.0267 | CRD1 | 4.41402 | 10.8901 |
| Q382F5 | Tb927.11.13820 | hypothetical protein | 4.76568 | 9.39333 |
| Q382Z7 | Tb927.11.11840 | hypothetical protein | 5.30182 | 9.15674 |
| D6XIZ6 | Tb927.7.4650 | J Biosynthesis Protein 2 (JBP2) | 5.09272 | 9.10164 |
| Q582U8 | Tb927.3.2350 | hypothetical protein | 4.70479 | 8.88102 |
| Q38BT9 | Tb927.10.3070 | hypothetical protein | 3.00745 | 4.65672 |
| Q57Z72 | Tb927.5.3210 | small ubiquitin-related modifier | 2.2191 | 4.57882 |
| Q38EP6 | Tb927.9.6920 | hypothetical protein | 2.10149 | 4.34594 |
| Q38BC1 | Tb927.10.4800 | hypothetical protein | 2.71533 | 4.25565 |
| Q586X9 | Tb927.2.3580 | transcription elongation factor s-II (TFIIS2-1) | 2.21306 | 4.05266 |
| Q38BS4 | Tb927.10.3220 | nucleolus protein | 2.90768 | 4.0165 |
| P86938 | Tb927.11.13640 | J-binding protein 1 (JBP1) | 3.18784 | 3.5602 |
| Q57W25 | Tb927.5.800 | casein kinase I, isoform 2 | 5.83603 | 3.04821 |
| Q389E6 | Tb927.10.11960 | hypothetical protein | 2.37392 | 2.98484 |
| Q57W24 | Tb927.5.790 | casein kinase I isoform 1 | 3.60939 | 2.82452 |
| Q57VC7 | Tb927.5.1360 | nucleoside 2-deoxyribosyltransferase | 3.59736 | 2.56284 |
| Q586H7 | Tb927.6.5050 | V-type ATPase, C subunit | 3.63701 | 2.53399 |
| Q388E9 | Tb927.10.14530 | 19S proteasome non-atpase subunit 8 | 2.16045 | 2.44652 |
| Q57UB5 | Tb927.5.3510 | structural maintenance of chromosome 3 (SMC3) | 3.09014 | 2.17858 |
| Q38E40 | Tb927.9.9550 | hypothetical protein | 3.18533 | 2.14501 |
| Q38C86 | Tb927.10.1550 | proteasome regulatory non-ATP-ase subunit 5 | 4.10981 | 2.00096 |

### BDF1 IP significantly enriched interactors (over untagged control)

| UniProt ID | TriTryp ID | Description | $-\log_{10}$ Student's t-test p-value | $\log_2$ (tagged/untagged) |
|---|---|---|---|---|
| Q38AE9 | Tb927.10.8150 | BDF1 | 4.93793 | 9.72633 |
| Q57UR8 | Tb927.7.4380 | BDF4 | 4.54907 | 7.93766 |
| Q389A2 | Tb927.10.12410 | hypothetical protein, conserved | 2.98317 | 3.23467 |
| Q383J7 | Tb927.11.10820 | Casein kinase II subunit beta | 2.73998 | 2.51666 |
| Q57WL5 | Tb927.8.8150 | C2 domain containing protein | 2.40712 | 2.18084 |

### BDF4 IP significantly enriched interactors (over untagged control)

| UniProt ID | TriTryp ID | Description | $-\log_{10}$ Student's t-test p-value | $\log_2$ (tagged/untagged) |
|---|---|---|---|---|
| Q38AE9 | Tb927.10.8150 | BDF1 | 4.188294496 | 10.46347491 |
| Q57UR8 | Tb927.7.4380 | BDF4 | 4.899856567 | 9.580053965 |
| Q38D21 | Tb927.9.14430 | Casein kinase II | 2.61634521 | 4.563868841 |
| Q383S2 | Tb927.11.10070 | BDF3 | 2.321038534 | 3.810703278 |
| Q57YY8 | Tb927.8.3250 | dynein heavy chain | 2.187183878 | 3.700003306 |
| Q57V45 | Tb927.3.3410 | aspartyl aminopeptidase | 2.343303395 | 3.043106079 |
| Q383J7 | Tb927.11.10820 | Casein kinase II subunit beta | 3.324763121 | 2.964817047 |
| Q586Q9 | Tb927.2.4580 | UNC119 | 3.52554182 | 2.820368449 |
| Q57VC7 | Tb927.5.1360 | nucleoside 2-deoxyribosyltransferase | 2.708928634 | 2.726233164 |
| Q38D92 | Tb927.9.13320 | hypothetical protein | 2.267759854 | 2.656667074 |
| Q382J7 | Tb927.11.13400 | BDF5 | 2.3335 | 2.2561 |
| Q57UC1 | Tb927.5.3450 | eukaryotic translation initiation factor eIF2A | 2.15945 | 2.01132 |

### BDF3 IP significantly enriched interactors (over untagged control)

| UniProt ID | TriTryp ID | Description | $-log_{10}$ Student's t-test p-value | $log_2$ (tagged/untagged) |
|---|---|---|---|---|
| Q385P5 | Tb927.11.5230 | hypothetical protein, conserved | 4.60069 | 15.7219 |
| Q383S2 | Tb927.11.10070 | BDF3 | 5.64379 | 15.3806 |
| Q584F3 | Tb927.4.2340 | hypothetical protein, conserved | 7.46251 | 14.3231 |
| Q38D92 | Tb927.9.13320 | hypothetical protein, conserved | 4.32584 | 14.1699 |
| Q382J7 | Tb927.11.13400 | BDF5 | 4.65476 | 13.3481 |
| Q57Y87 | Tb927.7.2770 | hypothetical protein, conserved | 4.90171 | 11.7022 |
| Q580M8 | Tb927.3.4140 | hypothetical protein, conserved | 4.96735 | 11.4365 |
| Q585E0 | Tb927.6.1070 | hypothetical protein, conserved | 7.05768 | 11.3361 |
| Q383C5 | Tb927.11.11530 | HAT2 | 4.98438 | 11.199 |
| Q4FKD8 | Tb927.11.18680 | dynein light chain LC8 | 4.38862 | 7.06557 |
| Q586Q9 | Tb927.2.4580 | UNC119 | 3.75556 | 4.53563 |
| Q57Z72 | Tb927.5.3210 | small ubiquitin-related modifier | 3.50341 | 4.12536 |
| Q38BR9 | Tb927.10.3280 | 60S ribosomal proteins L38 | 4.13696 | 3.68518 |
| Q383E9 | Tb927.11.11290 | heat shock protein 70 | 2.54297 | 3.16894 |
| Q4GYZ8 | Tb927.1.2230 | small myristoylated protein 1-1 | 5.00505 | 2.87762 |

### BDF5 IP significantly enriched interactors (over untagged control)

| UniProt ID | TriTryp ID | Description | $-log_{10}$ Student's t-test p-value | $log_2$ (tagged/untagged) |
|---|---|---|---|---|
| Q383S2 | Tb927.11.10070 | BDF3 | 5.03137 | 14.4394 |
| Q385P5 | Tb927.11.5230 | hypothetical protein, conserved | 4.16393 | 14.2577 |
| Q38D92 | Tb927.9.13320 | hypothetical protein, conserved | 4.03255 | 13.5657 |
| Q584F3 | Tb927.4.2340 | hypothetical protein, conserved | 4.76342 | 13.3877 |
| Q382J7 | Tb927.11.13400 | BDF5 | 4.64404 | 13.0329 |
| Q57Y87 | Tb927.7.2770 | hypothetical protein, conserved | 5.38435 | 11.5875 |
| Q585E0 | Tb927.6.1070 | hypothetical protein, conserved | 5.26236 | 11.0428 |
| Q580M8 | Tb927.3.4140 | hypothetical protein, conserved | 4.98517 | 11.001 |
| Q383C5 | Tb927.11.11530 | HAT2 | 4.83268 | 10.0504 |
| Q4FKD8 | Tb927.11.18680 | dynein light chain LC8 | 4.21709 | 6.66445 |
| Q586Q9 | Tb927.2.4580 | UNC119 | 3.36202 | 4.49549 |
| Q38BR9 | Tb927.10.3280 | 60S ribosomal proteins L38 | 3.73772 | 4.04956 |
| Q57Z72 | Tb927.5.3210 | small ubiquitin-related modifier | 3.2482 | 2.9886 |
| Q4GYZ8 | Tb927.1.2230 | small myristoylated protein 1-1 | 2.54476 | 2.80002 |
| Q383E9 | Tb927.11.11290 | heat shock protein 70 | 2.77564 | 2.57324 |

### HAT2 IP significantly enriched interactors (over untagged control)

| UniProt ID | TriTryp ID | Description | $-log_{10}$ Student's t-test p-value | $log_2$ (tagged/untagged) |
|---|---|---|---|---|
| Q385P5 | Tb927.11.5230 | hypothetical protein, conserved | 6.54066 | 12.2937 |
| Q38D92 | Tb927.9.13320 | hypothetical protein, conserved | 5.66256 | 12.1545 |
| Q383S2 | Tb927.11.10070 | BDF3 | 5.78366 | 12.0736 |
| Q383C5 | Tb927.11.11530 | HAT2 | 4.7375 | 11.497 |
| Q382J7 | Tb927.11.13400 | BDF5 | 5.32275 | 11.381 |
| Q584F3 | Tb927.4.2340 | hypothetical protein, conserved | 3.48604 | 11.3471 |
| Q585E0 | Tb927.6.1070 | hypothetical protein, conserved | 6.59864 | 10.8439 |
| Q57Y87 | Tb927.7.2770 | hypothetical protein, conserved | 4.81634 | 10.5086 |
| Q580M8 | Tb927.3.4140 | hypothetical protein, conserved | 5.08953 | 9.86611 |
| Q4FKD8 | Tb927.11.18680 | dynein light chain LC8 | 4.15852 | 7.369 |
| Q57Z72 | Tb927.5.3210 | small ubiquitin-related modifier | 2.76433 | 5.35268 |
| Q38BI9 | Tb927.10.4110 | 60S ribosomal protein L30 | 2.42707 | 3.23005 |
| Q9N942 | Tb927.1.420 | retrotransposon hot spot protein 5 (RHS5) | 3.10308 | 3.01725 |
| Q38BE4 | Tb927.10.4570 | translation elongation factor 2 | 2.94249 | 2.17828 |

### BDF2 IP significantly enriched interactors (over untagged control)

| UniProt ID | TriTryp ID | Description | $-\log_{10}$ Student's t-test p-value | $\log_2$ (tagged/untagged) |
|---|---|---|---|---|
| Q38AM1 | Tb927.10.7420 | BDF2 | 4.94142 | 13.812 |
| Q586J9 | Tb927.2.2190 | HDAC3 | 5.48101 | 12.925 |
| Q582M3 | Tb927.7.6360 | Histone H2A.Z | 5.5163 | 8.77685 |
| Q384T0 | Tb927.11.7350 | Histone H2B.V | 5.64211 | 8.65261 |
| Q38E38 | Tb927.9.9580 | tubulin tyrosine ligase protein | 4.66156 | 7.7985 |
| Q582V9 | Tb927.3.2460 | hypothetical protein | 4.9989 | 7.06729 |
| Q383U2 | Tb927.11.9870 | Telomere-associated protein 1 (TelAP1) | 4.1549 | 6.97577 |
| Q586Z9 | Tb927.6.4330 | hypothetical protein | 3.34214 | 6.9275 |
| Q388W1 | Tb927.10.12850 | ttaggg binding factor (TRF) | 5.35967 | 6.55814 |
| Q57XY9 | Tb927.3.1560 | TRF-Interacting Factor 2 (TIF2) | 4.40537 | 6.3738 |
| Q38FD7 | Tb927.9.4000 | hypothetical protein | 2.89011 | 5.81918 |
| Q586Q9 | Tb927.2.4580 | UNC119 | 4.33193 | 5.70718 |
| Q57UN4 | Tb927.7.4040 | hypothetical protein | 3.53327 | 4.9661 |
| Q385L3 | Tb927.11.5550 | DNA polymerase theta (POLQ) | 3.27198 | 4.95997 |
| Q57ZL0 | Tb927.3.3020 | actin-like protein | 5.7851 | 4.94816 |
| Q584P9 | Tb927.6.2570 | SUMO-interacting motif-containing protein | 4.62419 | 4.90393 |
| Q383K6 | Tb927.11.10730 | SWI/SNF-related helicase | 3.65983 | 4.76426 |
| Q389H2 | Tb927.10.11690 | YEATS family | 3.27706 | 4.27104 |
| Q38EB9 | Tb927.9.8520 | hypothetical protein | 2.92994 | 4.2494 |
| Q57YH0 | Tb927.8.600 | Bucentaur or craniofacial development | 2.94405 | 3.88251 |
| Q389D3 | Tb927.10.12100 | RNA-binding protein 7B | 2.71219 | 3.49313 |
| Q385T8 | Tb927.11.4800 | clathrin coat assembly protein | 3.80316 | 3.47689 |
| Q38BZ3 | Tb927.10.2520 | PrimPol-like protein 2 | 2.73316 | 3.47355 |
| Q583J3 | Tb927.4.2000 | ruvB-like DNA helicase | 3.61615 | 3.41918 |
| Q580C6 | Tb927.4.980 | actin | 3.02507 | 3.32672 |
| Q38C43 | Tb927.10.2000 | actin-like protein | 2.37492 | 3.31999 |
| Q57WS6 | Tb927.7.1240 | Sphingosine kinase | 2.11279 | 3.29644 |
| Q581V4 | Tb927.4.1270 | ruvB-like DNA helicase | 3.29543 | 3.20032 |
| Q385E0 | Tb927.11.6290 | HIT zinc finger | 2.49815 | 3.0698 |
| Q57YW0 | Tb927.8.3530 | glycerol-3-phosphate dehydrogenase [NAD+], glycosomal | 5.18765 | 2.94186 |

### BDF2 IP significantly enriched interactors (over untagged control) - continued

| UniProt ID | TriTryp ID | Description | $-log_{10}$ Student's t-test p-value | $log_2$ (tagged/untagged) |
|---|---|---|---|---|
| Q389G9 | Tb927.10.11720 | ZCW1 | 2.89967 | 2.89912 |
| Q57ZS8 | Tb927.5.2090 | kinesin | 3.26358 | 2.74008 |
| Q57W25 | Tb927.5.800 | casein kinase I, isoform 2 | 3.7468 | 2.69126 |
| Q389D9 | Tb927.10.12030 | hypothetical protein | 3.67541 | 2.67071 |
| Q384I8 | Tb927.11.8310 | class I transcription factor A, subunit 4 (CITFA-4) | 2.451 | 2.64382 |
| Q385A8 | Tb927.11.6610 | hypothetical protein | 2.16774 | 2.44445 |
| Q387B3 | Tb927.11.1390 | class I transcription factor A, subunit 1 (CITFA-1) | 3.77834 | 2.28532 |
| Q38DS7 | Tb927.9.10960 | ATP-dependent DEAD/H RNA helicase | 4.30734 | 2.26913 |
| Q57XM9 | Tb927.7.2450 | SUMO-interacting motif-containing protein | 2.87997 | 2.2086 |
| Q38D76 | Tb927.9.13510 | Poly(A)-specific ribonuclease PARN-3 | 2.66067 | 2.00854 |

### HDAC3 IP significantly enriched interactors (over untagged control)

| UniProt ID | TriTryp ID | Description | $-\log_{10}$ Student's t-test p-value | $\log_2$ (tagged/untagged) |
|---|---|---|---|---|
| Q586J9 | Tb927.2.2190 | HDAC3 | 5.46467 | 12.9502 |
| Q57XY9 | Tb927.3.1560 | TRF-Interacting Factor 2 (TIF2) | 5.1545 | 8.95426 |
| Q586Z9 | Tb927.6.4330 | hypothetical protein | 4.54883 | 8.81523 |
| Q38AM1 | Tb927.10.7420 | BDF2 | 4.99548 | 8.60155 |
| Q388W1 | Tb927.10.12850 | ttaggg binding factor (TRF) | 4.01868 | 8.55698 |
| Q383U2 | Tb927.11.9870 | Telomere-associated protein 1 (TelAP1) | 3.9975 | 8.24039 |
| Q38EB9 | Tb927.9.8520 | hypothetical protein | 3.65687 | 7.71109 |
| Q385L3 | Tb927.11.5550 | DNA polymerase theta (POLQ) | 4.17948 | 7.53688 |
| Q582M3 | Tb927.7.6360 | Histone H2A.Z | 4.33192 | 7.20786 |
| Q582V9 | Tb927.3.2460 | hypothetical protein | 3.9418 | 7.12454 |
| Q38FD7 | Tb927.9.4000 | hypothetical protein | 3.40403 | 7.00728 |
| Q38BZ3 | Tb927.10.2520 | PrimPol-like protein 2 | 4.02346 | 6.39412 |
| Q384T0 | Tb927.11.7350 | Histone H2B.V | 4.86712 | 6.21943 |
| Q387L4 | Tb927.11.370 | repressor activator protein 1 (RAP1) | 3.94259 | 5.0573 |
| Q586Q9 | Tb927.2.4580 | UNC119 | 3.48225 | 4.21712 |
| Q57UB5 | Tb927.5.3510 | structural maintenance of chromosome 3 (SMC3) | 3.94742 | 4.09873 |
| Q57W25 | Tb927.5.800 | casein kinase I, isoform 2 | 4.09069 | 4.08785 |
| Q580Q4 | Tb927.4.5310 | Repressor of differentiation kinase 2 | 3.04617 | 3.66189 |
| Q388S9 | Tb927.10.13180 | Nrap protein | 2.48075 | 3.65437 |
| Q4GZ82 | Tb927.1.1370 | rRNA biogenesis protein | 2.54895 | 3.52187 |
| Q38D76 | Tb927.9.13510 | Poly(A)-specific ribonuclease PARN-3 | 3.07138 | 3.34021 |
| Q580T1 | Tb927.3.5440 | SNF2 DNA repair protein | 2.29208 | 3.18214 |
| Q38DK9 | Tb927.9.11850 | structural maintenance of chromosome 1 (SMC1) | 2.34857 | 2.96831 |
| Q385E2 | Tb927.11.6270 | inositol-1,4,5-trisphosphate (IP3) 5-phosphatase | 2.02254 | 2.94661 |
| Q57WS6 | Tb927.7.1240 | Sphingosine kinase | 3.01619 | 2.87586 |
| Q385P5 | Tb927.11.5230 | hypothetical protein | 3.09052 | 2.2089 |
| Q38DH5 | Tb927.9.12300 | replication factor C, subunit 3 | 2.75055 | 2.00741 |

### SET26 IP significantly enriched interactors (over untagged control)

| UniProt ID | TriTryp ID | Description | $-\log_{10}$ Student's t-test p-value | $\log_2$ (tagged/untagged) |
|---|---|---|---|---|
| Q389G9 | Tb927.10.11720 | ZCW1 | 7.02945 | 12.2049 |
| Q38AF4 | Tb927.10.8100 | SET26 | 4.22372 | 9.24843 |
| Q384T0 | Tb927.11.7350 | Histone H2B.V | 2.89069 | 8.17785 |
| Q582M3 | Tb927.7.6360 | Histone H2A.Z | 4.46897 | 6.16313 |
| Q580R3 | Tb927.3.5620 | FACT complex subunit SPT16 | 3.63648 | 5.89062 |
| Q385D4 | Tb927.11.6350 | AAA ATPase (BDF7) | 3.91991 | 5.78368 |
| Q388G3 | Tb927.10.14390 | FACT complex subunit POB3 | 3.37037 | 5.22017 |
| Q387X7 | Tb927.10.15350 | Histone H3.V | 3.6525 | 5.00264 |
| Q57Z31 | Tb927.5.4170 | Histone H4 | 3.36077 | 4.61553 |
| Q383E9 | Tb927.11.11290 | heat shock protein 70 | 5.03661 | 4.61157 |
| Q384B5 | Tb927.11.9130 | Replication factor A protein 1 | 3.37426 | 4.60043 |
| Q57YA3 | Tb927.7.2820 | Histone H2A | 4.0639 | 4.55846 |
| Q386Y4 | Tb927.11.1700 | hypothetical protein | 2.88207 | 4.39468 |
| Q583H5 | Tb927.4.3810 | DNA-directed RNA polymerase II subunit 2 (RPB2) | 2.33135 | 4.08113 |
| Q386P3 | Tb927.11.2670 | Nucleoporin NUP59 | 3.46948 | 4.04731 |
| Q38D92 | Tb927.9.13320 | hypothetical protein | 2.47669 | 3.66993 |
| Q389T1 | Tb927.10.10590 | Histone H2B | 4.31282 | 3.59015 |
| Q583S0 | Tb927.6.2640 | importin alpha subunit | 2.30335 | 3.56287 |
| Q38AE0 | Tb927.10.8240 | hypothetical protein | 2.78086 | 3.54509 |
| Q580N2 | Tb927.3.4100 | Pyruvate transporter | 4.27969 | 3.47669 |
| Q38A77 | Tb927.10.8970 | kinetoplast-associated protein 4 isoform 2 | 3.85018 | 3.44607 |
| Q38E19 | Tb927.9.9810 | hypothetical protein | 3.20778 | 3.29031 |
| Q57YU2 | Tb927.8.7400 | RNA polymerase IIA largest subunit (RPB1) | 2.15392 | 3.152 |
| Q385C3 | Tb927.11.6460 | hypothetical protein | 3.31061 | 3.00282 |
| Q38FW0 | Tb927.9.1980 | hypothetical protein | 3.74352 | 3.00098 |
| Q38D21 | Tb927.9.14430 | Casein kinase II | 4.09447 | 2.94132 |
| Q57WL5 | Tb927.8.8150 | C2 domain containing protein | 4.11178 | 2.93721 |
| Q4GYX7 | Tb927.1.2430 | Histone H3 | 2.39318 | 2.84673 |
| Q57ZS7 | Tb927.5.2080 | GMP reductase | 2.82782 | 2.811 |
| Q387Q3 | Tb927.10.16120 | inosine-5'-monophosphate dehydrogenase | 3.08141 | 2.80256 |

### SET26 IP significantly enriched interactors (over untagged control) - continued

| UniProt ID | TriTryp ID | Description | $-log_{10}$ Student's t-test p-value | $log_2$ (tagged/untagged) |
|---|---|---|---|---|
| Q388C7 | Tb927.10.14770 | Associated kinase of Tb14-3-3 | 2.07295 | 2.61634 |
| Q388K3 | Tb927.10.13960 | paralyzed flagella protein 20 | 2.60087 | 2.52675 |
| Q389A2 | Tb927.10.12410 | hypothetical protein | 3.01927 | 2.48542 |
| Q38DT1 | Tb927.9.10920 | kinetoplastid kinetochore protein 3 (KKT3) | 2.78001 | 2.45683 |
| Q584R4 | Tb927.6.2420 | p22 protein precursor | 3.74551 | 2.4026 |
| Q386D5 | Tb927.11.3770 | Dpy-30 motif containing protein | 2.77261 | 2.29603 |
| Q381T5 | Tb927.11.15000 | survival of motor neuron (SMN)-like protein | 2.214 | 2.23543 |
| Q38B98 | Tb927.10.5030 | ubiquitin/ribosomal protein S27a | 3.55043 | 2.16014 |
| Q580S5 | Tb927.3.5500 | DNA-directed RNA polymerase II subunit 3 (RPB3) | 3.71295 | 2.12249 |
| Q582W2 | Tb927.3.2490 | hypothetical protein | 2.03244 | 2.09296 |
| Q57X47 | Tb927.5.4570 | Flagellum adhesion protein 3 | 2.25438 | 2.09208 |
| Q57YE9 | Tb927.8.820 | VID27 cytoplasmic protein | 2.57063 | 2.0567 |

### ZCW1 IP significantly enriched interactors (over untagged control)

| UniProt ID | TriTryp ID | Description | $-log_{10}$ Student's t-test p-value | $log_2$ (tagged/untagged) |
|---|---|---|---|---|
| Q57UN4 | Tb927.7.4040 | hypothetical protein | 5.00494 | 11.7469 |
| Q383K6 | Tb927.11.10730 | SWI/SNF-related helicase | 6.66968 | 11.6905 |
| Q580C6 | Tb927.4.980 | actin | 6.24716 | 11.6278 |
| Q389H2 | Tb927.10.11690 | YEATS family | 4.11583 | 11.561 |
| Q57ZL0 | Tb927.3.3020 | actin-like protein | 4.47839 | 11.3822 |
| Q584P9 | Tb927.6.2570 | SUMO-interacting motif-containing protein | 3.71735 | 10.8057 |
| Q38C43 | Tb927.10.2000 | actin-like protein | 4.04326 | 10.1429 |
| Q385E0 | Tb927.11.6290 | HIT zinc finger | 5.72496 | 10.0378 |
| Q389G9 | Tb927.10.11720 | ZCW1 | 5.52659 | 9.89842 |
| Q57YH0 | Tb927.8.600 | Bucentaur or craniofacial development | 4.74135 | 9.8613 |
| Q583J3 | Tb927.4.2000 | ruvB-like DNA helicase | 5.70316 | 9.40665 |
| Q38AF4 | Tb927.10.8100 | SET26 | 4.38096 | 8.99975 |
| Q385I5 | Tb927.11.5830 | YL1 nuclear protein | 4.51071 | 8.95025 |
| Q581V4 | Tb927.4.1270 | ruvB-like DNA helicase | 6.30647 | 8.89021 |
| Q385D4 | Tb927.11.6350 | AAA ATPase (BDF7) | 3.83483 | 6.00911 |
| Q384T0 | Tb927.11.7350 | Histone H2B.V | 4.47655 | 5.79935 |
| Q582M3 | Tb927.7.6360 | Histone H2A.Z | 5.32221 | 5.55756 |
| Q388G3 | Tb927.10.14390 | FACT complex subunit POB3 | 4.00616 | 5.16272 |
| Q387D1 | Tb927.11.1210 | Domain of unknown function (DUF4470) | 4.0592 | 4.56921 |
| Q583S0 | Tb927.6.2640 | importin alpha subunit | 2.95249 | 4.54393 |
| Q383E9 | Tb927.11.11290 | heat shock protein 70 | 5.58338 | 4.41933 |
| Q386P3 | Tb927.11.2670 | Nucleoporin NUP59 | 2.14711 | 3.80024 |
| Q580R3 | Tb927.3.5620 | FACT complex subunit SPT16 | 3.61713 | 3.74787 |
| Q38AM1 | Tb927.10.7420 | BDF2 | 2.9484 | 3.65793 |
| Q38D21 | Tb927.9.14430 | Casein kinase II | 2.69264 | 3.60069 |
| Q385P5 | Tb927.11.5230 | hypothetical protein protein | 3.28779 | 3.24957 |
| Q57XG4 | Tb927.8.2250 | Fungal tRNA ligase phosphodiesterase domain | 2.20582 | 3.13698 |
| Q386Y7 | Tb927.11.1670 | cysteine desulfurase | 3.33443 | 3.08449 |
| Q57Z31 | Tb927.5.4170 | Histone H4 | 3.9249 | 3.01404 |
| Q57YA3 | Tb927.7.2820 | Histone H2A | 3.46827 | 2.92822 |

### ZCW1 IP significantly enriched interactors (over untagged control) - continued

| UniProt ID | TriTryp ID | Description | $-\log_{10}$ Student's t-test p-value | $\log_2$ (tagged/untagged) |
|---|---|---|---|---|
| Q386Y4 | Tb927.11.1700 | hypothetical protein | 3.49597 | 2.77325 |
| Q4GYX7 | Tb927.1.2430 | Histone H3 | 4.12096 | 2.61517 |
| Q4GYK7 | Tb927.1.3670 | expression site-associated gene 8 (ESAG8) protein | 2.35331 | 2.57668 |
| Q389T1 | Tb927.10.10590 | Histone H2B | 3.33103 | 2.55487 |
| Q586Q9 | Tb927.2.4580 | UNC119 | 2.50076 | 2.23221 |
| Q382J7 | Tb927.11.13400 | BDF5 | 2.29002 | 2.11787 |
| Q387T9 | Tb927.10.15750 | Tripartite attachment complex protein 197 | 3.57696 | 2.07161 |
| Q57ZX9 | Tb927.5.3030 | Intraflagellar transport protein 121 | 2.63744 | 2.74248 |

### BDF6 IP significantly enriched interactors (over untagged control)

| UniProt ID | TriTryp ID | Description | $-\log_{10}$ Student's t-test p-value | $\log_2$ (tagged/untagged) |
|---|---|---|---|---|
| Q57ZF8 | Tb927.7.5310 | YEATS family | 6.11969 | 12.5739 |
| Q4GYN3 | Tb927.1.3400 | BDF6 | 5.6584 | 12.1121 |
| Q388I3 | Tb927.10.14190 | hypothetical protein, conserved | 5.54719 | 11.8692 |
| Q4GZG2 | Tb927.1.650 | conserved protein, unknown function | 5.39763 | 11.6205 |
| Q57UK4 | Tb927.8.5320 | hypothetical protein, conserved | 4.60683 | 11.1868 |
| D6XIY7 | Tb927.7.4560 | HAT1 | 5.87644 | 10.7995 |
| Q38FN2 | Tb927.9.2910 | EAF6 (NuA4 complex component) | 3.68363 | 9.94123 |
| Q386G8 | Tb927.11.3430 | hypothetical protein, conserved | 4.58029 | 8.80534 |
| Q585C3 | Tb927.6.1240 | hypothetical protein, conserved | 5.19258 | 7.78062 |
| Q57UX1 | Tb927.8.4810 | prohibitin 1 | 2.86836 | 2.39806 |
| Q57Z72 | Tb927.5.3210 | small ubiquitin-related modifier | 3.24902 | 2.05029 |

### HAT1 IP significantly enriched interactors (over untagged control)

| UniProt ID | TriTryp ID | Description | $-\log_{10}$ Student's t-test p-value | $\log_2$ (tagged/untagged) |
|---|---|---|---|---|
| Q57ZF8 | Tb927.7.5310 | YEATS family | 5.21336 | 12.6562 |
| Q388I3 | Tb927.10.14190 | hypothetical protein, conserved | 4.12823 | 12.0313 |
| Q57UK4 | Tb927.8.5320 | hypothetical protein, conserved | 5.21298 | 11.3855 |
| Q4GZG2 | Tb927.1.650 | conserved protein, unknown function | 4.49989 | 11.2084 |
| Q4GYN3 | Tb927.1.3400 | BDF6 | 5.49118 | 11.1912 |
| Q386G8 | Tb927.11.3430 | hypothetical protein, conserved | 6.13574 | 10.9019 |
| D6XIY7 | Tb927.7.4560 | HAT1 | 4.97472 | 10.4031 |
| Q38FN2 | Tb927.9.2910 | EAF6 (NuA4 complex component) | 4.74772 | 10.1789 |
| Q585C3 | Tb927.6.1240 | hypothetical protein, conserved | 5.64965 | 7.83824 |
| Q383W5 | Tb927.11.9640 | glycyl-tRNA synthetase | 3.96282 | 2.34455 |
| Q586Q9 | Tb927.2.4580 | UNC119 | 2.93824 | 2.33758 |
| Q389D8 | Tb927.10.12040 | mitogen-activated protein kinase 11 | 3.69224 | 2.23945 |

### EAF6 IP significantly enriched interactors (over untagged control)

| UniProt ID | TriTryp ID | Description | $-\log_{10}$ Student's t-test p-value | $\log_2$ (tagged/untagged) |
|---|---|---|---|---|
| Q388I3 | Tb927.10.14190 | hypothetical protein, conserved | 5.6127 | 12.6188 |
| Q57ZF8 | Tb927.7.5310 | YEATS family | 5.31752 | 12.5041 |
| D6XIY7 | Tb927.7.4560 | HAT1 | 4.77593 | 11.562 |
| Q4GYN3 | Tb927.1.3400 | BDF6 | 4.64662 | 11.43 |
| Q4GZG2 | Tb927.1.650 | conserved protein, unknown function | 5.23768 | 10.8304 |
| Q386G8 | Tb927.11.3430 | hypothetical protein, conserved | 4.27064 | 10.5847 |
| Q57UK4 | Tb927.8.5320 | hypothetical protein, conserved | 4.60791 | 10.5651 |
| Q585C3 | Tb927.6.1240 | hypothetical protein, conserved | 3.91776 | 9.07108 |
| Q389Y3 | Tb927.10.9930 | PHD1 | 4.89452 | 8.76258 |
| Q38FN2 | Tb927.9.2910 | EAF6 (NuA4 complex component) | 3.44309 | 8.58075 |
| Q384N0 | Tb927.11.7880 | hypothetical protein, conserved | 3.45527 | 6.17389 |
| Q38AD3 | Tb927.10.8310 | HAT3 | 2.86916 | 5.27809 |
| Q38C15 | Tb927.10.2290 | chaperone protein DnaJ | 4.44281 | 5.13168 |
| Q586Q9 | Tb927.2.4580 | UNC119 | 2.53018 | 4.21007 |
| Q57V04 | Tb927.7.2070 | heat shock protein DNAJ | 3.00472 | 4.10741 |
| Q38DC5 | Tb927.9.12900 | RNA polymerase-associated protein LEO1 | 2.72506 | 2.77654 |
| Q586X9 | Tb927.2.3580 | transcription elongation factor s-II (TFIIS2-1) | 2.50704 | 2.56127 |
| D6XHJ3 | Tb927.6.1090 | proteasome regulatory ATPase subunit 3 | 2.48845 | 2.47009 |
| Q383D0 | Tb927.11.11480 | Trichohyalin | 2.66667 | 2.22829 |
| Q57Y42 | Tb927.8.7980 | Pyrophosphate-energized vacuolar membrane proton pump 2 | 2.4109 | -2.87184 |

### HDAC1 IP significantly enriched interactors (over untagged control)

| UniProt ID | TriTryp ID | Description | $-log_{10}$ Student's t-test p-value | $log_2$ (tagged/untagged) |
|---|---|---|---|---|
| Q38FV1 | Tb927.9.2070 | hypothetical protein | 6.05488 | 11.7449 |
| Q57TZ5 | Tb927.7.1650 | hypothetical protein | 5.39599 | 11.4167 |
| Q584W0 | Tb927.6.3170 | hypothetical protein | 4.1002 | 10.7475 |
| Q57WX2 | Tb927.3.890 | hypothetical protein | 4.84634 | 9.67638 |
| Q38C74 | Tb927.10.1680 | HDAC1 | 4.00622 | 7.22132 |
| Q585H7 | Tb927.4.3730 | hypothetical protein | 4.26412 | 6.34073 |

### BDF7 IP significantly enriched interactors (over untagged control)

| UniProt ID | TriTryp ID | Description | $-\log_{10}$ Student's t-test p-value | $\log_2$ (tagged/untagged) |
|---|---|---|---|---|
| Q385D4 | Tb927.11.6350 | BDF7 | 7.67914 | 14.416 |
| Q582P7 | Tb927.3.4880 | nucleosome assembly protein (NAP2) | 5.41527 | 10.1791 |
| Q4GZ00 | Tb927.1.2210 | nucleosome assembly protein (NAP1) | 5.53343 | 8.66135 |
| Q38D21 | Tb927.9.14430 | Casein kinase II | 4.07334 | 7.92549 |
| Q383J7 | Tb927.11.10820 | Casein kinase II subunit beta | 5.39243 | 7.86722 |
| Q386Y7 | Tb927.11.1670 | cysteine desulfurase | 5.11041 | 6.62681 |
| Q585P0 | Tb927.6.1710 | hypothetical protein, conserved | 2.99608 | 6.03265 |
| Q586Q9 | Tb927.2.4580 | UNC119 | 2.88511 | 4.59575 |
| Q385C7 | Tb927.11.6420 | hypothetical protein, conserved | 5.21268 | 4.53223 |
| Q388S2 | Tb927.10.13250 | hypothetical protein, conserved | 3.00103 | 4.37745 |
| Q388P7 | Tb927.10.13510 | zinc metallopeptidase | 4.53851 | 4.27171 |
| Q381W2 | Tb927.11.14730 | Metalloprotease M41 FtsH | 2.77481 | 4.13688 |
| Q389E2 | Tb927.10.12000 | iron-sulfur cluster assembly protein | 4.06848 | 3.71724 |
| Q38DH1 | Tb927.9.12340 | hypothetical protein, conserved | 5.25919 | 3.69929 |
| Q387Z3 | Tb927.10.15180 | nucleosome assembly protein (NAP3) | 4.13692 | 3.43985 |
| Q383E9 | Tb927.11.11290 | heat shock protein 70 | 2.87951 | 3.26538 |
| Q38BN5 | Tb927.10.3640 | nuclear transmembrane protein | 3.5369 | 3.14475 |
| Q57WG0 | Tb927.7.3450 | protein I/6 | 2.31469 | 3.07365 |
| Q38BJ3 | Tb927.10.4070 | hypothetical protein, conserved | 2.77233 | 3.07108 |
| Q386R6 | Tb927.11.2440 | Tubulin cofactor C domain-containing protein 1 | 5.83909 | 3.04755 |
| Q57YB8 | Tb927.7.3080 | Kinetochore interacting protein 4 (KKIP4) | 4.07542 | 3.00762 |
| Q584Z0 | Tb927.6.3500 | endosomal trafficking protein RME-8 | 2.8357 | 2.94232 |
| B3GVR6 | Tb427.BES126.2 | Expression site-associated gene 7 (ESAG7) protein | 3.175 | 2.75723 |
| Q584K3 | Tb927.4.5020 | DNA-directed RNA polymerase II subunit RPB1-A | 3.43896 | 2.68048 |
| Q585L0 | Tb927.4.3400 | hypothetical protein, conserved | 3.55021 | 2.30728 |
| B3GVQ1 | Tb427.BES56.7 | Expression site-associated gene 8 (ESAG8) protein | 3.32151 | 2.22846 |
| Q38D92 | Tb927.9.13320 | hypothetical protein, conserved | 2.8659 | 2.16731 |
| Q384U3 | Tb927.11.7220 | neurobeachin/beige protein | 2.1597 | 2.06637 |
| Q580Q4 | Tb927.4.5310 | Repressor of differentiation kinase 2 | 3.01007 | 2.05326 |

### TFIIS2-2 IP significantly enriched interactors (over untagged control)

| UniProt ID | TriTryp ID | Description | $-log_{10}$ Student's t-test p-value | $log_2$ (tagged/untagged) |
|---|---|---|---|---|
| Q582N0 | Tb927.3.5070 | hypothetical protein | 4.99559 | 14.1957 |
| Q38DC5 | Tb927.9.12900 | RNA polymerase-associated protein LEO1 | 8.53225 | 14.0585 |
| Q57V64 | Tb927.3.3220 | RNA polymerase-associated protein CTR9 | 5.09399 | 13.7842 |
| Q586X9 | Tb927.2.3580 | transcription elongation factor s-II (TFIIS2-1) | 4.71915 | 13.7123 |
| Q57UN3 | Tb927.7.4030 | hypothetical protein | 4.37899 | 12.2545 |
| Q383Q6 | Tb927.11.10230 | RNA polymerase-associated protein CDC73 | 4.83204 | 11.654 |
| Q586Y0 | Tb927.2.3480 | transcription elongation factor s-II (TFIIS2-2) | 4.58521 | 11.4279 |
| Q586C8 | Tb927.2.5810 | Holliday-junction resolvase-like of SPT6/SH2 domain | 3.46556 | 6.32434 |
| Q583H5 | Tb927.4.3810 | DNA-directed RNA polymerase II subunit 2 (RPB2) | 4.72879 | 5.87253 |
| Q57YU2 | Tb927.8.7400 | RNA polymerase IIA largest subunit (RPB1) | 2.56766 | 5.65476 |
| Q383E9 | Tb927.11.11290 | heat shock protein 70 | 3.89194 | 5.60172 |
| Q57Z69 | Tb927.5.3240 | hypothetical protein | 3.53278 | 5.28492 |
| Q38C15 | Tb927.10.2290 | chaperone protein DnaJ | 3.90992 | 5.05717 |
| Q38BC1 | Tb927.10.4800 | hypothetical protein | 4.30469 | 4.65933 |
| Q580N2 | Tb927.3.4100 | Pyruvate transporter | 3.01628 | 4.3271 |
| Q38DK9 | Tb927.9.11850 | structural maintenance of chromosome 1 (SMC1) | 4.50362 | 4.13835 |
| Q4GYZ8 | Tb927.1.2230 | small myristoylated protein 1-1 | 2.92265 | 4.08843 |
| Q38DH5 | Tb927.9.12300 | replication factor C, subunit 3 | 3.86106 | 3.9829 |
| Q38AE0 | Tb927.10.8240 | hypothetical protein | 4.76951 | 3.95355 |
| Q57X47 | Tb927.5.4570 | Flagellum adhesion protein 3 | 3.91342 | 3.95229 |
| Q389A2 | Tb927.10.12410 | hypothetical protein | 3.78775 | 3.88992 |
| Q57V04 | Tb927.7.2070 | heat shock protein DNAJ | 3.60006 | 3.81925 |
| Q9N942 | Tb927.1.420 | retrotransposon hot spot protein 5 (RHS5) | 2.72687 | 3.73281 |
| Q580S5 | Tb927.3.5500 | DNA-directed RNA polymerase II subunit 3 | 4.01615 | 3.69604 |
| Q383S2 | Tb927.11.10070 | BDF3 | 2.12292 | 3.67039 |
| Q389E6 | Tb927.10.11960 | hypothetical protein | 4.15932 | 3.66012 |
| Q582N5 | Tb927.3.5020 | Flagellar Member 6 | 2.3036 | 3.64086 |
| Q381Q0 | Tb927.11.15350 | RNA-binding protein | 4.57513 | 3.61854 |
| Q383X4 | Tb927.11.9550 | replication factor C, subunit 4 | 3.8174 | 3.54192 |
| Q57YB8 | Tb927.7.3080 | Kinetochore interacting protein 4 | 4.60916 | 3.45554 |

### TFIIS2-2 IP significantly enriched interactors (over untagged control) - continued

| UniProt ID | TriTryp ID | Description | $-log_{10}$ Student's t-test p-value | $log_2$ (tagged/untagged) |
|---|---|---|---|---|
| Q382X6 | Tb927.11.12050 | hypothetical protein | 3.11596 | 3.4261 |
| Q385C3 | Tb927.11.6460 | hypothetical protein | 3.11483 | 3.40504 |
| Q388C7 | Tb927.10.14770 | Associated kinase of Tb14-3-3 | 3.33222 | 3.40478 |
| Q385K3 | Tb927.11.5650 | replication factor C, subunit 1 | 3.48977 | 3.38828 |
| Q38D92 | Tb927.9.13320 | hypothetical protein | 4.34031 | 3.1995 |
| Q381Y8 | Tb927.11.14490 | RNA polymerase subunit (RPB7) | 2.88983 | 3.1697 |
| Q585Y7 | Tb927.6.3890 | replication factor C, subunit 2 | 3.38762 | 3.12531 |
| Q386Y8 | Tb927.11.1660 | vesicular transport protein (CDC48 homologue) | 3.59205 | 3.10783 |
| Q38AG5 | Tb927.10.7990 | replication factor C subunit 3 | 2.73335 | 3.06639 |
| Q581W7 | Tb927.8.2150 | hypothetical protein | 2.0861 | 2.96903 |
| Q387X4 | Tb927.10.15390 | Flagellum attachment zone protein 7 | 2.13513 | 2.93651 |
| Q38D56 | Tb927.9.13970 | hypothetical protein | 2.66294 | 2.87394 |
| Q381J5 | Tb927.11.15910 | iron superoxide dismutase | 2.7698 | 2.85969 |
| Q582W2 | Tb927.3.2490 | hypothetical protein | 2.22011 | 2.83506 |
| Q381B2 | Tb927.11.16740 | chaperone protein DnaJ | 3.3618 | 2.75458 |
| Q57WL5 | Tb927.8.8150 | C2 domain containing protein | 3.36274 | 2.73377 |
| Q580R3 | Tb927.3.5620 | FACT complex subunit SPT16 | 2.30752 | 2.71646 |
| Q38D21 | Tb927.9.14430 | Casein kinase II | 3.7948 | 2.66072 |
| Q388V2 | Tb927.10.12940 | predicted zinc finger protein | 2.18054 | 2.66 |
| Q386Q6 | Tb927.11.2540 | hypothetical protein | 2.50232 | 2.54926 |
| Q388R5 | Tb927.10.13320 | DNA-directed RNA polymerase II/III subunit | 2.82681 | 2.48833 |
| Q38DM0 | Tb927.9.11740 | cyclophilin type peptidyl-prolyl cis-trans isomerase | 4.05265 | 2.44251 |
| Q388X5 | Tb927.10.12680 | 60S ribosomal protein L34 | 3.33235 | 2.40742 |
| Q38EX8 | Tb927.9.5770 | Tryparedoxin peroxidase | 3.94584 | 2.40469 |
| Q583Q4 | Tb927.6.2800 | hypothetical protein | 4.36211 | 2.31914 |
| Q386D5 | Tb927.11.3770 | Dpy-30 motif containing protein | 2.61587 | 2.28773 |
| Q387U0 | Tb927.10.15740 | hypothetical protein | 3.34011 | 2.27737 |
| Q38DT2 | Tb927.9.10890 | Regulatory subunit of type II PKA R-subunit | 2.22984 | 2.2477 |
| Q583S0 | Tb927.6.2640 | importin alpha subunit | 2.42317 | 2.22998 |
| Q38E79 | Tb927.9.9100 | hypothetical protein | 3.45709 | 2.22126 |

### TFIIS2-2 IP significantly enriched interactors (over untagged control) - continued

| UniProt ID | TriTryp ID | Description | $-log_{10}$ Student's t-test p-value | $log_2$ (tagged/untagged) |
|---|---|---|---|---|
| Q381F0 | Tb927.11.16360 | hypothetical protein | 2.14231 | 2.15977 |

### DOT1A IP significantly enriched interactors (over untagged control)

| UniProt ID | TriTryp ID | Description | $-\log_{10}$ Student's t-test p-value | $\log_2$ (tagged/untagged) |
|---|---|---|---|---|
| Q4GYF6 | Tb927.1.4220 | Ydr279p protein family (RNase H2 complex component) | 5.13483 | 10.0395 |
| Q581Z0 | Tb927.8.1920 | DOT1A | 4.76344 | 9.91715 |
| Q38B94 | Tb927.10.5070 | ribonuclease H | 6.02426 | 9.22535 |
| Q38F34 | Tb927.9.5190 | proliferating cell nuclear antigen (PCNA) | 5.74715 | 9.04071 |
| Q4GYA5 | Tb927.1.4730 | Ribonuclease H2 non-catalytic subunit (Ylr154p-like) | 5.55902 | 7.96804 |
| Q583S0 | Tb927.6.2640 | importin alpha subunit | 5.11794 | 4.35021 |
| Q586Q9 | Tb927.2.4580 | UNC119 | 3.18195 | 3.15221 |
| Q387Q3 | Tb927.10.16120 | IMPDH1 | 3.77913 | 2.54422 |
| Q383E9 | Tb927.11.11290 | HSP70 | 3.62208 | 2.51085 |

### TRF IP significantly enriched interactors (over untagged control)

| UniProt ID | TriTryp ID | Description | $-log_{10}$ Student's t-test p-value | $log_2$ (tagged/untagged) |
|---|---|---|---|---|
| Q388W1 | Tb927.10.12850 | ttaggg binding factor (TRF) | 4.5821 | 10.2585 |
| Q586Z9 | Tb927.6.4330 | telomere-associated protein | 5.17342 | 9.60136 |
| Q38FD7 | Tb927.9.4000 | hypothetical protein, conserved | 7.10736 | 9.53791 |
| Q57XY9 | Tb927.3.1560 | TRF-Interacting Factor 2 (TIF2) | 5.76737 | 9.27732 |
| Q383U2 | Tb927.11.9870 | Telomere-associated protein 1 (TelAP1) | 5.50112 | 8.97668 |
| Q385L3 | Tb927.11.5550 | DNA polymerase theta (POLQ) | 5.76751 | 8.19689 |
| Q38BZ3 | Tb927.10.2520 | PrimPol-like protein 2 | 5.07689 | 7.84982 |
| Q387L4 | Tb927.11.370 | repressor activator protein 1 (RAP1) | 3.75433 | 6.0696 |
| Q586J9 | Tb927.2.2190 | histone deacetylase 3 (HDAC3) | 3.22852 | 4.91487 |
| Q387X7 | Tb927.10.15350 | Histone H3.V | 2.85869 | 3.715 |
| Q385E2 | Tb927.11.6270 | inositol-1,4,5-trisphosphate (IP3) 5-phosphatase | 4.2341 | 3.60051 |
| Q38FR6 | Tb927.9.2520 | microtubule-associated protein (GB4) | 2.41828 | 3.5649 |
| Q57YA3 | Tb927.7.2820 | Histone H2A | 2.40159 | 3.11452 |
| Q586Q9 | Tb927.2.4580 | UNC119 | 2.78003 | 2.81699 |
| Q583S0 | Tb927.6.2640 | importin alpha subunit | 2.59694 | 2.64684 |
| Q57VM1 | Tb927.8.1330 | 60S ribosomal protein L7a | 3.4478 | 2.59139 |
| Q388B1 | Tb927.10.14950 | Zinc finger CCCH domain-containing protein 40 (ZC3H40) | 3.0887 | 2.5017 |
| Q384T0 | Tb927.11.7350 | Histone H2B.V | 2.51354 | 2.43428 |
| Q38CW9 | Tb927.9.15420 | 60S ribosomal protein L32 | 2.98648 | 2.14164 |
| Q38AC1 | Tb927.10.8430 | 40S ribosomal protein S12 | 2.40777 | 2.05111 |
| Q38EY6 | Tb927.9.5690 | 60S acidic ribosomal protein | 2.4053 | 2.03389 |

### TBP IP significantly enriched interactors (over untagged control)

| UniProt ID | TriTryp ID | Description | $-\log_{10}$ Student's t-test p-value | $\log_2$ (tagged/untagged) |
|---|---|---|---|---|
| Q38E05 | Tb927.9.9970 | Small nuclear RNA gene activation protein 50 (SNAP50) | 5.10704 | 10.3893 |
| Q599N4 | Tb927.10.15570 | Transcription factor IIA alpha-beta subunit (TFIIA-1) | 5.57909 | 10.3761 |
| Q599N5 | Tb927.10.7070 | Small nuclear RNA gene activation protein 3 (SNAP3) | 5.25598 | 10.1384 |
| Q387R9 | Tb927.10.15950 | TATA-box-binding protein (TBP) | 5.05796 | 9.56953 |
| Q599N6 | Tb927.5.3910 | Small nuclear RNA activating protein 2 (SNAP2) | 5.26631 | 9.28801 |
| Q387K4 | Tb927.11.470 | TFIIB-related factor BRF1 | 6.2795 | 9.00377 |
| Q599N3 | Tb927.10.4840 | Transcription factor IIA gamma subunit | 5.52352 | 8.6298 |
| Q387V6 | Tb927.10.15570 | transcription factor IIA (TFIIA-1) | 5.33766 | 7.33029 |
| Q38DC3 | Tb927.9.12950 | hypothetical protein, conserved | 4.03431 | 7.05365 |
| Q384L9 | Tb927.11.8000 | hypothetical protein, conserved | 3.84548 | 6.45067 |

### ELP3b IP significantly enriched interactors (over untagged control)

| UniProt ID | TriTryp ID | Description | $-log_{10}$ Student's t-test p-value | $log_2$ (tagged/untagged) |
|---|---|---|---|---|
| Q57YY2 | Tb927.8.3310 | Elongator-like Protein 3b (ELP3b) | 4.31259 | 12.0095 |
| Q585H2 | Tb927.2.1210 | retrotransposon hot spot protein 4 (RHS4) | 2.23588 | 2.67507 |
| Q586Q9 | Tb927.2.4580 | UNC119 | 2.91717 | 2.56359 |
| Q57ZH2 | Tb927.7.5170 | 60S ribosomal protein L25 | 2.66388 | 2.39845 |
| Q583F5 | Tb927.4.4700 | hypothetical protein, conserved | 2.01684 | 2.24062 |
| Q387A9 | Tb927.11.1430 | Component of motile flagella 2 | 2.18741 | 2.04843 |

### PHD2 IP significantly enriched interactors (over untagged control)

| UniProt ID | TriTryp ID | Description | $-\log_{10}$ Student's t-test p-value | $\log_2$ (tagged/untagged) |
|---|---|---|---|---|
| Q387D3 | Tb11.v5.0388 | Ring finger domain/PHD-finger (PHD2) | 3.9765 | 7.4299 |

### PHD4 IP significantly enriched interactors (over untagged control)

| UniProt ID | TriTryp ID | Description | $-log_{10}$ Student's t-test p-value | $log_2$ (tagged/untagged) |
|---|---|---|---|---|
| Q57Z97 | Tb927.3.2140 | transcription activator (PHD4) | 5.39723 | 9.03261 |
| Q587E7 | Tb927.2.3370 | UDP-Gal or UDP-GlcNAc-dependent glycosyltransferase | 2.30539 | 2.0727 |

### PHD1 IP significantly enriched interactors (over untagged control)

| UniProt ID | TriTryp ID | Description | $-\log_{10}$ Student's t-test p-value | $\log_2$ (tagged/untagged) |
|---|---|---|---|---|
| Q389Y3 | Tb927.10.9930 | PHD1 | 5.75346 | 10.6464 |
| Q384N0 | Tb927.11.7880 | hypothetical protein, conserved | 4.14919 | 10.3396 |
| Q38FN2 | Tb927.9.2910 | EAF6 (NuA4 complex component) | 4.78537 | 8.70341 |
| Q38AD3 | Tb927.10.8310 | HAT3 | 5.36842 | 7.71911 |

### HAT3 IP significantly enriched interactors (over untagged control)

| UniProt ID | TriTryp ID | Description | $-log_{10}$ Student's t-test p-value | $log_2$ (tagged/untagged) |
|---|---|---|---|---|
| Q389Y3 | Tb927.10.9930 | PHD1 | 4.63916 | 7.69124 |
| Q384N0 | Tb927.11.7880 | hypothetical protein, conserved | 4.3526 | 6.73332 |
| Q38FN2 | Tb927.9.2910 | EAF6 (NuA4 complex component) | 3.41918 | 5.63806 |
| Q38AD3 | Tb927.10.8310 | HAT3 | 3.25866 | 5.62241 |
| Q57YG1 | Tb927.8.690 | peptidyl-prolyl cis-trans isomerase/rotamase, putative | 2.94219 | 2.48432 |
| B3GVR6 | Tb427.BES126.2 | Expression site-associated gene 7 (ESAG7) protein | 2.29797 | 2.02109 |

### AGO1 IP significantly enriched interactors (over untagged control)

| UniProt ID | TriTryp ID | Description | $-\log_{10}$ Student's t-test p-value | $\log_2$ (tagged/untagged) |
|---|---|---|---|---|
| Q389P5 | Tb927.10.10850 | AGO1 | 5.65625 | 13.9389 |
| Q38A86 | Tb927.10.8880 | RNA interference factor 4 (RIF4) | 4.02946 | 9.3504 |
| Q586Q9 | Tb927.2.4580 | UNC119 | 2.68214 | 5.73447 |
| Q389B0 | Tb927.10.12330 | zinc finger protein family member, putative (ZC3H34) | 3.65955 | 3.58715 |
| Q38E63 | Tb927.9.9290 | polyadenylate-binding protein 1 (PABP1) | 2.42994 | 2.75077 |
| Q585M4 | Tb927.6.1870 | Eukaryotic translation initiation factor 4E-4 (EIF4E4) | 3.06851 | 2.25288 |

### NUP110 IP significantly enriched interactors (over untagged control)

| UniProt ID | TriTryp ID | Description | $-log_{10}$ Student's t-test p-value | $log_2$ (tagged/untagged) |
|---|---|---|---|---|
| Q387L8 | Tb927.11.330 | Myosin-like protein 1 (NUP110/MLP1) | 6.08194 | 11.7416 |
| Q38ET5 | Tb927.9.6460 | hypothetical protein, conserved | 6.79431 | 11.5223 |
| Q57V68 | Tb927.3.3180 | Nucleoporin NUP98 | 4.35025 | 10.262 |
| Q38G01 | Tb927.9.1410 | SUMO-interacting motif-containing protein | 4.81713 | 9.78628 |
| Q57Y35 | Tb927.4.4310 | Nucleoporin NUP64 | 7.06158 | 9.52875 |
| Q586V1 | Tb927.2.4230 | NUP-1 protein | 4.74201 | 9.45264 |
| Q38A10 | Tb927.10.9650 | Nucleoporin NUP152 | 5.0676 | 9.18434 |
| Q583W5 | Tb927.4.2880 | Nucleoporin NUP225 | 4.87759 | 9.14371 |
| Q38FF7 | Tb927.9.3760 | Nucleoporin GLE2 | 3.78726 | 9.03303 |
| Q57Y62 | Tb927.7.6670 | hypothetical protein, conserved | 5.92805 | 8.99941 |
| Q38AQ7 | Tb927.10.7060 | Nucleoporin NUP96 | 5.03288 | 8.56275 |
| Q581B5 | Tb927.8.3950 | hypothetical protein, conserved | 5.4272 | 8.53964 |
| Q386L6 | Tb927.11.2950 | Nucleoporin NUP89 | 3.95216 | 8.50077 |
| Q387F2 | Tb927.11.980 | Nucleoporin NUP158 | 4.5024 | 8.43801 |
| Q381I7 | Tb927.11.15990 | Nucleoporin NUP109 | 5.77776 | 8.38197 |
| Q38BL8 | Tb927.10.3810 | Nucleoporin NUP65 | 3.99843 | 8.27312 |
| Q38D32 | Tb927.9.14240 | Nucleoporin NUP82 | 5.01823 | 8.05696 |
| Q57XP4 | Tb927.7.2300 | Nucleoporin NUP132 | 3.24449 | 7.7052 |
| Q585F7 | Tb927.6.890 | hypothetical protein, conserved | 4.70788 | 7.70289 |
| Q584I6 | Tb927.4.5200 | Nucleoporin NUP62 | 5.5861 | 7.33425 |
| Q582Q9 | Tb927.3.4760 | dynamin-1-like protein | 3.45599 | 7.25029 |
| Q383V1 | Tb927.11.9780 | Nucleoporin NUP119 | 5.49855 | 7.20329 |
| Q38BN5 | Tb927.10.3640 | nuclear transmembrane protein | 5.32585 | 7.17 |
| Q38AE7 | Tb927.10.8170 | Nucleoporin NUP144 | 5.5177 | 7.15809 |
| Q38DR0 | Tb927.9.11150 | hypothetical protein, conserved | 5.07784 | 7.13933 |
| Q385W3 | Tb927.11.4540 | Nucleoporin NUP48 | 4.22511 | 7.11522 |
| Q57XK4 | Tb927.3.3540 | Nucleoporin NUP53b | 5.40541 | 6.91711 |
| Q57X92 | Tb927.7.4760 | hypothetical protein, conserved | 3.40266 | 6.81533 |
| Q388I4 | Tb927.10.14180 | protein transport protein SEC13 | 4.52415 | 6.74846 |
| Q38C12 | Tb927.10.2320 | Nucleoporin NUP41 | 3.72137 | 6.73405 |

### NUP110 IP significantly enriched interactors (over untagged control) - continued

| UniProt ID | TriTryp ID | Description | $-log_{10}$ Student's t-test p-value | $log_2$ (tagged/untagged) |
|---|---|---|---|---|
| Q57XE3 | Tb927.8.2460 | hypothetical protein, conserved | 3.35065 | 6.70225 |
| Q387F3 | Tb927.11.970 | hypothetical protein, conserved | 2.93832 | 6.39177 |
| Q38FU0 | Tb927.9.2220 | SUMO1/Ulp2 | 5.74657 | 6.38493 |
| Q383Z7 | Tb927.11.9320 | hypothetical protein, conserved | 4.94793 | 6.33722 |
| Q385L2 | Tb927.11.5560 | hypothetical protein, conserved | 4.02665 | 6.2611 |
| Q57U40 | Tb927.8.8050 | Nucleoporin NUP75 | 3.16413 | 6.24056 |
| Q381M9 | Tb927.11.15560 | Nucleoporin NUP53a | 3.87782 | 6.23161 |
| Q386D3 | Tb927.11.3790 | hypothetical protein, conserved | 3.91642 | 6.03219 |
| Q382M8 | Tb927.11.13080 | hypothetical protein, conserved | 3.32361 | 5.66999 |
| Q582J8 | Tb927.7.5830 | hypothetical protein, conserved | 3.66063 | 5.1738 |
| Q38AE8 | Tb927.10.8160 | hypothetical protein, conserved | 5.94554 | 5.10643 |
| Q38A83 | Tb927.10.8910 | Nucleoporin NUP181 | 3.65194 | 5.07702 |
| Q582R3 | Tb927.3.4720 | dynamin-1-like protein | 3.51272 | 5.03322 |
| Q57Z74 | Tb927.5.3190 | hypothetical protein, conserved | 4.60178 | 4.91148 |
| Q585L9 | Tb927.6.1920 | hypothetical protein, conserved | 3.56403 | 4.90318 |
| Q57V60 | Tb927.3.3260 | hypothetical protein, conserved | 4.30145 | 4.77298 |
| Q57UY5 | Tb927.8.4660 | hypothetical protein, conserved | 3.43363 | 4.69182 |
| Q4FKD8 | Tb927.11.18680 | dynein light chain LC8 | 2.70859 | 4.49924 |
| Q57WZ3 | Tb927.3.1120 | GTP-binding nuclear protein rtb2 | 4.82583 | 4.31706 |
| Q57YP5 | Tb927.8.6440 | RNA-binding protein (RBP20) | 5.12896 | 3.95232 |
| Q38FR1 | Tb927.9.2590 | hypothetical protein, conserved | 4.57717 | 3.93099 |
| Q386H3 | Tb927.11.3380 | Ran-binding protein 1 | 2.6877 | 3.86423 |
| Q57ZJ7 | Tb927.3.3150 | hypothetical protein, conserved | 3.92573 | 3.62835 |
| Q383H0 | Tb927.11.11080 | Nucleoporin NUP149 | 4.76231 | 3.50611 |
| Q383G9 | Tb927.11.11090 | Nucleoporin NUP140 | 4.08639 | 3.39356 |
| Q386S5 | Tb927.11.2340 | hypothetical protein, conserved | 3.82124 | 3.34146 |
| Q583W2 | Tb927.4.2850 | hypothetical protein, conserved | 2.9211 | 3.25883 |
| Q57YE9 | Tb927.8.820 | VID27 cytoplasmic protein | 2.89878 | 3.20937 |
| Q38AJ6 | Tb927.10.7680 | GTPase activating protein | 3.06235 | 3.20847 |
| Q57YM6 | Tb927.8.6250 | Nucleoporin NUP76 | 2.14152 | 3.13853 |

### NUP110 IP significantly enriched interactors (over untagged control) - continued

| UniProt ID | TriTryp ID | Description | $-log_{10}$ Student's t-test p-value | $log_2$ (tagged/untagged) |
|---|---|---|---|---|
| Q384L3 | Tb927.11.8060 | SNARE associated Golgi protein | 2.41413 | 3.12 |
| Q582S7 | Tb927.3.4580 | hypothetical protein, conserved | 2.85737 | 3.05497 |
| Q386Q0 | Tb927.11.2600 | hypothetical protein, conserved | 4.03664 | 2.76767 |
| P35315 | Tb927.8.1180 | vacuolar-type Ca2+-ATPase 1 | 3.13313 | 2.73411 |
| Q583S0 | Tb927.6.2640 | importin alpha subunit | 3.01621 | 2.72712 |
| Q582J1 | Tb927.7.5760 | mRNA transport regulator MTR2 | 3.00139 | 2.35135 |
| Q38G07 | Tb927.9.1340 | Myosin-like protein 2 (MLP2) | 4.006 | 2.12899 |
| Q586Q9 | Tb927.2.4580 | UNC119 | 2.52185 | 2.11811 |

### SET13 IP significantly enriched interactors (over untagged control)

| UniProt ID | TriTryp ID | Description | $-log_{10}$ Student's t-test p-value | $log_2$ (tagged/untagged) |
|---|---|---|---|---|
| Q584E3 | Tb927.4.2440 | SET13 | 4.24447 | 7.11086 |
| Q57VU5 | Tb927.7.990 | chaperone protein DnaJ, putative | 2.06949 | 4.64322 |

### SET15 IP significantly enriched interactors (over untagged control)

| UniProt ID | TriTryp ID | Description | $-\log_{10}$ Student's t-test p-value | $\log_2$ (tagged/untagged) |
|---|---|---|---|---|
| Q57ZV3 | Tb927.5.2770 | SET15 | 5.19135 | 13.137 |
| Q57UH7 | Tb927.8.5050 | OTU-like cysteine protease | 4.49346 | 12.3099 |
| Q388D4 | Tb927.10.14700 | hypothetical protein, conserved | 3.02213 | 7.88572 |
| Q585P0 | Tb927.6.1710 | hypothetical protein, conserved | 4.92275 | 7.63556 |
| Q57ZS7 | Tb927.5.2080 | GMP reductase | 2.90688 | 7.40885 |
| Q38DN9 | Tb927.9.11470 | 60S ribosomal protein L27a | 3.00961 | 6.90772 |
| Q386Y7 | Tb927.11.1670 | cysteine desulfurase | 3.27227 | 6.90033 |
| Q386I1 | Tb927.11.3300 | Spindle assembly abnormal 4 | 3.63865 | 6.86669 |
| Q38C20 | Tb927.10.2240 | RNA binding protein | 3.00736 | 6.50689 |
| Q38EZ6 | Tb927.9.5590 | DNA topoisomerase ii (TOP2) | 3.25468 | 6.42083 |
| Q389X5 | Tb927.10.10010 | mRNA turnover protein 4 homolog | 3.57797 | 6.20563 |
| Q38CY6 | Tb927.9.15210 | ribosomal protein L36 | 3.98462 | 6.06961 |
| Q386T4 | Tb927.11.2250 | conserved protein, unknown function | 4.93118 | 5.83651 |
| Q38A76 | Tb927.10.8980 | hypothetical protein, conserved | 6.21198 | 5.68091 |
| Q381T9 | Tb927.11.14960 | pumilio/PUF RNA binding protein 7 | 3.73724 | 5.66011 |
| Q57UX0 | Tb927.8.4820 | eukaryotic translation initiation factor 4 gamma 3 | 3.43404 | 5.66001 |
| Q386C9 | Tb927.11.3850 | AMP deaminase | 4.46495 | 5.64093 |
| D7SG81 | Tb927.11.3230 | 60S ribosomal protein L44 | 4.29375 | 5.6298 |
| Q385R2 | Tb927.11.5060 | Mitochondrial small ribosomal subunit Rsm22 | 3.92785 | 5.61396 |
| Q586G9 | Tb927.6.4970 | serine/arginine-rich protein specific kinase SRPK | 3.42122 | 5.54974 |
| Q582F1 | Tb927.5.2140 | regulator of nonsense transcripts 1 | 6.18513 | 5.51517 |
| Q388D6 | Tb927.10.14680 | ribosome biogenesis protein | 3.13061 | 5.50463 |
| Q38FW8 | Tb927.9.1810 | 60S ribosomal protein L35 | 3.4727 | 5.47661 |
| Q383S6 | Tb927.11.10030 | 60S ribosomal protein L29 | 4.35516 | 5.47136 |
| Q38D54 | Tb927.9.13990 | RNA-binding protein | 4.24849 | 5.4077 |
| Q57XZ3 | Tb927.4.3890 | ATP-dependent RNA helicase | 3.29763 | 5.34227 |
| Q57YV4 | Tb927.8.7260 | kinetoplast-associated protein | 3.70907 | 5.30335 |
| Q583S9 | Tb927.6.740 | ATP-dependent DEAH-box RNA helicase | 3.51075 | 5.28101 |
| Q57VA5 | Tb927.5.1580 | zinc finger CCCH domain containing protein 13 | 3.54964 | 5.26218 |
| Q384V7 | Tb927.11.7140 | cell cycle sequence binding phosphoprotein (RBP33) | 2.95492 | 5.24986 |

### SET15 IP significantly enriched interactors (over untagged control) - continued

| UniProt ID | TriTryp ID | Description | $-log_{10}$ Student's t-test p-value | $log_2$ (tagged/untagged) |
|---|---|---|---|---|
| Q382K0 | Tb927.11.13360 | AAA ATPase | 5.10819 | 5.19323 |
| Q57WS6 | Tb927.7.1240 | Sphingosine kinase | 4.67664 | 5.18177 |
| Q389Y5 | Tb927.10.9910 | PSP1 C-terminal conserved region | 2.50741 | 5.09675 |
| Q584Q1 | Tb927.6.2550 | RNA-binding protein | 2.0502 | 5.07847 |
| Q57U27 | Tb927.7.270 | Ribosome production factor 2 homolog | 3.50337 | 5.07752 |
| Q389B0 | Tb927.10.12330 | zinc finger protein family member | 3.04493 | 4.97903 |
| Q388N7 | Tb927.10.13610 | zinc-finger of a C2HC-type/Double zinc ribbon | 3.68603 | 4.92499 |
| Q38EA4 | Tb927.9.8720 | Fructose-1,6-bisphosphate, cytosolic | 2.81419 | 4.9025 |
| Q580H2 | Tb927.8.1510 | ATP-dependent DEAD/H RNA helicase | 3.84264 | 4.89523 |
| Q38B98 | Tb927.10.5030 | ubiquitin/ribosomal protein S27a | 2.61783 | 4.84472 |
| Q585M4 | Tb927.6.1870 | Eukaryotic translation initiation factor 4E | 3.27292 | 4.7831 |
| Q38E96 | Tb927.9.8820 | hypothetical protein | 3.76301 | 4.74228 |
| Q38E63 | Tb927.9.9290 | Polyadenylate-binding protein (PABP) | 3.30433 | 4.70896 |
| Q38AL4 | Tb927.10.7500 | Fibrillarin | 2.30138 | 4.7049 |
| Q57W71 | Tb927.8.3020 | Q_MOTIF domain-containing protein | 2.70837 | 4.67721 |
| Q38CZ4 | Tb927.9.15060 | rRNA processing protein | 4.09063 | 4.67001 |
| Q384T4 | Tb927.11.7310 | RNA binding protein | 3.08847 | 4.622 |
| Q583E3 | Tb927.4.4580 | hypothetical protein | 2.77331 | 4.5719 |
| Q386I3 | Tb927.11.3280 | Kinesin-like protein | 4.65668 | 4.54025 |
| Q384Z1 | Tb927.11.6790 | Ribosome biogenesis protein BOP1 homolog | 4.49999 | 4.46505 |
| D6XKD1 | Tb927.7.2140 | hypothetical protein | 3.36301 | 4.45784 |
| Q57W56 | Tb927.11.16280 | 60S ribosomal protein L2 | 3.20381 | 4.43985 |
| Q586X7 | Tb927.2.3620 | GTP-binding elongation factor Tu family | 3.28946 | 4.4312 |
| Q57Y68 | Tb927.7.6610 | FHA domain-containing protein | 3.46503 | 4.41503 |
| Q384J9 | Tb927.11.8200 | 40S ribosomal protein S26 | 2.24998 | 4.41346 |
| Q38B42 | Tb927.10.5620 | Fructose-bisphosphate aldolase | 3.94919 | 4.35694 |
| Q38AY0 | Tb927.10.6260 | ATP-dependent DEAD/H RNA helicase | 3.07219 | 4.31092 |
| Q38C15 | Tb927.10.2290 | chaperone protein DnaJ | 3.22005 | 4.26772 |
| Q57V37 | Tb927.7.1730 | 60S ribosomal protein L7 | 5.53509 | 4.26439 |
| Q386D1 | Tb927.11.3830 | hypothetical protein | 3.69379 | 4.24606 |

### SET15 IP significantly enriched interactors (over untagged control) - continued

| UniProt ID | TriTryp ID | Description | $-log_{10}$ Student's t-test p-value | $log_2$ (tagged/untagged) |
|---|---|---|---|---|
| Q586X0 | Tb927.2.3880 | Heterogeneous nuclear ribonucleoprotein H/F | 2.1852 | 4.24504 |
| Q38DU2 | Tb927.9.10770 | Polyadenylate-binding protein (PABP) | 4.16027 | 4.23051 |
| Q57YL9 | Tb927.8.6180 | 60S ribosomal protein L26 | 2.44842 | 4.20526 |
| Q57XA6 | Tb927.7.4900 | 5'-3' exonuclease XRNA | 2.33263 | 4.20182 |
| Q386J9 | Tb927.11.3120 | Nucleolar GTP-binding protein 1 | 3.70431 | 4.17426 |
| Q580D8 | Tb927.4.1100 | Ribosomal protein L21E (60S) | 2.72672 | 4.15507 |
| Q38C87 | Tb927.10.1540 | hypothetical protein | 3.57026 | 4.14729 |
| Q584N0 | Tb927.2.380 | Retrotransposon hot spot (RHS) protein | 3.6059 | 4.12329 |
| Q38AI3 | Tb927.10.7810 | hypothetical protein | 3.18018 | 4.08699 |
| Q382C4 | Tb927.10.3840 | 60S ribosomal protein L18a | 3.48248 | 4.07294 |
| Q38E42 | Tb927.9.9520 | hypothetical protein | 3.17523 | 4.06933 |
| Q386J2 | Tb927.11.3190.1 | LisH domain-containing protein | 2.63572 | 4.05821 |
| Q57VC9 | Tb927.5.1340 | hypothetical protein | 3.54771 | 4.05029 |
| Q389E2 | Tb927.10.12000 | iron-sulfur cluster assembly protein | 3.03889 | 4.03155 |
| Q57VP3 | Tb927.10.5610 | 40S ribosomal protein S9 | 3.82509 | 4.01799 |
| Q38CW9 | Tb927.9.15440 | 60S ribosomal protein L32 | 3.52884 | 3.99987 |
| Q57ZF2 | Tb927.3.1590 | hypothetical protein | 3.69145 | 3.98869 |
| Q38D76 | Tb927.9.13510 | Poly(A)-specific ribonuclease PARN-3 | 2.34631 | 3.98201 |
| Q57YH1 | Tb927.8.590 | Carnitine O-palmitoyltransferase | 3.57014 | 3.97275 |
| Q382D5 | Tb927.11.14020 | RNA-binding protein | 4.00412 | 3.9513 |
| Q582W0 | Tb927.3.2470 | Pumilio/PUF RNA binding protein 8 | 3.3002 | 3.94942 |
| Q57U76 | Tb927.5.3900 | DUF4110 domain-containing protein | 3.67235 | 3.94064 |
| Q38AE4 | Tb927.10.8200 | hypothetical protein | 2.19665 | 3.90181 |
| Q4GYQ5 | Tb927.1.3180 | 40S ribosomal protein S11 | 3.92091 | 3.90132 |
| Q388L9 | Tb927.10.13800 | hypothetical protein | 3.26954 | 3.88722 |
| Q384Y9 | Tb927.11.6810 | Tubulin-tyrsoine ligase-like protein | 4.15935 | 3.8777 |
| Q57XS3 | Tb927.7.7030 | hypothetical protein | 2.42791 | 3.87299 |
| Q389K1 | Tb927.10.11390 | 60S ribosomal protein L6 | 4.05386 | 3.86197 |
| Q57XS5 | Tb927.7.7050 | hypothetical protein | 2.9185 | 3.8541 |
| Q38EE3 | Tb927.9.8200 | Pescadillo homolog | 3.53039 | 3.84321 |

### SET15 IP significantly enriched interactors (over untagged control) - continued

| UniProt ID | TriTryp ID | Description | $-log_{10}$ Student's t-test p-value | $log_2$ (tagged/untagged) |
|---|---|---|---|---|
| Q38AM9 | Tb927.10.7330 | 40S ribosomal protein S24 | 2.14687 | 3.83504 |
| Q388C7 | Tb927.10.14770 | Associated kinase of Tb14-3-3 | 3.33821 | 3.8304 |
| Q38A09 | Tb927.10.9660 | hypothetical protein | 2.81291 | 3.82221 |
| Q57ZH2 | Tb927.7.5170 | 60S ribosomal protein L23a | 4.1092 | 3.82092 |
| Q57V56 | Tb927.3.3300 | hypothetical protein | 4.00826 | 3.81542 |
| Q38FD1 | Tb927.9.4080 | DksA C4-type domain-containing protein | 3.4292 | 3.81003 |
| Q388E0 | Tb927.10.14630 | Fibrillarin | 2.85126 | 3.79445 |
| Q383J8 | Tb927.11.10810 | hypothetical protein | 3.7876 | 3.78787 |
| Q583H9 | Tb927.4.3850 | WD_REPEATS_REGION domain-containing protein | 2.31308 | 3.74813 |
| Q38EQ1 | Tb927.9.6870 | RNA-binding protein | 5.07229 | 3.73756 |
| Q388S9 | Tb927.10.13180 | Nrap protein | 4.18907 | 3.72577 |
| Q38FQ5 | Tb927.9.2650 | hypothetical protein | 4.92438 | 3.72512 |
| Q385B8 | Tb927.11.6510 | 40S ribosomal protein S21 | 3.36046 | 3.71635 |
| Q584N3 | Tb927.2.280 | Retrotransposon hot spot (RHS) protein | 2.23889 | 3.70949 |
| Q381E6 | Tb927.11.16400 | Kinetoplast DNA-associated protein | 2.7305 | 3.69274 |
| Q382F6 | Tb927.11.13810 | ATP-dependent DEAD/H RNA helicase | 4.08181 | 3.69083 |
| Q38AU3 | Tb927.10.6630 | ATP-dependent DEAD/H RNA helicase HEL64 | 2.5265 | 3.67671 |
| Q383D5 | Tb927.11.11430 | Cullin 1 | 2.98634 | 3.67634 |
| Q584T5 | Tb927.6.2210 | hypothetical protein | 2.16107 | 3.6613 |
| Q583R3 | Tb927.6.2710 | RING-type domain-containing protein | 2.38209 | 3.66077 |
| Q38CD1 | Tb927.10.1080 | 40S ribosomal protein S23 | 3.49435 | 3.64875 |
| Q383V8 | Tb927.10.13500 | Ribosomal protein | 3.54214 | 3.63036 |
| Q387J8 | Tb927.11.530 | RNA-binding protein | 3.52531 | 3.60473 |
| Q4GYR6 | Tb927.1.3070 | hypothetical protein | 3.35987 | 3.5893 |
| Q383K0 | Tb927.11.10790 | 40S ribosomal protein SA | 3.76721 | 3.57936 |
| Q38EI9 | Tb927.9.7590 | 60S ribosomal protein L11 | 3.14285 | 3.56804 |
| Q585W5 | Tb927.6.3670 | hypothetical protein | 3.20473 | 3.56031 |
| Q389G5 | Tb927.10.11760 | Pumillio RNA binding protein | 2.00216 | 3.53919 |
| Q585L0 | Tb927.4.3400 | hypothetical protein, conserved | 3.24737 | 3.53701 |
| Q38FD9 | Tb927.9.3990 | 40S ribosomal protein S7 | 4.80274 | 3.53245 |

### SET15 IP significantly enriched interactors (over untagged control) - continued

| UniProt ID | TriTryp ID | Description | $-log_{10}$ Student's t-test p-value | $log_2$ (tagged/untagged) |
|---|---|---|---|---|
| Q385Y8 | Tb927.11.4290 | 40S ribosomal protein S12 | 2.34993 | 3.48545 |
| Q587G6 | Tb927.2.2950 | hypothetical protein | 2.07158 | 3.48127 |
| Q386S5 | Tb927.11.2340 | hypothetical protein | 3.9396 | 3.4788 |
| Q388M6 | Tb927.10.13730 | 60S ribosomal protein L7 | 4.07673 | 3.46676 |
| Q386K4 | Tb927.11.3070 | WD_REPEATS_REGION domain-containing protein | 2.98843 | 3.46486 |
| Q584V1 | Tb927.6.2050 | Ribosome biogenesis regulatory protein | 3.71135 | 3.44493 |
| Q389M4 | Tb927.10.11150 | hypothetical protein | 4.66972 | 3.43146 |
| Q581A2 | Tb927.8.3820 | hypothetical protein | 3.77594 | 3.42405 |
| Q386W1 | Tb927.11.1980 | hypothetical protein | 2.99391 | 3.40884 |
| Q38D96 | Tb927.9.13280 | RNA-binding protein | 2.583 | 3.40112 |
| Q387P8 | Tb927.10.16170 | Potassium voltage-gated channel | 2.46185 | 3.39853 |
| Q585J4 | Tb927.4.3560 | Serine/threonine-protein phosphatase | 2.8735 | 3.38731 |
| Q381D1 | Tb927.11.16550 | hypothetical protein | 3.26232 | 3.38087 |
| Q584T3 | Tb927.6.2230 | hypothetical protein | 2.53357 | 3.36381 |
| Q57W29 | Tb927.5.840 | Kri1_C domain-containing protein | 3.83681 | 3.35617 |
| Q586Q9 | Tb927.2.4580 | UNC119 | 3.06884 | 3.35043 |
| Q38A77 | Tb927.10.8970 | kinetoplast-associated protein 4 isoform 2 | 3.29922 | 3.31246 |
| Q385T6 | Tb927.10.14580 | 60S ribosomal protein L17 | 4.30583 | 3.30292 |
| Q38FU7 | Tb927.9.2120 | hypothetical protein | 3.70276 | 3.30169 |
| Q38A89 | Tb927.10.8850 | hypothetical protein | 4.57183 | 3.29137 |
| Q389P5 | Tb927.10.10850 | Argonaute-like protein | 2.60182 | 3.28613 |
| Q38C06 | Tb927.10.2380 | hypothetical protein | 4.20126 | 3.28148 |
| Q381W0 | Tb927.11.14750 | PSP1 C-terminal domain-containing protein | 2.18462 | 3.27147 |
| Q57VM1 | Tb927.8.1330 | 60S ribosomal protein L7a | 3.04956 | 3.26175 |
| Q586I3 | Tb927.6.5110 | Damage-specific DNA binding protein | 3.42713 | 3.25236 |
| Q389I7 | Tb927.10.11540 | 40S ribosomal protein S3 | 3.3862 | 3.25215 |
| Q585R0 | Tb927.6.1500 | Alkyl-dihydroxyacetone phosphate synthase | 2.66972 | 3.24953 |
| Q38B59 | Tb927.10.5450 | WSD domain-containing protein | 2.16 | 3.24211 |
| Q382Z8 | Tb927.11.11820 | 40S ribosomal protein S17 | 4.03675 | 3.23999 |
| Q389Z2 | Tb927.10.9840 | Chaperone protein DNAJ | 4.18974 | 3.23137 |

### SET15 IP significantly enriched interactors (over untagged control) - continued

| UniProt ID | TriTryp ID | Description | $-log_{10}$ Student's t-test p-value | $log_2$ (tagged/untagged) |
|---|---|---|---|---|
| Q57VZ6 | Tb927.8.4390 | Translation initiation factor eIF-2B beta subunit | 2.70249 | 3.21469 |
| Q38DT1 | Tb927.9.10920 | kinetoplastid kinetochore protein 3 (KKT3) | 2.52493 | 3.20547 |
| Q38AL9 | Tb927.10.7440 | ATP-dependent DEAD/H RNA helicase | 2.98255 | 3.20171 |
| Q387E2 | Tb927.11.1080 | hypothetical protein | 3.9101 | 3.19574 |
| Q57XC9 | Tb927.8.2600 | WD_REPEATS_REGION domain-containing protein | 3.01492 | 3.17189 |
| Q386F2 | Tb927.11.3590 | 40S ribosomal protein S4 | 3.41196 | 3.14524 |
| Q38D68 | Tb927.9.13610 | Helicase | 2.63388 | 3.14487 |
| Q386Y8 | Tb927.11.1660 | vesicular transport protein (CDC48 homologue) | 2.93632 | 3.13838 |
| Q57YK4 | Tb927.8.6030 | 60S ribosomal protein L12 | 3.90584 | 3.11987 |
| Q57W53 | Tb927.5.1080 | RRM domain-containing protein | 3.11179 | 3.08933 |
| Q57UC8 | Tb927.7.3810 | CSD domain-containing protein | 2.64629 | 3.08698 |
| Q384B1 | Tb927.11.9170 | DNA topoisomerase | 5.00839 | 3.08673 |
| Q580Z5 | Tb927.8.3750 | Nucleolar protein | 4.27428 | 3.08103 |
| Q389D3 | Tb927.10.12100 | RNA-binding protein 7B | 2.23506 | 3.06525 |
| Q386V3 | Tb927.11.2050 | 60S acidic ribosomal protein P0 | 2.89859 | 3.04556 |
| Q38A41 | Tb927.10.9330 | hypothetical protein | 2.63072 | 3.03462 |
| Q385C6 | Tb927.11.6430 | hypothetical protein | 2.0276 | 3.02604 |
| Q57VV0 | Tb927.7.1040 | 40S ribosomal protein S16 | 3.21614 | 3.01444 |
| Q388D3 | Tb927.10.14600 | 40S ribosomal protein S2 | 3.31631 | 3.01003 |
| Q38B16 | Tb927.10.5880 | hypothetical protein | 2.28198 | 2.98036 |
| Q38AW9 | Tb927.10.220 | 60S ribosomal protein L37a (60S ribosomal proteins L37) | 2.11727 | 2.97034 |
| Q57UM7 | Tb927.7.3970 | hypothetical protein | 2.33563 | 2.9664 |
| Q381T5 | Tb927.11.15000 | survival of motor neuron (SMN)-like protein | 2.0564 | 2.96548 |
| Q57XX0 | Tb927.10.2840 | 40S ribosomal protein S25 | 2.95092 | 2.95547 |
| Q580A5 | Tb927.4.770 | Ubiquitin-protein ligase | 2.04869 | 2.94927 |
| Q384Z8 | Tb927.11.6710 | TPR_REGION domain-containing protein | 2.79763 | 2.93361 |
| Q57VV2 | Tb927.7.1060 | hypothetical protein | 2.26167 | 2.92866 |
| Q385E9 | Tb927.11.6180 | 60S ribosomal protein L28 | 3.40982 | 2.92177 |
| Q385D9 | Tb927.11.6300 | 40S ribosomal protein S5 | 3.48865 | 2.91507 |
| Q585U7 | Tb927.2.5240 | WD_REPEATS_REGION domain-containing protein | 2.80848 | 2.91294 |

### SET15 IP significantly enriched interactors (over untagged control) - continued

| UniProt ID | TriTryp ID | Description | $-\log_{10}$ Student's t-test p-value | $\log_2$ (tagged/untagged) |
|---|---|---|---|---|
| Q57VG4 | Tb927.5.760 | PSP1 C-terminal domain-containing protein | 2.97773 | 2.9101 |
| Q57XN5 | Tb927.7.2390 | hypothetical protein | 3.82771 | 2.90552 |
| Q8IFI8 | Tb927.1.120 | retrotransposon hot spot protein 4 (RHS4) | 3.52319 | 2.891 |
| Q381S8 | Tb927.11.15070 | hypothetical protein | 2.97025 | 2.88682 |
| Q38B69 | Tb927.10.5330 | 40S ribosomal protein S18 | 3.16758 | 2.8793 |
| Q385Z8 | Tb927.11.4180 | PSP1 C-terminal domain-containing protein | 3.50981 | 2.83662 |
| Q57XN5 | Tb927.7.2390 | hypothetical protein | 3.82771 | 2.90552 |
| Q8IFI8 | Tb927.1.120 | retrotransposon hot spot protein 4 (RHS4) | 3.52319 | 2.891 |
| Q381S8 | Tb927.11.15070 | hypothetical protein | 2.97025 | 2.88682 |
| Q38B69 | Tb927.10.5330 | 40S ribosomal protein S18 | 3.16758 | 2.8793 |
| Q385Z8 | Tb927.11.4180 | PSP1 C-terminal domain-containing protein | 3.50981 | 2.83662 |
| Q388B1 | Tb927.10.14950 | RNA binding protein | 2.10442 | 2.83194 |
| Q38FQ8 | Tb927.9.2620 | DUF4460 domain-containing protein | 4.01138 | 2.80817 |
| Q38D98 | Tb927.9.13240 | hypothetical protein | 2.32729 | 2.80118 |
| Q57WA3 | Tb927.3.3670 | RNA-binding protein | 2.91958 | 2.79741 |
| Q4GZ47 | Tb927.1.1720 | PPIase cyclophilin-type domain-containing protein | 3.71482 | 2.79029 |
| Q38A78 | Tb927.10.8960 | hypothetical protein | 3.22946 | 2.78949 |
| Q57VM8 | Tb927.8.1270 | hypothetical protein | 3.17046 | 2.76994 |
| Q38BK6 | Tb927.10.3940 | 40S ribosomal protein S3a | 3.43681 | 2.75714 |
| Q38A85 | Tb927.10.8890 | Kinetoplast DNA-associated protein | 2.37494 | 2.75642 |
| Q57V04 | Tb927.7.2070 | heat shock protein DNAJ | 4.20609 | 2.74908 |
| Q38DS7 | Tb927.9.10960 | ATP-dependent DEAD/H RNA helicase | 4.87165 | 2.74222 |
| Q57ZC0 | Tb927.3.1910 | SLBP_RNA_bind domain-containing protein | 2.27418 | 2.736 |
| Q385U8 | Tb927.11.4690 | Mitochondrial DNA polymerase I protein B | 3.65971 | 2.70458 |
| Q387Q3 | Tb927.10.16120 | inosine-5'-monophosphate dehydrogenase | 4.43817 | 2.704 |
| Q57Z68 | Tb927.5.3250.1 | WW domain-containing protein | 2.58578 | 2.69314 |
| Q38BJ4 | Tb927.10.4060 | ANK_REP_REGION domain-containing protein | 2.09685 | 2.68063 |
| Q57ZI6 | Tb927.7.5030 | hypothetical protein | 2.01053 | 2.67893 |
| Q586R7 | Tb927.2.4710 | RNA-binding protein | 3.08724 | 2.67873 |
| Q585W7 | Tb927.6.3690 | Pre-mRNA cleavage complex II Clp1-like, conserved | 2.40348 | 2.65934 |

### SET15 IP significantly enriched interactors (over untagged control) - continued

| UniProt ID | TriTryp ID | Description | $-\log_{10}$ Student's t-test p-value | $\log_2$ (tagged/untagged) |
|---|---|---|---|---|
| Q38C70 | Tb927.10.1720 | ATP-dependent DEAD/H RNA helicase | 2.50131 | 2.65842 |
| Q38CY8 | Tb927.9.15130 | 60S ribosomal protein L5 | 3.21633 | 2.65776 |
| Q388B3 | Tb927.10.14930 | RNA binding protein | 3.65408 | 2.63675 |
| Q580I2 | Tb927.8.1410 | hypothetical protein | 2.1087 | 2.61964 |
| Q38BQ9 | Tb927.10.3370 | 60S acidic ribosomal protein P2 | 3.24002 | 2.61218 |
| Q585A4 | Tb927.6.1430 | MATH domain-containing protein | 2.69023 | 2.57222 |
| Q389Z8 | Tb927.10.9780 | ATP-dependent DEAD/H RNA helicase | 3.06332 | 2.56546 |
| Q583H8 | Tb927.4.3840 | Nucleolar protein | 2.7113 | 2.53828 |
| Q586C2 | Tb927.10.15120 | 40S ribosomal protein S13 | 2.65386 | 2.53534 |
| Q383R3 | Tb927.11.10160 | 60S ribosomal protein L22 | 4.02774 | 2.53296 |
| Q580V1 | Tb927.3.5240 | hypothetical protein | 2.70324 | 2.53034 |
| Q38F23 | Tb927.9.5320 | Nucleolar RNA binding protein | 4.15374 | 2.51347 |
| Q38EY6 | Tb927.9.5690 | 60S acidic ribosomal protein | 2.95999 | 2.51149 |
| Q581X8 | Tb927.8.2040 | DUF1767 domain-containing protein | 2.13478 | 2.50631 |
| Q586U9 | Tb927.2.4280 | Dual specificity protein phosphatase | 2.03869 | 2.50609 |
| Q57VA4 | Tb927.5.1590 | hypothetical protein | 2.9407 | 2.48611 |
| Q57XW2 | Tb927.3.1290 | Cullin 4B | 2.13664 | 2.47057 |
| Q57YB5 | Tb927.7.3050 | hypothetical protein | 2.40919 | 2.4608 |
| Q383P6 | Tb927.11.10330 | hypothetical protein | 2.75613 | 2.4545 |
| Q38BC7 | Tb927.10.4740 | Nucleolar RNA-binding protein | 3.05298 | 2.45369 |
| Q4GYF0 | Tb927.1.4280 | hypothetical protein | 2.53048 | 2.44099 |
| D6XDN4 | Tb927.3.3270 | ATP-dependent 6-phosphofructokinase (ATP-PFK) | 3.33743 | 2.43232 |
| Q385X2 | Tb927.11.4450 | Alba domain-containing protein | 3.38772 | 2.40524 |
| Q38AC1 | Tb927.10.8430 | 40S ribosomal protein S12 | 3.25531 | 2.39142 |
| Q387J6 | Tb927.11.550 | hypothetical protein | 3.25812 | 2.38947 |
| Q57VZ0 | Tb927.8.4450 | RNA-binding protein | 2.37816 | 2.37347 |
| Q388E8 | Tb927.10.14550 | ATP-dependent DEAD/H RNA helicase | 3.58184 | 2.3714 |
| Q38C52 | Tb927.10.1900 | DNA topoisomerase IA | 3.48388 | 2.36735 |
| Q57YW0 | Tb927.8.3530 | glycerol-3-phosphate dehydrogenase [NAD+], glycosomal | 4.54295 | 2.36698 |
| Q385D7 | Tb927.11.6320 | C3H1-type domain-containing protein | 2.67242 | 2.36186 |

### SET15 IP significantly enriched interactors (over untagged control) - continued

| UniProt ID | TriTryp ID | Description | $-\log_{10}$ Student's t-test p-value | $\log_2$ (tagged/untagged) |
|---|---|---|---|---|
| Q38EH3 | Tb927.9.7820 | hypothetical protein | 3.86725 | 2.35923 |
| Q38AF8 | Tb927.10.8060 | SET domain-containing protein (SET8) | 2.31375 | 2.34632 |
| Q383B0 | Tb927.11.11700 | NTP_transferase domain-containing protein | 4.16059 | 2.3409 |
| Q580W2 | Tb927.3.5130 | Exonuclease domain-containing protein | 2.9251 | 2.33829 |
| Q383C3 | Tb927.11.11570 | hypothetical protein | 2.7362 | 2.32769 |
| Q57ZJ9 | Tb927.3.3130 | SAM domain-containing protein | 2.43051 | 2.31054 |
| Q4FKA6 | Tb11.1400.1 | Translation initiation factor eIF2B delta subunit | 2.3716 | 2.30415 |
| Q4FKG9 | Tb11.0290 | 40S ribosomal protein S14 (40s ribosomal protein S14) | 2.61375 | 2.2664 |
| Q38B66 | Tb927.10.5370 | 40S ribosomal protein S10 | 2.71203 | 2.26611 |
| Q38AD1 | Tb927.10.8330 | PSP1 C-terminal domain-containing protein | 2.0349 | 2.22758 |
| Q57WI5 | Tb927.7.3700 | hypothetical protein | 2.61439 | 2.21652 |
| Q38AY4 | Tb927.10.6220 | 5'-3' exoribonuclease XRND | 2.03223 | 2.21397 |
| Q387C7 | Tb927.11.1250 | hypothetical protein | 4.40491 | 2.17145 |
| Q389R6 | Tb927.10.10620 | hypothetical protein | 2.9099 | 2.1467 |
| Q386S9 | Tb927.11.2300 | Eukaryotic peptide chain release factor subunit 1 | 2.22266 | 2.11068 |
| Q38B00 | Tb927.10.6060 | Universal minicircle sequence binding protein (UMSBP) | 3.96851 | 2.08543 |
| Q57XY3 | Tb927.3.1500 | Variant surface glycoprotein (VSG)-related | 3.01641 | 2.08463 |
| Q584U2 | Tb927.6.2140 | hypothetical protein | 3.74363 | 2.07661 |
| Q385I3 | Tb927.11.5850 | RNA-binding protein | 2.33252 | 2.06997 |
| Q586P0 | Tb927.2.6070 | hypothetical protein | 2.23715 | 2.05721 |
| Q386Q9 | Tb927.11.2510 | NUC173 domain-containing protein | 3.22904 | 2.05103 |
| Q57WY4 | Tb927.3.1010 | hypothetical protein | 3.32166 | 2.01911 |

### SET20 IP significantly enriched interactors (over untagged control)

| UniProt ID | TriTryp ID | Description | $-log_{10}$ Student's t-test p-value | $log_2$ (tagged/untagged) |
|---|---|---|---|---|
| Q57XE0 | Tb927.8.2490 | SET20 | 5.95491 | 9.11766 |
| Q38C15 | Tb927.10.2290 | chaperone protein DnaJ, putative | 3.36358 | 2.56159 |
| Q38DH5 | Tb927.9.12300 | replication factor C, subunit 3, putative | 2.85488 | 2.17278 |

### SET23 IP significantly enriched interactors (over untagged control)

| UniProt ID | TriTryp ID | Description | $-\log_{10}$ Student's t-test p-value | $\log_2$ (tagged/untagged) |
|---|---|---|---|---|
| Q57YP8 | Tb927.8.6470 | SET23 | 5.85826 | 13.5365 |
| Q57UQ9 | Tb927.7.4290 | Nuclear distribution protein C homolog | 5.5979 | 7.94808 |
| Q382B5 | Tb927.11.14220 | Regulator of ESAG9-1 | 3.87279 | 3.76776 |
| Q38EI4 | Tb927.9.7690 | SUMO-interacting motif-containing protein | 3.8291 | 3.70809 |
| Q586Q9 | Tb927.2.4580 | UNC119 | 3.28072 | 3.15086 |
| Q57XP8 | Tb927.7.2260 | SEP domain containing protein, putative | 2.7166 | 2.00686 |

### SET25 IP significantly enriched interactors (over untagged control)

| UniProt ID | TriTryp ID | Description | $-log_{10}$ Student's t-test p-value | $log_2$ (tagged/untagged) |
|---|---|---|---|---|
| Q38FZ5 | Tb927.9.1510 | SET25 | 5.11563 | 12.386 |

### Table S6. Homology detected for proteins enriched with candidate chromatin regulators

Identified interactors shared between two or more proteins are presented as a network followed by those that are uniquely associated with each protein in that network.

[1] RIT-seq data from Alsford et al (2011). Scoring matrix key: 0-0-0-0 group - significant loss of fitness in all four experiments; 0-0-1-0 group - significant loss of fitness in BF experiments only; 1-1-0-0 group - significant loss of fitness in PF experiments only; 1-1-1-0 group - significant loss of fitness in the differentiation (BF to PF) experiment only; 1-1-1-1 group - no significant loss of fitness in any experiment; none of the above - other groups.

[2] The estimated probability of the template to be (at least partly) homologous to a query sequence is the key criterion for deciding whether a template HMM is actually homologous or just a high-scoring chance hit. Scores >95%: homology is nearly certain. Serious consideration should be given to hits whenever (1) it has >50% probability, or (2) it has >30% probability and is among the top three hits (Soding et al., 2005). Detected homologies were inspected manually and, where relevant, hits were checked by specific BLAST searches with the same *T. brucei* protein against all *S. cerevisiae* or *S. pombe* proteins.

Proteins marked as nuclear enriched by mass spectrometry were analyzed by Goos et al. (2017).

NRM – no relevant match

**CRD1-SET27 network**

| UniProt ID | TriTryp ID | Gene name/description | [1]RIT-seq scoring matrix | Localization (BF) | PDB entry | HHpred domain | [2]HHpred domain probability % |
|---|---|---|---|---|---|---|---|
| Q57X70 | Tb11.v5.0267 | CRD1 | 1-1-1-1 | nuclear (this study) | 4O42_A | Chromodomain CHD1 | 97.38 |
| Q38D80 | Tb927.9.13470 | SET27 | 1-1-1-1 | nuclear/cytoplasmic (this study); nuclear enriched (mass spectrometry) | 5JLB_A | Histone lysine N-methyltransferase | 99.06 |
| Q4GYF3 | Tb927.1.4250 | uncharacterized | 1-1-1-1 | nuclear enriched (mass spectrometry) | NRM | NRM | NRM |
| Q382F5 | Tb927.11.13820 | uncharacterized | other | nuclear enriched (mass spectrometry) | NRM | NRM | NRM |
| Q582U8 | Tb927.3.2350 | uncharacterized; similar to Bye1 (*S. cerevisiae*) | other | - | 5XFO_A | PHD Finger domain | 97.18 |
| Q382Z7 | Tb927.11.11840 | uncharacterized; similar to Chp2 (*S. pombe*) | 1-1-1-1 | - | 6FTO_B | Chromoshadow domain | 94.75 |
| D6XIZ6 | Tb927.7.4650 | JBP2 | 1-1-1-1 | - | 4LT5_A<br>6G7E_B | Tet-like Dioxygenase;<br>ATPase Helicase | 100<br>97.97 |
| Q586X9 | Tb927.2.3580 | TFIIS2-1; also detected with EAF6 & TFIIS2-2 | 1-1-1-1 | nuclear enriched (mass spectrometry) | 1PQV_S<br>2M1H_A | TFIIS;<br>PWWP domain | 100<br>99.22 |

### Enriched with YFP-CRD1 affinity selections

| UniProt ID | TriTryp ID | Gene name/description | [1]RIT-seq scoring matrix | Localization (BF) | PDB entry | HHpred domain | [2]HHpred domain probability % |
|---|---|---|---|---|---|---|---|
| Q38FV1 | Tb927.9.2070 | uncharacterized; also detected with HDAC1 | other | - | NRM | NRM | NRM |
| Q387L4 | Tb927.11.370 | RAP1; also detected with HDAC3 & TRF | 0-0-1-0 | nuclear enriched (mass spectrometry) | 3AL2_A | BRCT domain | 94.26 |
| Q57YU2 | Tb927.8.7400 | RPB1; also detected with SET26 & TFIIS2-2 | 0-0-0-0 | nuclear enriched (mass spectrometry) | 4XPZ_A | Rpb1 RNAPII subunit | 96.21 |
| Q580S5 | Tb927.3.5500 | RPB3; also detcted with SET26 & TFIIS2-2 | other | nuclear enriched (mass spectrometry) | 1TWF_C | Rpb3 RNAPII subunit | 100 |

### Enriched in YFP-SET27 affinity selections

| UniProt ID | TriTryp ID | Gene name/description | [1]RIT-seq scoring matrix | Localization (BF) | PDB entry | HHpred domain | [2]HHpred domain probability % |
|---|---|---|---|---|---|---|---|
| Q38BT9 | Tb927.10.3070 | uncharacterized | other | nuclear enriched (mass spectrometry) | 4QQ6_A | Tudor Domain | 47.7 |
| Q57Z72 | Tb927.5.3210 | SUMO; also detected with BDF3, BDF6 & HAT2 | other | - | 3PGE_A | SUMO (small ubiquitin like protein) | 99.03 |
| Q38EP6 | Tb927.9.6920 | uncharacterized | 1-1-1-1 | - | NRM | NRM | NRM |
| P86938 | Tb927.11.13640 | JPB1 | 1-1-1-1 | - | 2XSE_A<br>5DEU_A | Thymine Dioxygenase;<br>Tet-like | 100<br>96.4 |
| Q389E6 | Tb927.10.11960 | PNUTS; also detected with TFIIS2-2 | 1-1-1-1 | - | NRM | NRM | NRM |
| Q57W24 | Tb927.5.790 | Casein kinase I isoform 1 | other | - | 6GZD_A | Casein Kinase 1 | 100 |
| Q57VC7 | Tb927.5.1360 | NDRT; also detected with BDF4 | 1-1-1-1 | - | 2F62_B | Nucleoside 2-deoxyribosyltransferase | 99.87 |
| Q586H7 | Tb927.6.5050 | V-type ATPase | 1-1-1-1 | - | 6O7U_c | V-type proton ATPase subunit | 99.95 |
| Q388E9 | Tb927.10.14530 | RPN8 | other | - | 6EPF | 26S Proteasome subunit | 100 |
| Q57UB5 | Tb927.5.3510 | SMC3; also detected with HDAC3 | other | nuclear enriched (mass spectrometry) | 6WG3_B | SMC Cohesin/Condensin | 100 |
| Q38E40 | Tb927.9.9550 | ERBP2 | 0-0-1-0 | - | NRM | NRM | NRM |
| Q38C86 | Tb927.10.1550 | RPN5 | 0-0-1-0 | - | 3JCK_B | 26S Proteasome subunit | 100 |

### BDF1-BDF4 network

| UniProt ID | TriTryp ID | Gene name/description | [1]RIT-seq scoring matrix | Localization (BF) | PDB entry | HHpred domain | [2]HHpred domain probability % |
|---|---|---|---|---|---|---|---|
| Q38AE9 | Tb927.10.8150 | BDF1 | 1-1-1-1 | nuclear (this study); nuclear enriched (mass spectrometry) | 6DNE_B<br>2NCZ_A | Bromo domain;<br>ET Domain | 99.0<br>97.4 |
| Q57UR8 | Tb927.7.4380 | BDF4 | other | nuclear/cytoplasmic (this study) | 6DNE_B<br>2E61_A | Bromo domain;<br>Znf-CW PWWP | 97.8<br>93.24 |
| Q383J7 | Tb927.11.10820 | Casein kinase II subunit beta;<br>also detected with BDF7 | other | - | 4DGL_A | Casein Kinase II subunit beta | 100 |

### Enriched in YFP-BDF1 affinity selections

| UniProt ID | TriTryp ID | Gene name/description | [1]RIT-seq scoring matrix | Localization (BF) | PDB entry | HHpred domain | [2]HHpred domain probability % |
|---|---|---|---|---|---|---|---|
| Q389A2 | Tb927.10.12410 | uncharacterized | other | - | 1G0S_A | ADP-ribose pyrophosphatase | 99.72 |
| Q57WL5 | Tb927.8.8150 | C2 domain;<br>also detected with SET26 | 0-0-0-0 | - | 6FLJ_A | C2 domain (Calcium binding, membrane bound) | 99.2 |

### Enriched in YFP-BDF4 affinity selections

| UniProt ID | TriTryp ID | Gene name/description | [1]RIT-seq scoring matrix | Localization (BF) | PDB entry | HHpred domain | [2]HHpred domain probability % |
|---|---|---|---|---|---|---|---|
| Q38D21 | Tb927.9.14430 | Casein kinase II; also detected with BDF7, SET26 & ZCW1 | other | - | 5XVU_B | Casein Kinase CKII | 100 |
| Q57YY8 | Tb927.8.3250 | Dynein Heavy Chain | 0-0-0-0 | flagellar (Dean et al., 2016) | 6SC2_B | Dynein Heavy Chain | 100 |
| Q57V45 | Tb927.3.3410 | Aspartyl amino peptidase | 1-1-1-0 | - | 4EME_A | Aspartyl Amino peptidase | 100 |
| Q57VC7 | Tb927.5.1360 | NDRT | 1-1-1-1 | - | 2F62_B | Nucleoside 2-deoxyribosyltransferase | 99.87 |
| Q57UC1 | Tb927.5.3450 | eIF2A | 1-1-1-0 | - | 3WJ9_A | Eukaryotic translation initiation factor 2A | 100 |

### BDF3-BDF5-HAT2 network

| UniProt ID | TriTryp ID | Gene name/description | [1]RIT-seq scoring matrix | Localization (BF) | PDB entry | HHpred domain | [2]HHpred domain probability % |
|---|---|---|---|---|---|---|---|
| Q383S2 | Tb927.11.10070 | BDF3 | other | nuclear (this study); nuclear enriched (mass spectrometry) | 4QB3_A<br>2NCZ_A | Bromo domain;<br>ET Domain | 99.11<br>98.72 |
| Q382J7 | Tb927.11.13400 | BDF5; also detected with ZCW1 | 0-0-1-0 | nuclear (this study); nuclear enriched (mass spectrometry) | 2R0Y_A<br>2F5J_B | Tandem Bromo Domains;<br>MRG Domain | 99.7<br>96.87 |
| Q383C5 | Tb927.11.11530 | HAT2; similar to KAT8 (*H. sapiens*) and Esa1 (*S. cerevisiae*) | 1-1-1-1 | nuclear (this study); nuclear enriched (mass spectrometry) | 3TO7_A | Esa1 Histone acetyltransferase | 100 |
| Q4FKD8 | Tb927.11.18680 | Dynein Light Chain | other | - | 5E0L_A | Dynein Light Chain | 99.97 |
| Q580M8 | Tb927.3.4140 | uncharacterized | other | - | 4ZZZ_B | Poly ADP Ribose Polymerase (PARP) | 57.11 |
| Q584F3 | Tb927.4.2340 | uncharacterized | 0-0-0-0 | nuclear enriched (mass spectrometry) | 5FUR_G | Bromo domain (TFIID-TFIIA) | 78.45 |
| Q585E0 | Tb927.6.1070 | uncharacterized | other | nuclear enriched (mass spectrometry) | NRM | NRM | NRM |
| Q57Y87 | Tb927.7.2770 | uncharacterized | other | nuclear enriched (mass spectrometry) | NRM | NRM | NRM |
| Q38D92 | Tb927.9.13320 | uncharacterized; also detected with SET26 | other | nuclear enriched (mass spectrometry); | 5XZW_A | FHA domain (Phospho-Threonine-Binding) | 96.3 |
| Q385P5 | Tb927.11.5230 | uncharacterized; also detected with HDAC3 & ZCW1 | 0-0-0-0 | nuclear (D'Archivio and Wickstead, 2017); nuclear enriched (mass spectrometry) | 1UZ3_A | EMSY ENT domain (Chromo shadow binding) | 95.93 |

### Enriched in YFP-BDF3 affinity selections

| UniProt ID | TriTryp ID | Gene name/description | [1]RIT-seq scoring matrix | Localization (BF) | PDB entry | HHpred domain | [2]HHpred domain probability % |
|---|---|---|---|---|---|---|---|
| Q57Z72 | Tb927.5.3210 | SUMO;<br>also detected with BDF6, HAT2 & SET27 | other | - | 3PGE_A | SUMO (Small Ubiquitin like protein) | 99.03 |
| Q38BR9 | Tb927.10.3280 | L38;<br>also detected with BDF5 | 0-0-0-0 | - | 5UMD_d | 60S ribosomal protein | 99.94 |
| Q383E9 | Tb927.11.11290 | HSP70;<br>also detected with BDF5, BDF7, SET26, ZCW1, TFIIS2-2 & DOT1A | 1-1-1-0 | nuclear enriched (mass spectrometry) | 5FPN_A | Heat Shock 70 protein | 100 |
| Q4GYZ8 | Tb927.1.2230 | SMP1-1;<br>also detected with BDF5 | 1-1-1-1 | - | 2FE0_A | Small myristoylated protein 1 | 100 |

### Enriched in YFP-BDF5 affinity selections

| UniProt ID | TriTryp ID | Gene name/description | [1]RIT-seq scoring matrix | Localization (BF) | PDB entry | HHpred domain | [2]HHpred domain probability % |
|---|---|---|---|---|---|---|---|
| Q38BR9 | Tb927.10.3280 | L38;<br>also detected with BDF3 | 0-0-0-0 | - | 5UMD_d | 60S ribosomal protein | 99.94 |
| Q383E9 | Tb927.11.11290 | HSP70;<br>also detected with BDF3, BDF7, SET26, ZCW1, TFIIS2-2 & DOT1A | 1-1-1-0 | nuclear enriched (mass spectrometry) | 2FE0_A | Heat Shock 70 protein | 100 |
| Q4GYZ8 | Tb927.1.2230 | SMP1-1;<br>also detected with BDF3 | 1-1-1-1 | - | 5FPN_A | Small myristoylated protein 1 | 100 |

### Enriched in YFP-HAT2 affinity selections

| UniProt ID | TriTryp ID | Gene name/description | [1]RIT-seq scoring matrix | Localization (BF) | PDB entry | HHpred domain | [2]HHpred domain probability % |
|---|---|---|---|---|---|---|---|
| Q57Z72 | Tb927.5.3210 | SUMO;<br>also detected with BDF3, BDF6 & SET27 | other | - | 3PGE_A | SUMO (Small Ubiquitin like protein) | 99.03 |
| Q38BI9 | Tb927.10.4110 | L30 | other | - | 5UMD_6 | 60S Ribosomal protein | 99.87 |
| Q9N942 | Tb927.1.420 | RHS5 | - | - | 6K0R_B | RuvB-like AAA-ATPase | 96.11 |
| Q38BE4 | Tb927.10.4570 | Elongation factor 2 | 0-0-0-0 | - | 6ID1_C | Splicing factor | 100 |

**BDF2-HDAC3 network**

| UniProt ID | TriTryp ID | Gene name/description | [1]RIT-seq scoring matrix | Localization (BF) | PDB entry | HHpred domain | [2]HHpred domain probability % |
|---|---|---|---|---|---|---|---|
| Q38AM1 | Tb927.10.7420 | BDF2;<br>similar to Bdf1<br>(*S. cerevisiae*);<br>also detected with ZCW1 | 1-1-1-1 | nuclear (this study) | 6DNE_B<br>2ZD7_A | Bromo domain;<br>NAP domain (histone chaperone) | 99.29<br>83.7 |
| Q586J9 | Tb927.2.2190 | HDAC3;<br>also detected with TRF & ZCW1 | 0-0-0-0 | nuclear (this study);<br>nuclear (Wang et al., 2010) | 2VQM_A | HDAC | 100 |
| Q582V9 | Tb927.3.2460 | uncharacterized | 0-0-0-0 | - | 5IJO_N | Nuclear pore (Nup155) | 56.06 |
| Q586Z9 | Tb927.6.4330 | uncharacterized;<br>also detected with TRF | 0-0-0-0 | nuclear enriched (mass spectrometry);<br>telomeric (Reis et al., 2018) | NRM | NRM | NRM |
| Q38FD7 | Tb927.9.4000 | uncharacterized;<br>also detected with TRF | - | nuclear enriched (mass spectrometry);<br>telomeric (Reis et al., 2018) | NRM | NRM | NRM |
| Q38EB9 | Tb927.9.8520 | uncharacterized | other | - | 5Y27_A | DNA pol epsilon | 91.92 |
| Q388W1 | Tb927.10.12850 | TRF; telomere-repeat bining protein | 0-0-0-0 | nuclear (this study);<br>nuclear enriched (mass spectrometry);<br>telomeric (Jehi et al., 2014; Li et al., 2005) | 1GV2_A | MYB domain DNA binding | 80.11 |
| Q57XY9 | Tb927.3.1560 | TIF2;<br>also detected with TRF | other | nuclear;<br>telomeric (Jehi et al., 2014) | NRM | NRM | NRM |
| Q383U2 | Tb927.11.9870 | TelAP1;<br>also detected with TRF | 1-1-1-1 | nuclear enriched (mass spectrometry);<br>telomeric (Reis et al., 2018) | 4B0Z_A | Proteasome subunit | 56.61 |
| Q385L3 | Tb927.11.5550 | DNA polymerase theta (POLQ);<br>also detected with TRF | 1-1-1-1 | nuclear enriched (mass spectrometry);<br>telomeric (Reis et al., 2018) | 4X0Q_B | DNA polymerase theta | 100 |
| Q38BZ3 | Tb927.10.2520 | PrimPol-like protein 2;<br>also detected with TRF | 0-0-1-0 | nuclear enriched (mass spectrometry) | 5L2X_B | DNA Primase Polymerase | 100 |
| Q57W25 | Tb927.5.800 | Casein kinase I, isoform 2 | - | - | 5CYZ_A | Casein kinase I homolog HRR25 | 100 |
| Q38D76 | Tb927.9.13510 | PARN3 | 1-1-1-1 | - | 3D45_B | Poly(A)-specific ribonuclease PARN | 100 |
| Q57WS6 | Tb927.7.1240 | Sphingosine kinase | other | - | 4WER_A | Diacylglycerol/lipid kinase | 100 |

### Enriched in YFP-BDF2 affinity selections

| UniProt ID | TriTryp ID | Gene name/description | [1]RIT-seq scoring matrix | Localization (BF) | PDB entry | HHpred domain | [2]HHpred domain probability % |
|---|---|---|---|---|---|---|---|
| Q57UN4 | Tb927.7.4040 | Similar to Swc4 (*S. cerevisiae*); also detected with ZCW1 | other | nuclear enriched (mass spectrometry) | 4IEJ_A | DMAP1, chromatin regulator; SWR1 complex | 97.52 |
| Q57ZL0 | Tb927.3.3020 | actin-related; similar to Arp6 (*S. cerevisiae*); also detected with ZCW1 | other | nuclear enriched (mass spectrometry) | 4FO0_A | Actin-related protein Chromatin remodelling | 100 |
| Q584P9 | Tb927.6.2570 | actin-related; also detected with ZCW1 | other | nuclear enriched (mass spectrometry) | 4I6M_B | Actin-related protein Chromatin remodelling | 97.36 |
| Q383K6 | Tb927.11.10730 | SWI/SNF-related helicase; similar to Swr1 (*S. cerevisiae*); also detected with ZCW1 | other | nuclear enriched (mass spectrometry) | 6FML_G | ATPase Helicase, SWR1/INO80 complexes | 100 |
| Q389H2 | Tb927.10.11690 | YEATS; similar to YEA1; Yaf9 (*S. cerevisiae*); also detected with ZCW1 | other | nuclear enriched (mass spectrometry) | 6T1I_A | YEATS domain Acetyl-/Crontonyl-lysine histone binding | 98.21 |
| Q57YH0 | Tb927.8.600 | Bucentaur/craniofacial development; also detected with ZCW1 | other | nuclear enriched (mass spectrometry) | 6PX3_S | Histone/nucleosome binding (possible *S. cerevisiae* equivalent SWR1-C Swc5) | 53.46 |
| Q389D3 | Tb927.10.12100 | RBP7B; stumpy driver (Mony et al., 2014) | other | - | 5YVG_Y | RRM RNA binding | 99.17 |
| Q385T8 | Tb927.11.4800 | clathrin coat assembly protein | other | - | 5OO7_B | Phospholipid binding | 100 |
| Q583J3 | Tb927.4.2000 | ruvB-like DNA helicase; similar to Rvb1/2 (*S. cerevisiae*); also detected with ZCW1 | 0-0-1-0 | - | 6H7X_A | RuvB-like ATPase helicase, SWR1/INO80 complexes | 99.96 |
| Q580C6 | Tb927.4.980 | actin-related; similar to Act1 (*S. cerevisiae*); also detected with ZCW1 | 1-1-1-1 | nuclear enriched (mass spectrometry) | 4AM6_A | Actin or Actin-related protein, Chromatin remodelling | 100 |
| Q38C43 | Tb927.10.2000 | actin-related; similar to Arp4 (*S. cerevisiae*); also detected with ZCW1 | other | nuclear enriched (mass spectrometry) | 6GEN_R | Actin-related protein, Chromatin remodelling | 100 |
| Q581V4 | Tb927.4.1270 | ruvB-like DNA helicase; similar to Rvb1/2 (*S. cerevisiae*); also detected with ZCW1 | 0-0-1-0 | - | 2C9O_C | RuvB-like ATPase helicase; SWR1/INO80 complexes | 100 |
| Q385E0 | Tb927.11.6290 | HIT zinc finger; similar to Swc6 (*S. cerevisiae*); also detected with ZCW1 | 1-1-1-1 | nuclear enriched (mass spectrometry) | 2N95_A<br>6GEJ_S | HIT Zinc Finger; SWR/INO80 complex linked to RuvB | 97.12<br>99.4 |
| Q57YW0 | Tb927.8.3530 | G3PDH | 0-0-0-0 | glycosomal | 1EVY_A | Glycerol-3-phosphate dehydrogenase | 100 |
| Q389G9 | Tb927.10.11720 | ZCW1 | other | nuclear (this study); nuclear enriched (mass spectrometry) | 6O5W_A | MORC family CW-type zinc finger | 97.53 |
| Q57ZS8 | Tb927.5.2090 | kinesin | 0-0-1-0 | - | 5MIO_C<br>6YUF_A | Kinesin; Cohesin Rad21 | 100<br>99.9 |
| Q389D9 | Tb927.10.12030 | uncharacterized | 1-1-1-1 | nuclear enriched (mass spectrometry) | 4AN6_A | Protease inhibitor | 28.6 |

### Enriched in YFP-BDF2 affinity selections – continued

| UniProt ID | TriTryp ID | Gene name/description | [1]RIT-seq scoring matrix | Localization (BF) | PDB entry | HHpred domain | [2]HHpred domain probability % |
|---|---|---|---|---|---|---|---|
| Q384I8 | Tb927.11.8310 | CITFA-4 | 1-1-1-0 | nuclear enriched (mass spectrometry) | 1Q1V_A | Winged Helix-related DNA binding domain | 65.99 |
| Q385A8 | Tb927.11.6610 | uncharacterized | 1-1-1-1 | - | 2VOO_B<br>4WKR_A | La NTD RNA binding;<br>La RNA recognition Motif | 99.96<br>99.85 |
| Q387B3 | Tb927.11.1390 | CITFA-1 | 0-0-1-0 | nuclear enriched (mass spectrometry) | NRM | NRM | NRM |
| Q38DS7 | Tb927.9.10960 | DEAD/H RNA helicase | other | nuclear enriched (mass spectrometry) | 5SUP_B | DEAD-box RNA helicase | 100 |
| Q57XM9 | Tb927.7.2450 | uncharacterized | 0-0-1-0 | - | 3NFQ_B<br>6PWV_A | Spn1/Iws1;<br>Histone binding | 100<br>92.74 |

### Enriched in YFP-HDAC3 affinity selections

| UniProt ID | TriTryp ID | Gene name/description | [1]RIT-seq scoring matrix | Localization (BF) | PDB entry | HHpred domain | [2]HHpred domain probability % |
|---|---|---|---|---|---|---|---|
| Q387L4 | Tb927.11.370 | RAP1; also detected with CRD1 & TRF | 0-0-1-0 | nuclear enriched (mass spectrometry); telomeric | 3AL2_A | BRCT domain | 94.26 |
| Q57UB5 | Tb927.5.3510 | SMC protein | other | nuclear enriched (mass spectrometry) | 6WG3_B | SMC Cohesin/Condensin | 100 |
| Q580Q4 | Tb927.4.5310 | RDK2 | other | - | 6VPM_A | Serine/ThreonineProtein Kinase | 100 |
| Q388S9 | Tb927.10.13180 | NRAP | other | nuclear enriched (mass spectrometry) | 4M5D_A | small nucleolar RNA-associated protein | 100 |
| Q4GZ82 | Tb927.1.1370 | rRNA biogenesis protein | other | nuclear enriched (mass spectrometry) | 6LQU_RD | Ribosome assembly protein | 100 |
| Q580T1 | Tb927.3.5440 | SNF2 DNA repair protein | other | - | 6IGM_H | SWI/SNF family chromatin remodeler | 100 |
| Q38DK9 | Tb927.9.11850 | SMC protein | other | nuclear enriched (mass spectrometry) | 6WG3_A | SMC Cohesin/Condenin | 100 |
| Q385E2 | Tb927.11.6270 | PIP5Pase1; also detected with TRF | other | nuclear enriched (mass spectrometry) | 1I9Z_A | Phosphatidyl-Inositol phosphate 5-phosphatase | 99.89 |
| Q385P5 | Tb927.11.5230 | uncharacterized; also detected with BDF3, BDF5, HAT2 & ZCW1 | 0-0-0-0 | nuclear enriched (mass spectrometry); nuclear (D'Archivio and Wickstead, 2017) | 1UZ3_A | EMSY ENT domain (Chromo shadow binding | 95.93 |
| Q38DH5 | Tb927.9.12300 | RFC | 1-1-1-0 | nuclear enriched (mass spectrometry) | 6VVO_E | Replication factor C subunit 1 | |

### SET26-ZCW1 network

| UniProt ID | TriTryp ID | Gene name/description | [1]RIT-seq scoring matrix | Localization (BF) | PDB entry | HHpred domain | [2]HHpred domain probability % |
|---|---|---|---|---|---|---|---|
| Q38AF4 | Tb927.10.8100 | SET26 | 1-1-1-1 | nuclear/cytoplasmic (this study) | 3RQ4_A<br>1T6S_A | SET SUVAR20H2;<br>Winged-HTH | 99.54<br>66.7 |
| Q389G9 | Tb927.10.11720 | ZCW1;<br>also detected with BDF2 | other | nuclear (this study); nuclear enriched (mass spectrometry) | 6O5W_A | MORC family CW-type zinc finger | 97.53 |
| Q580R3 | Tb927.3.5620 | SPT16 (FACT) | 0-0-0-0 | nuclear enriched (mass spectrometry) | 6UPL_G | Histone Chaperone FACT Complex | 100 |
| Q385D4 | Tb927.11.6350 | AAA ATPase; BDF7; similar to Yta7 (*S. cerevisiae*), ATAD2 (*H. sapiens*) | 1-1-1-1 | nuclear enriched (mass spectrometry) | 6JQ0_C<br>6DNE_B | AAA+ ATPase Histone Chaperone;<br>Bromo (PFAM) | 99.95<br>98.53 |
| Q388G3 | Tb927.10.14390 | POB3 (FACT) | 0-0-0-0 | nuclear enriched (mass spectrometry) | 6UPL_G | Histone Chaperone FACT Complex | 100 |
| Q583S0 | Tb927.6.2640 | Importin alpha;<br>also detected with TFIIS2-2, DOT1A, NUP110 & TRF | 0-0-0-0 | nuclear enriched (mass spectrometry) | 4TNM_A | Importin Alpha 3 | 100 |
| Q383E9 | Tb927.11.11290 | HSP70;<br>also detected with BDF3, BDF5, BDF7, TFIIS2-2 & DOT1A | 1-1-1-0 | nuclear enriched (mass spectrometry) | 5FPN_A | Heast Shock 70 protein | 100 |
| Q386P3 | Tb927.11.2670 | NUP59 | 0-0-0-0 | nuclear enriched (mass spectrometry) | 5IJN_G | Nucleopore Complex (Nup155) | 99 |
| Q38D21 | Tb927.9.14430 | Casein Kinase II;<br>also detected with BDF4 & BDF7 | other | - | 5XVU_B | Caesin Kinase CKII | 100 |
| Q386Y4 | Tb927.11.1700 | uncharacterized | other | - | 2GIA_D | RNA binding | 97.54 |

### Enriched in YFP-SET26 affinity selections

| UniProt ID | TriTryp ID | Gene name/description | [1]RIT-seq scoring matrix | Localization (BF) | PDB entry | HHpred domain | [2]HHpred domain probability % |
|---|---|---|---|---|---|---|---|
| Q384B5 | Tb927.11.9130 | Replication factor A protein 1; possible SS telomeric DNA binding | 0-0-0-0 | - | 6D6V_D 1JB7_A | Telomere SS terminal DNA 3′ extension binding protein | 100 99.58 |
| Q583H5 | Tb927.4.3810 | RPB2 | 0-0-0-0 | nuclear enriched (mass spectrometry) | 6GMH_B | RNAPII Rpb2 | 100 |
| Q38D92 | Tb927.9.13320 | uncharacterized; also detected with BDF3, BDF5, & HAT2 | other | nuclear enriched (mass spectrometry) | 5XZW_A | FHA domain (binds phosphothreonine and phosphotyrosine peptides) | 96.13 |
| Q38AE0 | Tb927.10.8240 | Cytokinesis initiation factor 4 | 0-0-1-0 | - | 6JLB_D | Coiled-coil | 96.19 |
| Q580N2 | Tb927.3.4100 | Pyruvate transporter | other | - | 6G9X_B | Major facilitator superfamily transporter | 100 |
| Q38A77 | Tb927.10.8970 | kinetoplast-associated protein 4 isoform 2 | - | - | 3TQ6_A | DNA binding (mitochondrial) | 99.64 |
| Q38E19 | Tb927.9.9810 | uncharacterized | other | nuclear enriched (mass spectrometry) | NRM | NRM | NRM |
| Q57YU2 | Tb927.8.7400 | RPB1; also detected with CRD1 & TFIIS2-2 | 0-0-0-0 | nuclear enriched (mass spectrometry) | 4XPZ_A | RNAPII Rpb1 | 96.21 |
| Q385C3 | Tb927.11.6460 | uncharacterized | other | - | 4P5X_A | Mitochrondrial carrier protein | 99.77 |
| Q38FW0 | Tb927.9.1980 | uncharacterized | 1-1-1-0 | - | NRM | NRM | NRM |
| Q57WL5 | Tb927.8.8150 | C2 domain; also detected with BDF1 | 0-0-0-0 | - | 6FLJ_A | C2 domain (calcium binding, membrane bound) | 99.2 |
| Q57ZS7 | Tb927.5.2080 | GMP reductase; IMPDH | other | glycosomal (Guther et al., 2014) | 6RFU_A | Inosine-5'-monophosphate dehydrogenase | 100 |
| Q387Q3 | Tb927.10.16120 | IMPDH1; also detected with DOT1A & SET15 | 1-1-1-0 | glycosomal | 6RFU_A | Inosine-5'-monophosphate dehydrogenase | 100 |
| Q388C7 | Tb927.10.14770 | Associated kinase of Tb14-3-3 | 1-1-1-0 | - | 6VPM_A | Serine/Threonine protein kinase | 100 |
| Q388K3 | Tb927.10.13960 | paralyzed flagella protein 20 | 0-0-1-0 | flagellar | 5L8E_B | WD40 repeat protein | 100 |
| Q389A2 | Tb927.10.12410 | uncharacterized; also detected with BDF1 | other | - | 1G0S_A | ADP-ribose pyrophosphatase | 99.72 |
| Q38DT1 | Tb927.9.10920 | kinetochore protein KKT3 | 0-0-1-0 | nuclear enriched (mass spectrometry) | 6VPM_A | Serine/Threonine protein kinase | 100 |
| Q584R4 | Tb927.6.2420 | p22 protein precursor | other | - | 3JV1_A | P22, Mam33 family | 100 |

### Enriched in YFP-SET26 affinity selections - continued

| UniProt ID | TriTryp ID | Gene name/description | [1]RIT-seq scoring matrix | Localization (BF) | PDB entry | HHpred domain | [2]HHpred domain probability % |
|---|---|---|---|---|---|---|---|
| Q386D5 | Tb927.11.3770 | Dpy-30 motif containing protein | 0-0-1-0 | - | 4RT4_C<br>1XIQ_F | DPY-30 interaction domain (histone methyltransferase regulation);<br>Nucleoside diphosphate kinase | 98.73<br>99.85 |
| Q381T5 | Tb927.11.15000 | SMN-like | other | nuclear enriched (mass spectrometry) | 5XJL_M | SMN-related snRNA assembly | 87.4 |
| Q38B98 | Tb927.10.5030 | ubiquitin/ribosomal protein S27a | 0-0-1-0 | - | 5OPT_j | Kinetoplastid specific ribosomal protein (KSRP) | 100 |
| Q580S5 | Tb927.3.5500 | RPB3;<br>also detcted with CRD1 & TFIIS2-2 | other | nuclear enriched (mass spectrometry) | 1TWF_C | Rpb3 RNAPII subunit | 100 |
| Q582W2 | Tb927.3.2490 | Interacts with kinesin | 1-1-0-0 | - | 1V8K_A<br>4QJ3_B<br>2ELJ_A | Kinesin-related;<br>Nucleotide-binding;<br>Transcriptional adapter | 100<br>98.06<br>99.04 |
| Q57X47 | Tb927.5.4570 | Flagellum adhesion protein 3 | 0-0-1-0 | - | 6GC1_C | NHL repeat β-propeller | 99.21 |
| Q57YE9 | Tb927.8.820 | VID27 cytoplasmic protein | other | nuclear enriched (mass spectrometry) | 5ZWM_K<br>4J87_A | Splicing factor;<br>Vesicle trafficking | 99.88<br>99.90 |

### Enriched in YFP-ZCW1 affinity selections

| UniProt ID | TriTryp ID | Gene name/description | [1]RIT-seq scoring matrix | Localization (BF) | PDB entry | HHpred domain | [2]HHpred domain probability % |
|---|---|---|---|---|---|---|---|
| Q57UN4 | Tb927.7.4040 | Similar to Swc4 (*S. cerevisiae*); also detected with BDF2 | other | nuclear enriched (mass spectrometry) | 4IEJ_A | DMAP1, chromatin regulator, SWR1 complex | 97.52 |
| Q383K6 | Tb927.11.10730 | SWI/SNF-related helicase; similar to Swr1 (*S. cerevisiae*); also detected with BDF2 | other | nuclear enriched (mass spectrometry) | 6FML_G | ATPase Helicase, SWR1/INO80 complexes | 100 |
| Q580C6 | Tb927.4.980 | actin-related; similar to Act1 (*S. cerevisiae*); also detected with BDF2 | 1-1-1-1 | nuclear enriched (mass spectrometry) | 4AM6_A | Actin or Actin-related protein, Chromatin remodelling | 100 |
| Q389H2 | Tb927.10.11690 | YEATS; similar to YEA1; Yaf9 (*S. cerevisiae*); also detected with BDF2 | other | nuclear enriched (mass spectrometry) | 6T1I_A | YEATS domain Acetyl-/Crontonyl-lysine binding | 98.21 |
| Q57ZL0 | Tb927.3.3020 | actin-related; similar to Arp6 (*S. cerevisiae*); also detected with BDF2 | other | nuclear enriched (mass spectrometry) | 4FO0_A | Actin-related protein, Chromatin remodelling | 100 |
| Q584P9 | Tb927.6.2570 | actin-related; also detected with BDF2 | other | nuclear enriched (mass spectrometry) | 4I6M_B | Actin-related protein, Chromatin remodelling | 97.36 |
| Q38C43 | Tb927.10.2000 | actin-related; similar to Arp4 (*S. cerevisiae*); also detected with BDF2 | other | nuclear enriched (mass spectrometry) | 6GEN_R | Actin-related protein, Chromatin remodelling | 100 |
| Q385E0 | Tb927.11.6290 | HIT zinc finger; similar to Swc6 (*S. cerevisiae*); also detected with BDF2 | 1-1-1-1 | nuclear enriched (mass spectrometry) | 2N95_A<br>6GEJ_S | HIT Zinc Finger; SWR1/INO80 complex linked to RuvB | 97.12<br>99.4 |
| Q57YH0 | Tb927.8.600 | Bucentaur/craniofacial development; also detected with BDF2 | other | nuclear enriched (mass spectrometry) | 6PX3_S | Histone/nucleosome binding (possible *S. cerevisiae* equivalent SWR1-C Swc5) | 53.46 |
| Q583J3 | Tb927.4.2000 | ruvB-like DNA helicase; similar to Rvb1/2 (*S. cerevisiae*); also detected with BDF2 | 0-0-1-0 | - | 6H7X_A | RuvB-like ATPase helicase, SWR1/INO80 complexes | 99.96 |
| Q385I5 | Tb927.11.5830 | Similar to Swc2 (*S. cerevisiae*) | 0-0-0-0 | - | 5FUG_C | H2A.Z Chaperone, Nucleosome assembly, SWR1 complex | 99.51 |
| Q581V4 | Tb927.4.1270 | ruvB-like DNA helicase; similar to Rvb1/2 (*S. cerevisiae*); also detected with BDF2 | 0-0-1-0 | - | 2C9O_C | RuvB-like ATPase helicase, SWR1/INO80 complexes | 100 |
| Q387D1 | Tb927.11.1210 | DUF4470; DUF4471 | 1-1-1-0 | - | 5H02_A | Domain shared with methyltransferases | 83.83 |
| Q38AM1 | Tb927.10.7420 | BDF2; similar to Bdf1 (*S. cerevisiae*); also detected with HDAC3 | 1-1-1-1 | - | 6DNE_B<br>2ZD7_A | Bromo domain; NAP domain (histone chaperone) | 99.29<br>83.7 |
| Q385P5 | Tb927.11.5230 | uncharacterized; BDF3, BDF5, HAT2 & HDAC3 | 0-0-0-0 | nuclear (D'Archivio and Wickstead, 2017) nuclear enriched (mass spectrometry) | 1UZ3_A | EMSY ENT domain (Chromo shadow binding) | 95.93 |
| Q57XG4 | Tb927.8.2250 | tRNA ligase phosphodiesterase domain containing protein | other | - | 6N67_A<br>6TZO_A | tRNA ligase; nucleotide-binding | 98.27<br>99.96 |
| Q386Y7 | Tb927.11.1670 | Cysteine desulfurase; also detected with BDF7 | other | nuclear (Kovarova et al., 2014; Smid et al., 2006) | 4Q75_A | Cysteine Desulfurase | 100 |

### Enriched in YFP-ZCW1 affinity selections – continued

| UniProt ID | TriTryp ID | Gene name/description | [1]RIT-seq scoring matrix | Localization (BF) | PDB entry | HHpred domain | [2]HHpred domain probability % |
|---|---|---|---|---|---|---|---|
| Q4GYK7 | Tb927.1.3670 | ESAG8 protein | - | - | 3RGZ_A | Leucine Rich Repeat Receptor | 100 |
| Q382J7 | Tb927.11.13400 | BDF5; also detected with BDF3 & HAT2 | 0-0-1-0 | nuclear (this study); nuclear enriched (mass spectrometry) | 2R0Y_A<br>2F5J_B | Tandem Bromo Domains;<br>MRG Domain | 99.7<br>96.87 |
| Q387T9 | Tb927.10.15750 | TAC protein 197 | 0-0-0-0 | basal bodies (Gheiratmand et al., 2013; Halliday et al., 2019) | 6ZP4_A | 40S Ribosomal protein | 97.27 |
| Q57ZX9 | Tb927.5.3030 | Intraflagellar transport protein 121 | 0-0-1-0 | - | 5N4A_A | Intraflagellar transport WD40, ZnF_RBZ | 100 |

### BDF6-EAF6-HAT1 network

| UniProt ID | TriTryp ID | Gene name/description | [1]RIT-seq scoring matrix | Localization (BF) | PDB entry | HHpred domain | [2]HHpred domain probability % |
|---|---|---|---|---|---|---|---|
| Q4GYN3 | Tb927.1.3400 | BDF6 | other | nuclear (this study); nuclear enriched (mass spectrometry) | 2R10_B | Bromo domain | 99.14 |
| Q38FN2 | Tb927.9.2910 | EAF6 NuA4 HAT subunit; similar to Eaf6 (*S. cerevisiae*); also detected with PHD1 & HAT3 | 1-1-1-1 | nuclear (this study); nuclear enriched (mass spectrometry) | 5J9T_B | Eaf6 subunit of NuA4 histone acetyltransferase complex | 99.85 |
| D6XIY7 | Tb927.7.4560 | HAT1 | 1-1-1-1 | nuclear (this study) | 7CMR_A | MYST Histone acetyltransferase | 100 |
| Q57ZF8 | Tb927.7.5310 | YEATS; similar to YEA2; Yaf9 (*S. cerevisiae*) | other | nuclear enriched (mass spectrometry) | 5IQL_A | YEATS domain Acetyl-/Crotonyl-lysine histone binding | 99.82 |
| Q388I3 | Tb927.10.14190 | EPL1 NuA4 HAT subunit; similar to Epl1 (*S. cerevisiae*) | 1-1-1-1 | nuclear enriched (mass spectrometry) | 5J9T_K | Epl1 subunit of NuA4 histone acetyltransferase complex | 98.5 |
| Q4GZG2 | Tb927.1.650 | EAF3 NuA4 HAT subunit; similar to Eaf3 (*S. cerevisiae*) | 0-0-0-0 | nuclear enriched (mass spectrometry) | 2F5J_B | MRG domain protein; MRG15 | 97.87 |
| Q57UK4 | Tb927.8.5320 | uncharacterized | 1-1-1-1 | - | 2L9Z_A | Zinc binding | 46.06 |
| Q386G8 | Tb927.11.3430 | uncharacterized | other | - | NRM | NRM | NRM |
| Q585C3 | Tb927.6.1240 | uncharacterized | 0-0-0-0 | - | NRM | NRM | NRM |

### Enriched in YFP-BDF6 affinity selections

| UniProt ID | TriTryp ID | Gene name/description | [1]RIT-seq scoring matrix | Localization (BF) | PDB entry | HHpred domain | [2]HHpred domain probability % |
|---|---|---|---|---|---|---|---|
| Q57UX1 | Tb927.8.4810 | Prohibitin1 | other | - | 4FVG_A | SPFH (Stomatin, Prohibitin, Flotillin, HflK/C) domain | 99.71 |
| Q57Z72 | Tb927.5.3210 | SUMO;<br>also detected with BDF3, HAT2 & SET27 | other | - | 3PGE_A | SUMO (Small Ubiquitin-like protein) | 99.03 |

### Enriched in YFP-HAT1 affinity selections

| UniProt ID | TriTryp ID | Gene name/description | [1]RIT-seq scoring matrix | Localization (BF) | PDB entry | HHpred domain | [2]HHpred domain probability % |
|---|---|---|---|---|---|---|---|
| Q383W5 | Tb927.11.9640 | glycyl-tRNA synthetase | 0-0-1-0 | - | 2ZT5_A | Glycyl-tRNA synthetase | 100 |
| Q389D8 | Tb927.10.12040 | MAPK11 | other | - | 4IC7_A | Mitogen-activated protein kinase | 100 |

### Enriched in YFP-EAF6 affinity selections

| UniProt ID | TriTryp ID | Gene name/description | [1]RIT-seq scoring matrix | Localization (BF) | PDB entry | HHpred domain | [2]HHpred domain probability % |
|---|---|---|---|---|---|---|---|
| Q389Y3 | Tb927.10.9930 | PHD1 | 1-1-1-1 | nuclear (this study) | 4NN2_A<br>5J9T_K | PHD Zinc Finger domain (Nto1/NuA3);<br>Region similar to Epl1 | 99.45<br>97.0 |
| Q384N0 | Tb927.11.7880 | ING/NUA4 HAT;<br>similar to Yng2 (*S. cerevisiae*);<br>also detected with PHD1 & HAT3 | 0-0-1-0 | - | 5J9T_D | ING family (methyl-lysine histone), Esa1 NuA4 HAT complex | 96.76 |
| Q38AD3 | Tb927.10.8310 | HAT3 | 0-0-0-0 | nuclear (this study) | 3TO7_A | Esa1 MYST histone acetyltransferase | 100 |
| Q38C15 | Tb927.10.2290 | DnaJ3 | 1-1-0-0 | - | 3LZ8_B | DNAJ Chaperone | 100 |
| Q57V04 | Tb927.7.2070 | DNAJ | 1-1-1-0 | nuclear enriched (mass spectrometry) | 4J80_C | DNAJ Chaperone / Heat Shock protein 40 | 99.9 |
| Q38DC5 | Tb927.9.12900 | LEO1;<br>RNAPII associated;<br>also detected with TFIIS2-2 | 0-0-0-0 | - | 4M6T_A | Leo1 domain, Paf1 Complex - RNAPII transcription elongation regulation | 99.87 |
| Q586X9 | Tb927.2.3580 | TFIIS2-1;<br>also detected with CRD1, SET27 & TFIIS2-2 | 1-1-1-1 | nuclear enriched (mass spectrometry) | 1PQV_S<br>2M1H_A | TFIIS;<br>PWWP domain | 100<br>99.22 |
| D6XHJ3 | Tb927.6.1090 | proteasome regulatory ATPase subunit 3 | 0-0-1-0 | - | 6FVT_I | 26S Proteasome subunit | 99.96 |
| Q383D0 | Tb927.11.11480 | Trichohyalin (involved in stumpy formation) | other | - | NRM | NRM | NRM |
| Q57Y42 | Tb927.8.7980 | Pyrophosphate-energized vacuolar membrane proton pump 2 | 1-1-1-1 | - | 6AFW_B | Vacuolar membrane proton pump | 100 |

### Enriched in YFP-HDAC1 affinity selections

| UniProt ID | TriTryp ID | Gene name/description | [1]RIT-seq scoring matrix | Localization (BF) | PDB entry | HHpred domain | [2]HHpred domain probability % |
|---|---|---|---|---|---|---|---|
| Q38C74 | Tb927.10.1680 | HDAC1 | other | nuclear/cytoplasmic (this study); nuclear enriched (mass spectrometry) | 4A69_A | Histone Deacetylase | 100 |
| Q57WX2 | Tb927.3.890 | uncharacterized | 1-1-1-1 | - | 5JXT_L | ISWI Histone H4 tail bound | 35.78 |
| Q585H7 | Tb927.4.3730 | uncharacterized | other | - | 6R3E_B | PWWP Histone binding | 45.72 |
| Q584W0 | Tb927.6.3170 | uncharacterized | 1-1-1-1 | nuclear enriched (mass spectrometry) | 6IGM_H | RuvB-like, SWR/INO80 Complexes | 66.2 |
| Q57TZ5 | Tb927.7.1650 | uncharacterized | 1-1-1-1 | nuclear enriched (mass spectrometry) | NRM | NRM | NRM |
| Q38FV1 | Tb927.9.2070 | uncharacterized; also detected with CRD1 | other | nuclear enriched (mass spectrometry) | NRM | NRM | NRM |

### Enriched in YFP-BDF7 affinity selections (top interactors)

| UniProt ID | TriTryp ID | Gene name/description | [1]RIT-seq scoring matrix | Localization (BF) | PDB entry | HHpred domain | [2]HHpred domain probability % |
|---|---|---|---|---|---|---|---|
| Q385D4 | Tb927.11.6350 | AAA ATPase; BDF7; similar to Yta7 (*S. cerevisiae*), ATAD2 (*H. sapiens*); also detected with SET26 & ZCW1 | 1-1-1-1 | nuclear (this study); nuclear enriched (mass spectrometry) | 6JQ0_C<br>6DNE_B | AAA+ ATPase Histone Chaperone;<br>Bromo (PFAM) | 99.95<br>98.53 |
| Q4GZ00 | Tb927.1.2210 | NAP1; nucleosome assembly protein | other | - | 2AYU_A | Nucleosome Assembly Protein 1 histone chaperone | 100 |
| Q582P7 | Tb927.3.4880 | NAP2; nucleosome assembly protein | 1-1-1-1 | - | 5DAY_B | Nucleosome Assembly Protein 1-related histone chaperone | 100 |
| Q38D21 | Tb927.9.14430 | Casein kinase II; also detected with BDF4, SET26 & ZCW1 | other | - | 5XVU_B | Casein Kinase CKII | 100 |
| Q383J7 | Tb927.11.10820 | Casein kinase II subunit beta; also detected with BDF4 | other | - | 4DGL_A | Casein Kinase II subunit beta | 100 |
| Q386Y7 | Tb927.11.1670 | cysteine desulfurase; also detected with ZCW1 | other | nuclear (Kovarova et al., 2014; Smid et al., 2006) | 4Q75_A | Cysteine Desulfurase | 100 |
| Q585P0 | Tb927.6.1710 | uncharacterized | other | - | NRM | NRM | NRM |
| Q385C7 | Tb927.11.6420 | uncharacterized | other | nuclear enriched (mass spectrometry) | 5HDT_B | Cohesin regulator PDS5 | 100 |
| Q388S2 | Tb927.10.13250 | uncharacterized | 1-1-1-1 | - | 4J0W_A | Beta-Propeller, WD domain | 95.19 |
| Q388P7 | Tb927.10.13510 | zinc metallopeptidase | 1-1-1-1 | - | 6NYY_F | AAA+, ATPase, protease, mitochondria | 100 |
| Q381W2 | Tb927.11.14730 | Metalloprotease M41 FtsH | 1-1-1-0 | - | 6NYY_F | AAA+, ATPase, protease, mitochondria | 100 |
| Q389E2 | Tb927.10.12000 | iron-sulfur cluster assembly protein | other | - | 6UXE_B | Mitochondrial Cysteine desulfurase | 99.65 |
| Q38DH1 | Tb927.9.12340 | uncharacterized | 0-0-1-0 | nuclear enriched (mass spectrometry) | 4P5X_A | Calcium binding protein EF-Hand | 99.82 |
| Q387Z3 | Tb927.10.15180 | NAP3; nucleosome assembly protein | other | - | 2ZD7_A | NAP family histone Chaperone (Nap1/Vps75 S. cerevisiae) | 100 |
| Q383E9 | Tb927.11.11290 | HSP70; also detected with BDF3, BDF5, SET26, ZCW1, TFIIS2-2 & DOT1A | 1-1-1-0 | nuclear enriched (mass spectrometry) | 5FPN_A | Heast Shock 70 protein | 100 |
| Q38BN5 | Tb927.10.3640 | nuclear transmembrane protein | 1-1-1-0 | nuclear enriched (mass spectrometry) | NRM | NRM | NRM |
| Q57WG0 | Tb927.7.3450 | protein I/6 | - | - | 3MWU_A | Calcium binding protein EF-Hand | 99.59 |
| Q38BJ3 | Tb927.10.4070 | uncharacterized | other | nuclear enriched (mass spectrometry) | 2KMU_A | Possible ATP dependent helicase | 98.03 |

### Enriched in YFP-TFIIS2-2 affinity selections (top interactors)

| UniProt ID | TriTryp ID | Gene name/description | [1]RIT-seq scoring matrix | Localization (BF) | PDB entry | HHpred domain | [2]HHpred domain probability % |
|---|---|---|---|---|---|---|---|
| Q586Y0 | Tb927.2.3480 | TFIIS2-2 | other | nuclear (this study); nuclear enriched (mass spectrometry) | 3PMI_C 1PQV_S | PWWP domain; TFIIS | 99.04; 98.45 |
| Q582N0 | Tb927.3.5070 | uncharacterized | 0-0-1-0 | nuclear enriched (mass spectrometry) | NRM | NRM | NRM |
| Q38DC5 | Tb927.9.12900 | LEO1; RNAPII associated; also detected with EAF6 | 0-0-0-0 | - | 4M6T_A | Leo1 domain, Paf1 Complex - RNAPII transcription elongation regulation | 99.87 |
| Q57V64 | Tb927.3.3220 | CTR9; RNAPII associated | 0-0-0-0 | nuclear enriched (mass spectrometry) | 6AF0_A | Ctr9 / TPR Paf1 Complex - RNAPII transcription elongation regulation | 100 |
| Q586X9 | Tb927.2.3580 | TFIIS2-1; also detected with CRD1, EAF6 & SET27 | 1-1-1-1 | nuclear enriched (mass spectrometry) | 1PQV_S 2M1H_A | TFIIS; PWWP domain | 100 99.22 |
| Q57UN3 | Tb927.7.4030 | uncharacterized | 0-0-1-0 | nuclear enriched (mass spectrometry) | 4L1P_A | RTF1 subunit PAF Complex (RNAPII associated) | 96.9 |
| Q586C8 | Tb927.2.5810 | SPT6; similar to Spt6 (*S. cerevisiae*) | 0-0-0-0 | nuclear enriched (mass spectrometry) | 3PSI_A | Transcription elongation factor SPT6 | 100 |
| Q583H5 | Tb927.4.3810 | RPB2; also detected with SET26 | 0-0-0-0 | nuclear enriched (mass spectrometry) | 6GMH_B | RNAPII Rpb2 | 100 |
| Q57YU2 | Tb927.8.7400 | RPB1; also detected with CRD1 & SET26 | 0-0-0-0 | nuclear enriched (mass spectrometry) | 4XPZ_A | RNAPII Rpb1 | 96.21 |
| Q57Z69 | Tb927.5.3240 | uncharacterized; possible Rpb1-associated protein | 0-0-1-0 | nuclear enriched (mass spectrometry) | 3LLR_B 2C5Z_A 5IJO_N | PWWP domain; Set2 Rpb1-interacting (SRI) domain; Nuclear Pore Complex | 83.8 53.92 78.05 |
| Q38BC1 | Tb927.10.4800 | JBP3 | 1-1-1-1 | - | 2XSE_A | Thymine Dioxygenase | 98.34 |
| Q580S5 | Tb927.3.5500 | RPB3; also detcted with CRD1 & SET26 | | nuclear enriched (mass spectrometry) | 1TWF_C | Rpb3 RNAPII subunit | 100 |
| Q383S2 | Tb927.11.10070 | BDF3 | | nuclear enriched (mass spectrometry) | 4QB3_A 2NCZ_A | Bromo domain; ET Domain | 99.11 98.72 |
| Q389E6 | Tb927.10.11960 | PNUTS; also detected with SET27 | | - | NRM | NRM | NRM |

### Enriched in YFP-DOT1A affinity selections

| UniProt ID | TriTryp ID | Gene name/description | [1]RIT-seq scoring matrix | Localization (BF) | PDB entry | HHpred domain | [2]HHpred domain probability % |
|---|---|---|---|---|---|---|---|
| Q4GYF6 | Tb927.1.4220 | RNase H2B | 1-1-1-1 | nuclear enriched (mass spectrometry) | 3KIO_B | Ribonuclease H2 subunit B | 100 |
| Q581Z0 | Tb927.8.1920 | DOT1A | 1-1-1-1 | nuclear/cytoplasmic (this study) | 1U2Z_B | Histone lysine N-methyltransferase (H3K79-specific) | 99.75 |
| Q38B94 | Tb927.10.5070 | RNase H2A | other | - | 3P5J_A | Ribonuclease H2 subunit A | 100 |
| Q38F34 | Tb927.9.5190 | proliferating cell nuclear antigen (PCNA) | 0-0-0-0 | - | 5H0T_A | Proliferating cell nuclear antigen | 100 |
| Q4GYA5 | Tb927.1.4730 | RNase H2C | other | - | 3PUF_L | Ribonuclease H2 subunit C | 99.97 |
| Q583S0 | Tb927.6.2640 | importin alpha subunit; also detected with SET26, ZCW1, TFIIS2-2, NUP110 & TRF | 0-0-0-0 | nuclear enriched (mass spectrometry) | 4TNM_A | Importin Alpha 3 | 100 |
| Q387Q3 | Tb927.10.16120 | IMPDH1; also detected with SET26 & SET15 | 1-1-1-0 | glycosomal | 6RFU_A | Inosine-5'-monophosphate dehydrogenase | 100 |
| Q383E9 | Tb927.11.11290 | HSP70; also detected with BDF3, BDF5, BDF7, SET26, ZCW1 & TFIIS2-2 | 1-1-1-0 | nuclear enriched (mass spectrometry) | 2FE0_A | Heat Shock 70 protein | 100 |

### Enriched in YFP-TRF affinity selections (top interactors)

| UniProt ID | TriTryp ID | Gene name/description | [1]RIT-seq scoring matrix | Localization (BF) | PDB entry | HHpred domain | [2]HHpred domain probability % |
|---|---|---|---|---|---|---|---|
| Q388W1 | Tb927.10.12850 | TRF; telomere-repeat bining protein; also detected with BDF2 & HDAC3 | 0-0-0-0 | nuclear enriched (mass spectrometry), telomeric (Jehi et al, 2014; Li et al, 2005) | 1GV2_A | MYB domain DNA binding | 80.11 |
| Q586Z9 | Tb927.6.4330 | telomere-associated protein; also detected with BDF2 & HDAC3 | 0-0-0-0 | nuclear enriched (mass spectrometry), telomeric | NRM | NRM | NRM |
| Q38FD7 | Tb927.9.4000 | uncharacterized; also detected with BDF2 & HDAC3 | - | nuclear enriched (mass spectrometry), telomeric | NRM | NRM | NRM |
| Q57XY9 | Tb927.3.1560 | TIF2; also detected with BDF2 & HDAC3 | other | nuclear enriched (mass spectrometry), telomeric (Jehi et al, 2014) | NRM | NRM | NRM |
| Q383U2 | Tb927.11.9870 | TelAP1; also detected with BDF2 & HDAC3 | 1-1-1-1 | nuclear enriched (mass spectrometry), telomeric | 4B0Z_A | Proteasome subunit | 56.61 |
| Q385L3 | Tb927.11.5550 | DNA polymerase theta (POLQ); also detected with BDF2 & HDAC3 | 1-1-1-1 | nuclear enriched (mass spectrometry), telomeric | 4X0Q_B | DNA polymerase theta | 100 |
| Q38BZ3 | Tb927.10.2520 | PrimPol-like protein 2; also detected with BDF2 & HDAC3 | - | nuclear enriched (mass spectrometry) | 5L2X_B | DNA Primase Polymerase | 100 |
| Q387L4 | Tb927.11.370 | RAP1; also detected with CRD1 & HDAC3 | 0-0-1-0 | nuclear enriched (mass spectrometry), telomeric | 3AL2_A | BRCT domain | 94.26 |
| Q586J9 | Tb927.2.2190 | HDAC3; also detected with BDF2 | 0-0-0-0 | nuclear (Wang et al, 2010) | 2VQM_A | HDAC Histone Deacetylase | 100 |
| Q385E2 | Tb927.11.6270 | PIP5Pase1; also detetced with HDAC3 | - | nuclear enriched (mass spectrometry) | 1I9Z_A | Phosphatidyl-Inositol phosphate 5-phosphatase | 99.89 |

### Enriched in YFP-TBP affinity selections

| UniProt ID | TriTryp ID | Gene name/description | [1]RIT-seq scoring matrix | Localization (BF) | PDB entry | HHpred domain | [2]HHpred domain probability % |
|---|---|---|---|---|---|---|---|
| Q387R9 | Tb927.10.15950 | *TBP/TRF4 | other | nuclear (this study); nuclear enriched (mass spectrometry) | 1RM1_A | TATA-box binding protein | 100 |
| Q38E05 | Tb927.9.9970 | *SNAP50 | 1-1-1-1 | nuclear enriched (mass spectrometry) | 5N7Y_A | Zn Finger | 47.94 |
| Q599N4 | Tb927.10.15570 | *TFIIA-1 | other | - | NRM | NRM | NRM |
| Q599N5 | Tb927.10.7070 | *SNAP3 | other | nuclear enriched (mass spectrometry) | NRM | NRM | NRM |
| Q599N6 | Tb927.5.3910 | *SNAP2 | 1-1-0-0 | nuclear enriched (mass spectrometry) | 1FEX_A | Myb HTH DNA binding domain | 88.16 |
| Q387K4 | Tb927.11.470 | BRF1 (TFIIB) | - | - | 6F40_V<br>4BBR_M | TFIIIB;<br>TFIIB | 100<br>99.98 |
| Q599N3 | Tb927.10.4840 | *TFIIA-2 | 1-1-1-1 | - | 5M4S_A | TFIIA | 99.94 |
| Q38DC3 | Tb927.9.12950 | uncharacterized | other | - | NRM | NRM | NRM |
| Q384L9 | Tb927.11.8000 | uncharacterized | other | - | | | |

*TRF4/SNAPc/TFIIA Complex (Das et al, 2005; Schimanski et al, 2005)

### Enriched in YFP-ELP3b affinity selections

| UniProt ID | TriTryp ID | Gene name/description | [1]RIT-seq scoring matrix | Localization (BF) | PDB entry | HHpred domain | [2]HHpred domain probability % |
|---|---|---|---|---|---|---|---|
| Q57YY2 | Tb927.8.3310 | ELP3b | other | nuclear (this study); nuclear enriched (mass spectrometry) | 6IA8_A | Elp3 tRNA acetyltransferase | 100 |
| Q585H2 | Tb927.2.1210 | retrotransposon hot spot protein 4 (RHS4) | - | - | NRM | NRM | NRM |
| Q57ZH2 | Tb927.7.5170 | 60S ribosomal protein L25 | 0-0-0-0 | - | 4V8M_BX | Ribosome component | 100 |
| Q583F5 | Tb927.4.4700 | uncharacterized | 1-1-1-1 | - | 5IJO_N | Nuclear pore | 90.41 |
| Q387A9 | Tb927.11.1430 | Component of motile flagella 2 | 1-1-1-1 | flagellar | 6U42_6W | Tubulin related | 100 |

### Enriched in YFP-PHD2 affinity selections

| UniProt ID | TriTryp ID | Gene name/description | [1]RIT-seq scoring matrix | Localization (BF) | PDB entry | HHpred domain | [2]HHpred domain probability % |
|---|---|---|---|---|---|---|---|
| Q387D3 | Tb11.v5.0388 | PHD2 | other | nuclear (this study) | 6GLC_A | RBR-type ubiquitin ligase (Parkin) | 97.46 |

### Enriched in YFP-PHD4 affinity selections

| UniProt ID | TriTryp ID | Gene name/description | [1]RIT-seq scoring matrix | Localization (BF) | PDB entry | HHpred domain | [2]HHpred domain probability % |
|---|---|---|---|---|---|---|---|
| Q57Z97 | Tb927.3.2140 | PHD4 | 1-1-1-0 | nuclear (this study) | 3MWY_W<br>5XFO_A | Swi2/Snf2 ATPase remodeller (Chd1);<br>Tandem PHD Fingers | 100<br>97.54 |
| Q587E7 | Tb927.2.3370 | UDP-Gal or UDP-GlcNAc-dependent glycosyltransferase | other | - | 2J0A_A | Beta1,3-N-acetyl-glucosaminyl-transferase | 99.87 |

### Enriched in YFP-PHD1 affinity selections

| UniProt ID | TriTryp ID | Gene name/description | [1]RIT-seq scoring matrix | Localization (BF) | PDB entry | HHpred domain | [2]HHpred domain probability % |
|---|---|---|---|---|---|---|---|
| Q389Y3 | Tb927.10.9930 | PHD1;<br>also detected with EAF6 & HAT3 | 1-1-1-1 | nuclear (this study) | 4NN2_A<br>5J9T_K | PHD Zinc Finger domain (Nto1/NuA3);<br>Region similar to Epl1 | 99.45<br>97.0 |
| Q384N0 | Tb927.11.7880 | ING/NUA4 HAT; similar to Yng2 (*S. cerevisiae*);<br>also detected with EAF6 & HAT3 | 0-0-1-0 | - | 5J9T_D | ING family (methyl-lysine histone),<br>Esa1 NuA4 HAT complex | 96.76 |
| Q38FN2 | Tb927.9.2910 | EAF6 NUA4 HAT subunit; similar to Eaf6 (*S. cerevisiae*);<br>also detected with BDF6, HAT1 & HAT3 | 1-1-1-1 | nuclear (this study); nuclear enriched (mass spectrometry) | 5J9T_B | Eaf6 subunit of NuA4 histone acetyltransferase complex | 99.85 |
| Q38AD3 | Tb927.10.8310 | HAT3;<br>also detected with EAF6 | 0-0-0-0 | nuclear (this study) | 3TO7_A | Esa1 MYST histone acetyltransferase | 100 |

### Enriched in YFP-HAT3 affinity selections

| UniProt ID | TriTryp ID | Gene name/description | [1]RIT-seq scoring matrix | Localization (BF) | PDB entry | HHpred domain | [2]HHpred domain probability % |
|---|---|---|---|---|---|---|---|
| Q38AD3 | Tb927.10.8310 | HAT3;<br>also detected with EAF6 and PHD1 | 0-0-0-0 | nuclear (this study) | 3TO7_A | Esa1 MYST histone acetyltransferase | 100 |
| Q389Y3 | Tb927.10.9930 | PHD1;<br>also detected with EAF6 | 1-1-1-1 | nuclear (this study) | 4NN2_A<br>5J9T_K | PHD Zinc Finger domain (Nto1/NuA3);<br>Region similar to Epl1 | 99.45<br>97.0 |
| Q384N0 | Tb927.11.7880 | ING/NUA4 HAT; similar to Yng2 (*S. cerevisiae*);<br>also detected with EAF6 and PHD1 | 0-0-1-0 | - | 5J9T_D | ING family (methyl-lysine histone)<br>Esa1 NuA4 HAT complex | 96.76 |
| Q38FN2 | Tb927.9.2910 | EAF6 NUA4 HAT subunit; similar to Eaf6 (*S. cerevisiae*);<br>also detected with BDF6, HAT1, & PHD1 | 1-1-1-1 | nuclear (this study)<br>nuclear enriched (mass spectrometry) | 5J9T_B | Eaf6 subunit of NuA4 histone acetyltransferase complex | 99.85 |
| Q57YG1 | Tb927.8.690 | PIN1 | 1-1-1-1 | - | 2LJ4_A | Peptidyl-prolyl cis-trans isomerase | 99.68 |
| B3GVR6 | Tb427.BES126.2 | Transferrin-binding | - | - | 6SOY_B | ESAG6, subunit of heterodimeric transferrin | 100 |

### Enriched in YFP-AGO1 affinity selections

| UniProt ID | TriTryp ID | Gene name/description | [1]RIT-seq scoring matrix | Localization (BF) | PDB entry | HHpred domain | [2]HHpred domain probability % |
|---|---|---|---|---|---|---|---|
| Q389P5 | Tb927.10.10850 | AGO1 | 1-1-1-1 | cytoplasmic (this study) | 4Z4D_A | Argonaute protein | 100 |
| Q38A86 | Tb927.10.8880 | RIF4 | 1-1-1-1 | - | 4YBG_A | Ribonuclease, Maelstrom, silencing | 99.95 |
| Q389B0 | Tb927.10.12330 | ZC3H34 | 1-1-0-0 | - | 6EOJ_B<br>5GUH_A | 3' end processing Zinc finger;<br>Endonuclease | 92.87<br>65.86 |
| Q38E63 | Tb927.9.9290 | PABP1 | 0-0-0-0 | cytoplasmic (Kramer et al, 2013) | 6R5K_H | Polyadenylate-binding protein (Pab1) | 100 |
| Q585M4 | Tb927.6.1870 | EIF4E4 | 1-1-1-0 | cytoplasmic (Kramer et al, 2013) | 5T46_A | Eukaryotic translation initiation factor 4E | 100 |

### Enriched in YFP-NUP110 affinity selections

| UniProt ID | TriTryp ID | Gene name/description | [1]RIT-seq scoring matrix | Localization (BF) | PDB entry | HHpred domain | [2]HHpred domain probability % |
|---|---|---|---|---|---|---|---|
| Q387L8 | Tb927.11.330 | NUP110 (MLP1) | 0-0-1-0 | nuclear pores (this study); nuclear enriched (mass spectrometry) | 6FSA_A | Myosin light chain | 95.16 |
| | | 24 nucleoporins (see Table S5) | | | | | |

### Frequent non-specific contaminants

| UniProt ID | TriTryp ID | Gene name/description | [1]RIT-seq scoring matrix | Localization (BF) | PDB entry | HHpred domain | [2]HHpred domain probability % |
|---|---|---|---|---|---|---|---|
| Q586Q9 | Tb927.2.4580 | UNC119 | 1-1-1-1 | lipidated protein intraflagellar transport (Fritz et al., 2015; Pandey et al., 2020) | 3GQQ_E | human retinal protein 4 (unc-119) | 100 |

**Table S7. *T. brucei* homologs of SWR1-C/SRCAP-C/EP400 subunits**

| ***S. cerevisiae*** **SWR1-C** | ***H. sapiens*** **SRCAP-C or EP400** | ***T. brucei*** **SWR1-C** | **Description** |
|---|---|---|---|
| Swr1 | SRCAP | Tb927.11.10730 | Swi2/Snf2-related ATPase motor |
| Rvb1 | RuvBL1 | Tb927.4.2000/ Tb927.4.1270 | ATP-dependent DNA helicase |
| Rvb2 | RuvBL2 | Tb927.4.2000/ Tb927.4.1270 | ATP-dependent DNA helicase |
| Actin (Act1) | Actl6a/Arp4 | Tb927.4.980 | Actin |
| Arp4 | Cfdp1/Swc5 | Tb927.10.2000 | Actin-related |
| Arp6 | Arp6/Actr6 | Tb927.3.3020 | Actin-related |
| Swc2 | YL1 | Tb927.11.5830 | H2A.Z Chaperone |
| Swc3 | - | - | |
| Swc4 | DMAP1 | Tb927.7.4040 | |
| Swc6 | ZNHIT1 | Tb927.11. 6290 | |
| Swc7 | - | - | |
| Yaf9 | Gas41 | Tb927.10.11690 (YEA1) | YEATS domain; binds acetyl/crotonyl-lysine |
| Bdf1 | BRD8 | Tb927.10.7420 (BDF2) | Bromodomain protein |

**Table S8. *T. brucei* homologs of NuA4 subunits**

| ***S. cerevisiae*** **NuA4 Complex** | ***H. sapiens*** **NuA4** | ***T. brucei*** **NuA4 Complex** | **Description** |
|---|---|---|---|
| Eaf1 | [EP400] | - | |
| - | BRD8 | Tb927.1.3400 (BDF6) | Bromodomain |
| Epl1 | EPC1/EPC-like | Tb927.10.14190 | EPL1 domain |
| Esa1 | TIP60 | Tb927.7.4560 (HAT1)<br>Tb927.10.8310 (HAT3) | MYST Histone acetyltransferase |
| Eaf3 | MRG15 | Tb927.1.650 | MRG domain |
| Eaf6 | EAF6 | Tb927.9.2910 (EAF6) | NuA4 domain |
| Yng2 | ING3 | Tb927.11.7880 (YNG2) | ING family |
| Yaf9 | GAS41 | Tb927.7.5310 (YEA2) | YEATS domain binds acetyl/crotonyl-lysine |
